# Natural Killer Cell Receptor Signaling and Activation Depend on Cell Cycle Stages

**DOI:** 10.1101/2025.02.24.639882

**Authors:** Darren Wethington, Indrani Nayak, Helle Jensen, William C. Stewart, Oscar A. Aguilar, Shih-Yu Chen, Garry P. Nolan, Gregory K. Behbehani, Lewis L. Lanier, Jayajit Das

## Abstract

Receptor signaling in Natural Killer (NK) cells leads to post-translational modification (e.g., phosphorylation) of sub-cellular signaling proteins within minutes of receptor stimulation that eventually give rise to diverse effector functions including cell proliferation. Recent single-cell mass cytometry (i.e., CyTOF) experiments in macrophages showed variations of abundances of phosphorylated signaling proteins across cell cycle states indicating a dependence of cell signaling kinetics on an order of magnitude slower kinetics (~ several hours) of cell cycle transitions. We investigated cell cycle dependence of NKG2D signaling kinetics in NK cells by CyTOF measurements performed on IL-2-treated NKG2D-stimulated primary human CD56^dim^ NK cells. The CyTOF experiments revealed monotonic or semi-monotonic increases of the average protein abundances of the majority of signaling proteins such as pCrkL, pPLCγ2, and pErk, and the degranulation marker protein CD107a with progressing cell cycle states at specific time points post-NKG2D stimulation; however, several proteins such as pVav1, pS6, and pAkt, and early activation marker protein CD69 also showed non-monotonic variations in the average abundances with progressing cell cycle states. We used minimal mathematical and computational models coupling signaling and cell cycle processes to show that non-monotonic variations in the signaling protein abundances with progressing cell cycle stages are likely to arise in situations where protein synthesis and degradation and signaling kinetics are actively regulated by cell cycle processes.

## Introduction

Natural Killer (NK) cells are lymphocytes of our innate immune system and provide important protection against infection and tumors^1-3^. NK cells express a wide array of plasma membrane-bound activating and inhibitory NK cell receptors (NKRs), co-receptors, and adhesion receptors that interact with cognate ligands expressed on target cells^2^. In addition, NK cells express cytokine receptors such as IL-2 receptors that bind to soluble cytokine molecules present in the microenvironment^3^. Subcellular biochemical signaling reactions occurring in the time scales of seconds to minutes^4-6^, which are initiated by binding of diverse NKRs, co-receptors, and adhesion, and cytokine receptors under appropriate conditions may lead to NK cell activation^1,2^. Activated NK cells unleash a variety of effector functions ranging from cell proliferation, lysis of target cells, or secretion of cytokines such as interferon γ. NK cell division can take orders of magnitude longer time scales (~hours)^7,8^ to fully manifest compared to the signaling biochemical reactions. A proliferating eukaryotic cell spends most of the time during cell division in the cell cycle interphase where DNA, proteins, and metabolites are synthesized in the cell in preparation for the cell duplication^9^. The interphase is further subdivided into G1, S, and G2 stages that progress through the cell cycle as G1 → S → G2. At the end of the G2 stage, the cell goes through mitosis, M-stage, where the cell division takes place, and the resulting pair of daughter cells can exit the cell cycle by going into a quiescent stage G0 or continue dividing by entering the G1 stage^9^.

NKG2D is an activating receptor that transmits signals by its association with the DAP10 adapter protein through charge interactions between the transmembrane segments of NKG2D and DAP10^10^. The cytoplasmic domain of DAP10 possesses a Src homology-2 domain that upon tyrosine phosphorylation recruits and activates the p85 subunit of PI3 kinase and a Grb2-Vav1 complex that triggers NK cell activation and cell-mediated cytotoxicity^10,11^. The human NKG2D receptor binds a family of stress-induced ligands (MICA, MICB, and ULBP1-6) that are frequently displayed on the surface of tumor or pathogen-infected cells and render these cells susceptible to NK cell-mediated killing^12,13^. NK cells constitutively express an intermediate-affinity IL-2 receptor composed of IL-2Rγ and IL-2Rβ subunits that upon binding to IL-2 or IL-15 transmit activation and growth signals through the JAK – STAT pathway that initiates cellular proliferation ^14^. Therapeutics targeting NKG2D and IL-2 are currently under clinical investigation in cancer patients.

NK cells undergoing proliferation can reside in different cell cycle stages as they interact with target cells with plasma membrane-bound receptors. As single cells progress through cell cycle transitions many signaling proteins such as kinases and phosphatases, as well as their substrate proteins, are synthesized to maintain a steady per cell copy number or abundance of these proteins during cell growth^15,16^. Thus, as single NK cells progress through cell cycle stages the abundances of reacting signaling proteins could change resulting in different abundances of signaling products. For example, the abundance of a phosphorylated form of a signaling protein (e.g., pVav1) could be different in NK cells residing in different cell cycle stages as the abundances of phosphorylating or dephosphorylating enzymes, along with the abundance of the substrate protein (e.g., Vav1) vary across cell cycle stages. In addition, proteins regulating cell cycle processes, such as cyclin dependent kinases (CDKs), can actively participate in chemically modifying signaling proteins^17,18^. For example, in HeLa cells the phosphorylation of the site S473 in signaling protein Akt is regulated by CDK2^19^. This produces variations in the abundances of the phosphorylated protein Akt-pS473 similar to that of CDK2 abundances across cell cycle states while the total Akt abundance remains unchanged^19^. In another example, CDK proteins in G2-M state phosphorylate Wnt-binding co-receptor LRP6 promoting Wnt signaling in the state^18^.

Single-cell mass cytometry (CyTOF) experiments can simultaneously measure expressions of proteins that can track cell cycle progression and receptor signaling kinetics^20-23^. CyTOF experiments performed on human monocyte cell line THP-1 stimulated by TNFα showed that average abundances of modified forms of signaling proteins can depend on cell cycle stages^24^. Here we investigated cell cycle dependence on NK cell signaling and activation by performing CyTOF measurements on IL-2-treated primary human CD56^dim^ NK cells stimulated by plate-bound anti-NKG2D antibodies at different time points. A total of 40 different proteins were evaluated by CyTOF, mostly pertaining to signaling proteins, activation markers, and cell cycle markers. While the average abundance of the majority of the signaling proteins evaluated showed monotonic or semi-monotonic increases with the progression of cell cycle stages, the average abundances of some proteins such as pAkt, pMAPKAPK2, pPLK1, pS6, pNFκB, and pVav1 also showed non-monotonic variations at specific time points post-NKG2D stimulation. The potential mechanisms that underlie these variations of abundances of signaling products across cell cycle stages can range from changes in the abundances of reacting signaling proteins in different cell cycle stages to active participation of CDK proteins in cell signaling. We developed minimal mathematical and computational models that couple cell cycle transitions and signaling reactions to quantify influences of cell cycle transitions in producing variations in the abundances of signaling proteins across cell cycle states. We find that monotonic increases in the average abundances of phosphorylated proteins with progressing cell cycle states can be produced by protein synthesis/degradation and phosphorylation/dephosphorylation rates that do not change with cell cycle stages. However, the occurrence of non-monotonic changes or decreases in the average abundances of phosphorylated proteins with progressing cell cycle stages calls for specific conditions such as limitation of enzyme concentrations, or direct intervention of cell cycle regulating proteins in phosphorylation/dephosphorylation reactions. Thus, the nature of variation of the abundances of signaling proteins with cell cycle stages can point to different mechanisms of regulation of receptor signaling kinetics by cell cycle processes.

## Results

### Phosphoprotein abundances show monotonic and non-monotonic changes with progressing cell cycle stages in NKG2D-stimulated, IL-2-treated primary NK cells

The abundances of proteins associated with NKG2D signaling can potentially vary across cell cycle states in NK cells stimulated with NKG2D ligands due to the differences in the abundances of reacting proteins in cells residing in different cell cycle stages or the regulation of the signaling kinetics by proteins (e.g., CDKs) involved in driving cell cycle progression (**Figure 1**). We investigated the variation of NKG2D signaling proteins in primary human NK cells. Primary human CD56^dim^ NK cells were treated with IL-2 for 24 hours to prime the NKG2D signaling pathway and then stimulated with plate-bound anti-NKG2D agonist antibodies. NKG2D transmits signals through the DAP10 adaptor protein, which recruits and activates the phosphatidylinositol 3-kinase^10^. The NK cells were profiled at 2, 4, 8, 16, 32, 64, and 256 minutes post-NKG2D stimulation using CyTOF^25^, which assayed 31 different signaling proteins and surface markers simultaneously (**Figure 2A**). The various cell cycle stages (i.e., G0, G1, S, and G2 + M) were gated using UMAP based on the cell cycle markers, 5-iodo-2-deoxyuridine (IdU), cyclin B1, phosphorylated histone 3 (pH3), phosphorylated retinoblastoma protein (pRb) and Kiel 67 (Ki67) (**Figure 2B, Supplementary Figure S1**). Further details regarding the analysis are provided in Materials and Methods. ~10-20% of the NK cells were in G0, ~60-70% were in G1, and ~10% were in S and G2 + M stages, respectively, throughout the various time points post-NKG2D stimulation (see **Figure 2C** and **Figure 2D**). The variations of the average protein abundances across cell cycle stages G1, S, and G2 + M post-NKG2D stimulation were analyzed as follows. We classified the variations into nine different types (see labels in **Figure 2E**) based on the change, namely, rise, or decay, or no-change, in the average abundance between G1 to S, and S to G2+M for each time point by performing one-sided *t*-tests. Because the protein distributions follow non-normal distributions, we natural log-transformed the protein intensities before performing the *t*-test comparing average abundances of proteins residing in different cell cycle states (**Figure 2E**). More details are provided in Methods. At *t* = 0 min (i.e., unstimulated state of the IL-2-treated CD56^dim^ NK cells), 8 proteins that were not related to cell cycle displayed significant increases from G1 to S and S to G2-M, and the majority of the proteins (22 out of 31 showed no change between G1 to S followed by a rise from S to G2 + M stage (**Figure 2E**). Next, we investigated the variation of the average abundances with cell cycle stages at multiple time points post-NKG2D stimulation. The majority (~ 89%) of the proteins showed a strictly monotonic increase (i.e., increase for G1 → S and increase for S → G2 + M) or a semi-monotonic increase (i.e., non-change for G1 → S and increase for S → G2, or vice versa) with progressing cell cycle stages between 2 to 256 mins signaling kinetics. Proteins involved in cell signaling that showed monotonic or semi-monotonic behavior post-NKG2D stimulation included pCrkL, pPLCγ2, pAkt, pMAPKAPK2, pSLP76, pNFκB, pErk1-2, pPLK1, p38, pLAT, pZAP70, pS6, and pVav1. Interestingly, 12 proteins (~ 9%) including pAKT, pMAPKAPK2, pNFκB, pPLK1, pS6, and pVav1 also showed non-monotonic behavior in protein abundances at specific time points post-NKG2D stimulation marked by a decrease for G1→ S followed by an increase from S to G2 + M (**Figure 2E**). We did not find any protein showing a monotonic decrease, i.e., decrease for G1→ S and decrease for S → G2 + M. The early cell activation marker, CD69, showed non-monotonic behavior at 32, 64, and 256 min post-NKG2D stimulation (**Figure 2E-2G**). CD107a is a marker of NK cell activity, indicating degranulation and cytotoxic function when present on the cell surface. The average CD107a protein abundances exhibited monotonic or semi-monotonic behavior at all time points with the highest levels observed at 256 min post-NKG2D stimulation. These highest abundances displayed semi-monotonic behavior with progressing cell cycle states (**Figures 2E** and **2G, Supplementary Figure S2**). In summary, in NKG2D receptor-stimulated primary NK cells the average abundances of most of the measured signaling proteins, as well as the two activation markers, followed a strictly monotonic or semi-monotonic increase with progressing cell cycle states for most time points, with certain proteins such as pAkt, pMAPKAPK2, pNFκB, pPLK1, pS6, and pVav1 showing non-monotonic changes at specific time points post-NKG2D stimulation (**Supplementary Figure S2**).

**Figure 1:**
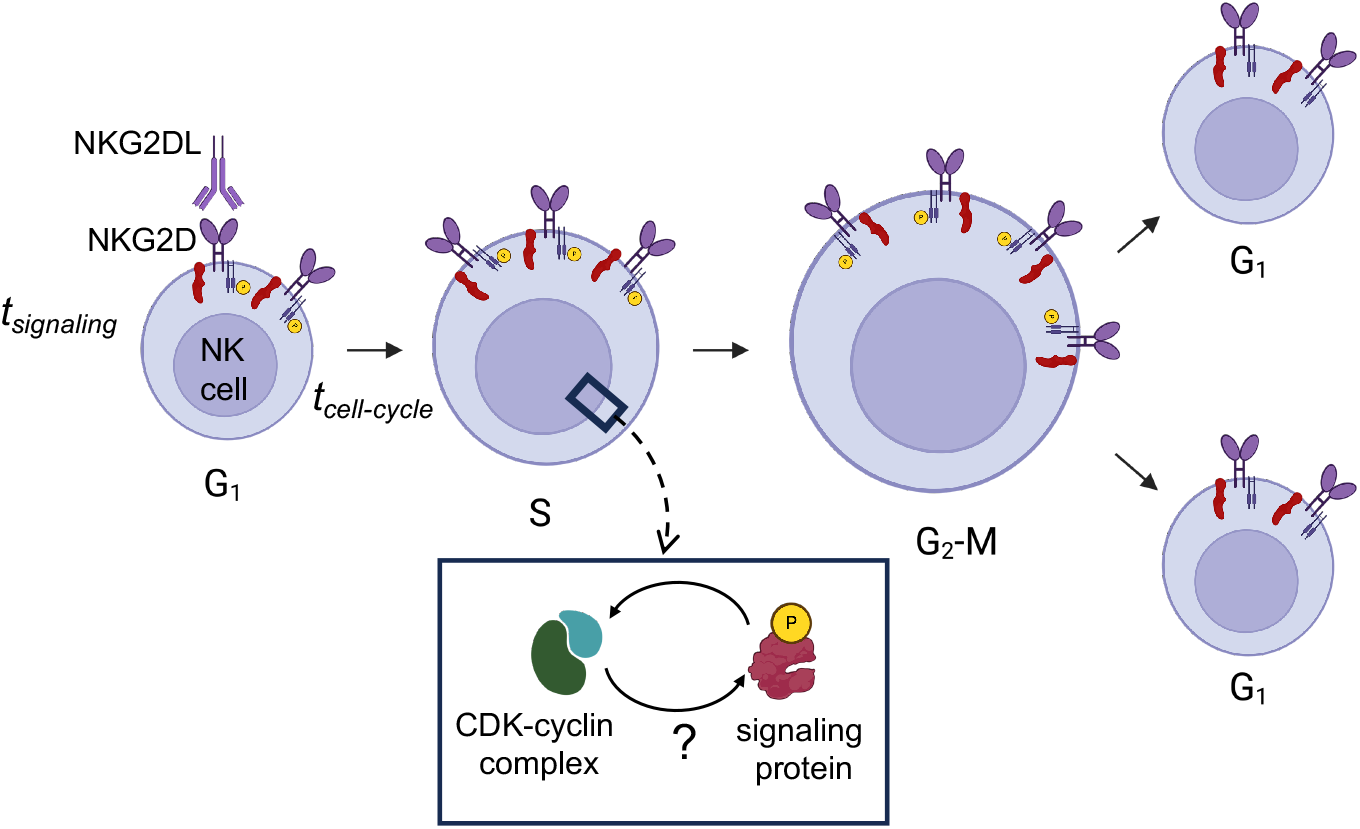
Potential influences of cell cycle transitions on NKG2D receptor signaling kinetics in NK cells. Schematic depiction of potential mechanisms with which cell cycle transitions can regulate NKG2D receptor signaling. As an NK cell progresses through cell cycle stages, the abundances of many proteins including ones involved in NKG2D signaling can increase. The time scale (*t*_*signaling*_) for cell signaling is typically in seconds to minutes and can be 10-100× smaller than the timescales (*t*_*cell cycle*_) of cell cycle transitions which can take hours to occur. The cell volume increases with cell cycle progression as well. The combined effect of increasing protein abundances and the cell volume can increase or decrease the frequency at which second-order biochemical signaling reactions occur. This can result in different abundances of signaling products arising from second order signaling reactions in different cell cycle stages. In addition, cell cycle regulating CDK-cyclin complexes can potentially influence post-translational modifications of signaling proteins in specific cell cycle stages affecting NKG2D signaling kinetics.

**Figure 2:**
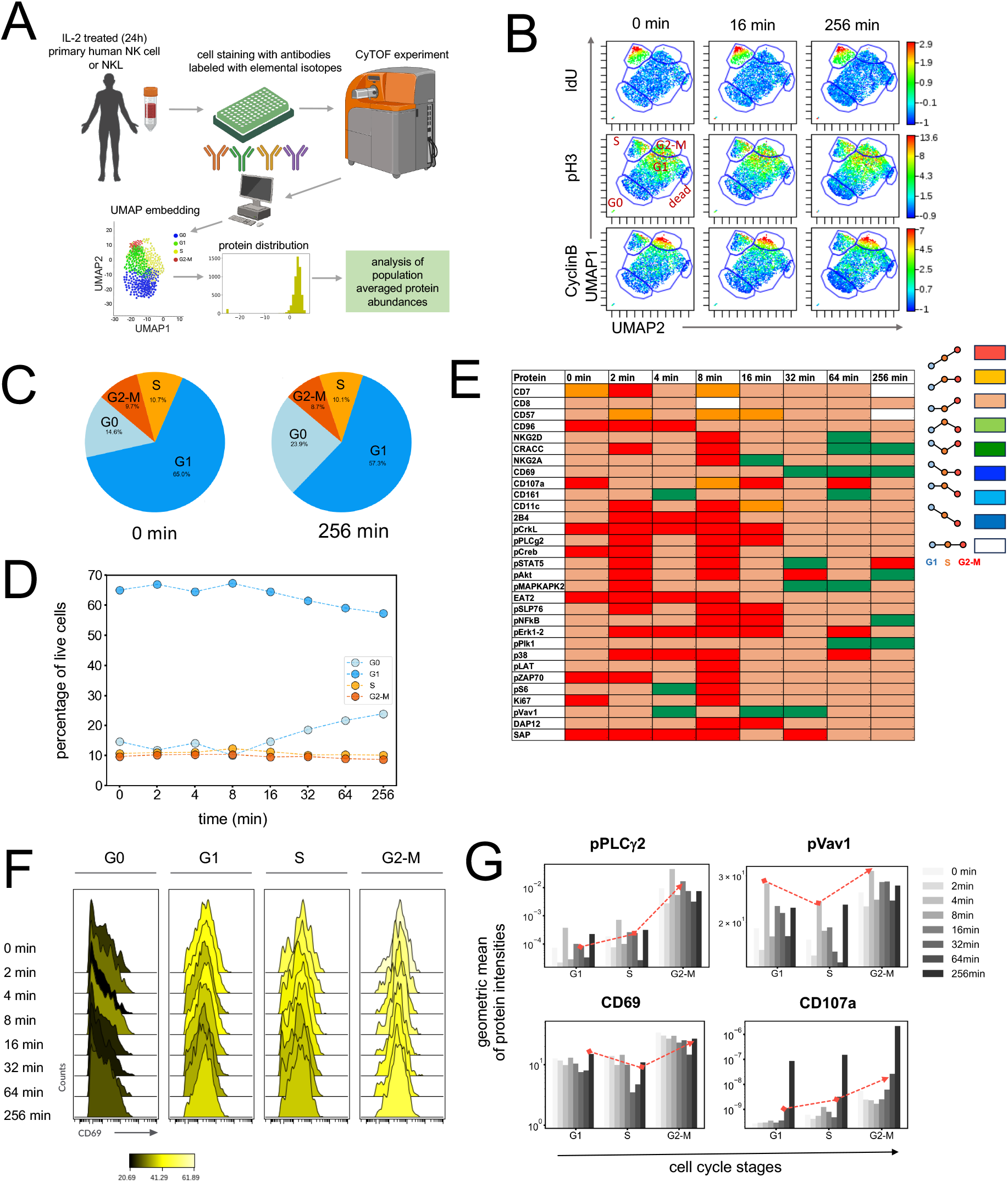
Variation of the abundances of signaling proteins in NKG2D-stimulated primary human NK cells residing in different cell cycle stages. **(A)** IL-2 treated primary CD56^dim^ NK cells were analyzed by CyTOF at 0 (i.e., unstimulated), 2, 4, 8, 16, 32, 64, and 256 min post-NKG2D stimulation. Cell populations were gated using UMAP embeddings into various cell cycle stages for different time points. Protein intensities follow asymmetric log-normal distributions across cell cycle stages. We log-transformed the protein intensity data to compare mean protein abundances in cell populations at different cell cycle stages (e.g., G1 and S or S and G2-M). We used one sided *t-*test for the comparison. **(B)** Cell populations are gated into G1, S and G2-M stages for different time points using cell cycle markers (see Methods). **(C)** Proportion of cells (in percentage) at different cell cycle stages are shown at 0 and 256 min post-NKG2D stimulation. **(D)** Proportion of live CD56^dim^ NK cells in G0, G1, S and G2-M stages are shown at the different time points post-NKG2D stimulation. **(E)** The variations of the mean log_e_ (Intensity) of signaling proteins across cell cycle transitions, G1→ S, and S → G2+M, are shown in terms of nine different types of variations. A change (increase or decrease) is marked when the change in the mean log_e_ (Intensity) from G1→ S, or S→G2+M, has a statistical significance level *α* = 0.05 in one sided *t-* test. The nine types are characterized by the changes in (G1→ S, S → G2+M) as follows: (increase, increase), (increase, no-change), (no-change, increase), (increase, decrease), (decrease, increase), (decrease, no-change), (no-change, decrease), (decrease, decrease), (no-change, no-change). **(F)** Shows histograms of protein intensities for untransformed (raw) data at different time points across cell cycle stages. The colors of the histograms represent the arithmetic mean. **(G)** The variation of geometric mean of protein intensities per cell are shown at different time points post-NKG2D stimulation (see label) for pPLCγ2, pVav1, CD69 and CD107a. Red dashed lines are used to guide monotonic increase of pPLCγ2 at 16 min and CD107a at 64 min post-NKG2D stimulation, non-monotonic behavior (decrease-increase) of pVav1 and CD69 at 4 min and 256 min post-NKG2D stimulation across cell cycle stages, respectively.

To investigate differences in the interplay between NKG2D signaling and cell cycle progression between NK cell lines and primary NK cells, we analyzed NKG2D signaling kinetics in a human NK cell line, NKL, which constitutively expresses the activating human NKG2D receptor. The NKL cells were stimulated by plate-bound anti-NKG2D. A similar analysis of changes in average signaling protein abundances with cell cycle states revealed a larger fraction (~15%) of proteins displaying non-monotonic variations post-NKG2D stimulation (**Supplementary Figure S3**).

Both in NKL and primary NK cells stimulated with anti-NKG2D, the early activation marker CD69 exhibited non-monotonic behavior at a few time points late in the signaling kinetics. For all other earlier time points, CD69 displayed semi-monotonic behavior across cell cycle stages in primary NK cells as opposed to no changes in the abundances across cell cycle stages in NKL cells (**Supplementary Figure S3**). The abundances of the degranulation marker CD107a increased semi-monotonically in both NKL and primary NK cells for most time points during NKG2D signaling kinetics. However, in NKL cells, CD107a showed a non-monotonic behavior, with the highest expression at 256 minutes. The abundance of the signaling protein pS6 showed non-monotonicity in both NKL and primary NK cells at certain time point and showed monotonic or semi-monotonic behavior at most timepoints in primary NK cells, whereas in NKL cells, there were no changes in the abundances.

In addition, we engineered the NKL cells to express activating mouse receptor Ly49H to study another NKR signaling kinetics that uses the ITAM-bearing adaptor and the DAP10 adaptor, in contrast to human NKG2D receptor signaling, which solely uses the DAP10 adaptor^12,26^. Anti-Ly49H antibodies were used to stimulate Ly49H-expressing NKL cells, and a similar analysis to the one above showed a higher fraction (~17%) of signaling proteins displaying non-monotonic changes compared to those in NKG2D-stimulated primary human NK cells (**Supplementary Figure S4**). A marked difference between NKG2D stimulated primary NK cells and NKL cells with Ly49H stimulation was no changes in the abundances of CD69 or CD107a across cell cycle stages for the majority of the time points during the signaling kinetics (**Supplementary Figure S4**). A few proteins showing non-monotonic changes with cell cycle stages upon Ly49H stimulation, which differed from the behaviors observed in NKG2D-stimulated NKL cells included pSTAT1, DAP12, and CCR7.

The monotonic and non-monotonic variations in the average protein abundances with progressing cell cycle states can arise due to various mechanisms as cells progress through cell cycle states such as (a) variations in the average abundances of interacting signaling protein (i.e., substrate, kinase or phosphatase), (b) increase in the cell size (volume), and (c) potential interactions of CDKs with cell signaling proteins. To elucidate mechanisms how these processes influence the abundances of signaling proteins at different cell cycle stages, we developed a set of minimal models that consider basic components of biochemical signaling reactions coupled with cell cycle processes. Furthermore, each of these processes is stochastic in nature, giving rise to randomness in both the duration of cell cycle stages and the production of signaling proteins. Therefore, we considered stochastic time evolution of our model systems. We developed our models in two steps: first, we considered models that can be solved analytically, and exactly and then we developed a stochastic simulation method, validated by the analytical calculations, to simulate the model kinetics. Next, we extended our models to include non-linear biochemical reactions, which were numerically simulated using our stochastic simulation approach.

### Development of minimal models that couples first-order cell signaling reactions and cell cycle processes

We developed a minimal model that combines signaling reactions with processes involved in cell cycle transitions. In this model, an unphosphorylated protein species A (e.g., Vav1, Erk) undergoes post-translational modifications such as tyrosine phosphorylation through a first-order reaction. Such a protein phosphorylation can occur during signaling kinetics initiated by the stimulation of NK cell receptors (e.g., NKG2D) and cytokine receptors (e.g., IL-2R). The phosphorylated form of A (denoted as A*) is dephosphorylated by phosphatases in the cell. The protein species A can be produced or degraded in individual cells residing in different cell cycle states. The synthesis, degradation, phosphorylation, and dephosphorylation of protein species A are modeled by zero- and first-order reactions occurring at rates r, δ, r_p_, and r_d_, respectively (**Figure 3A**). We represent the parameter set {r, δ, r_p_, r_d_} by a parameter vector ***θ*** for brevity. This model is amenable to exact analytical calculations, which helps in clearly deciphering the interplay between cell signaling and cell cycle processes. The description of phosphorylation/dephosphorylation reactions by first-order reactions is reasonable when the abundances of kinases and phosphatases are in excess compared to the substrate abundance^27^. The details of the rates for the reactions are provided in the captions of **Figure 3A**.

**Figure 3:**
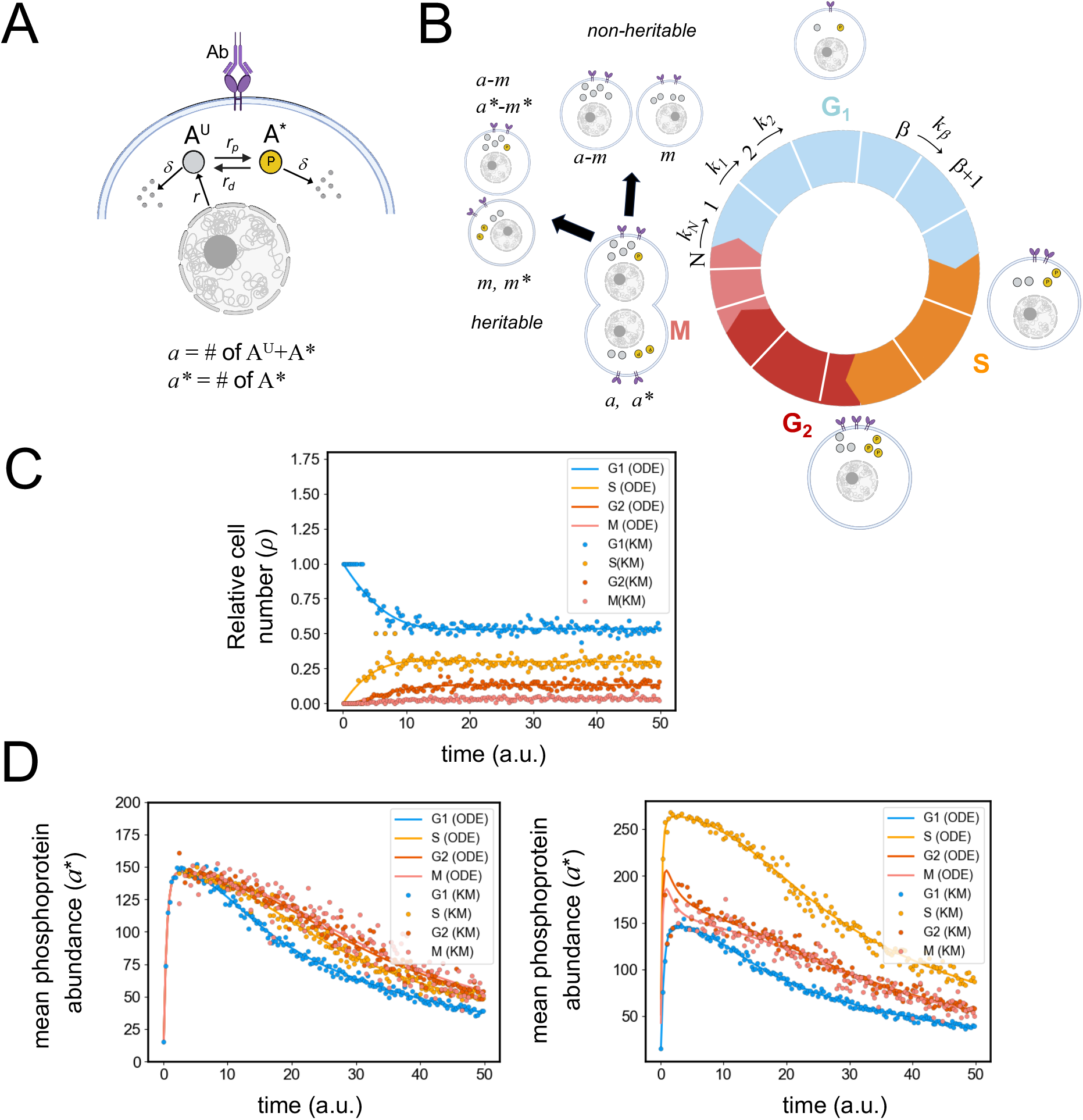
Kinetics of average protein abundances in single cells residing in different cell cycle stages for a first-order signaling reaction model. **(A)** Schematic representation of a protein phosphorylation-dephosphorylation model involving first-order reactions. Receptor signaling (implicit) leads to phosphorylation of a protein A^U^ (unphosphorylated form) to A* (phosphorylated form). A^U^ is synthesized inside the cell and both A^U^ and A* can be degraded in the cell. The rates of the reactions are shown in the schematic figure. **(B)** Schematic representation of a cell going through different cell cycle stages: G1, S, G2 and M. Each cell cycle stage consists of multiple intermediate stages, where the cell transitions from stage *β* to stage *β*+1 at a rate k_β_. The protein content of the mother cell in the M stage is divided between two daughter cells upon division following a binomial distribution. In the full heritability scenario, phosphorylated proteins in the mother cells are transferred to the daughter cells. In the non-heritability scenario, only the phosphorylated forms of proteins are de-phosphorylated upon transfer to daughter cells. **(C)** Shows the population averaged relative cell number or the proportion of average cell number in each cell cycle state. **(D)** Average protein abundances as a function of time in cell cycle stages for the signaling model shown in (**A)**, obtained from Gillespie simulation (circles) or the numerical solution of the ODEs (solid lines). The left panel shows the case when the reaction rates are independent of the cell cycle stages, while the right panel shows the case when the phosphorylation rate of A^U^ is increased 10 times in the S state. We used the following model parameters for the results shown: (left panel) r = r_p_ = r_d_ = 1.0, δ = 0.01, *k*_*β*_ = [0.07, 0.1, 0.2, 0.7] in arbitrary units (a.u.), and (right panel) same as in the left panel, except, r_p_=10.0 in the S state.

Next, we combined the above signaling, protein production and degradation processes with cell cycle transitions (**Figure 3B**). We considered that single cells can be in multiple virtual cell cycle states {1, 2,…, *β*,…, *N*}, where the cell cycle transitions between these states occur at the rates described below:

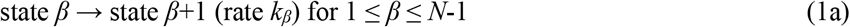

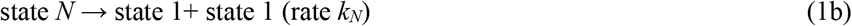

These transitions are described as Markovian^28^ processes where the transition to the next step depends only on the current state and the residence time of the single cells in each cell cycle state β follows an exponential distribution. The total number *N* of virtual cell cycle states and the transition rates {*k*_*β*_} can be chosen to fit experimentally measured probability distribution functions of cell division times^17^. The states {β} describe a collection of intermediate virtual states that correspond to a specific cell cycle stage (e.g., S stage). The durations of these intermediate virtual states are exponentially distributed and the durations of cell cycle states consisting of multiple of these virtual states follow an Erlang distribution^29^. This construct effectively models the residence times of cell cycle states observed in experiments^29^. Similar mathematical models have been widely used to describe cell cycle kinetics^30-32^. In our model, as a single cell transitions from the current state β to the next cell cycle state β+1, the abundances of unphosphorylated and phosphorylated forms of A in state β, given by (*a*^U^)_*β*_ and (*a*^*^)_*β*_, respectively, are fully transitioned to the state β+1. When the mother cell splits into two daughter cells, described by the N → 1 transition, the total number of A molecules in a single cell in the terminal cell cycle state N (i.e., *a*_*N*_ *=* (*a*^U^)_*N*_ + (*a*^*^)_*N*_) is split into the two daughter cells with abundances *m* and *a*_*N*_*-m* in cell cycle state β =1, with a binomial probability 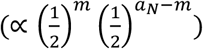 This assumption has been supported by experiments and has been incorporated into mathematical models describing cell cycle kinetics^33,34^. Single cell experiments in yeast cells responding to stress show the heritability of phosphorylated forms of proteins during mitosis^35^; therefore, we considered this possibility in our model. The transition of the phosphorylated protein abundance (*a*^*^)_*N*_ in state N to the daughter cells is considered in two ways (**Figure 3B**): (1) <u>No heritability</u>: The newly generated daughter cells at state 1 do not retain any phosphorylated A* molecule from the mother cell at state N. (2) <u>Full heritability</u>: The number of phosphorylated A* molecules (or *a*\*) in the mother cell at the time of cell division is split into *m\** and (*a*\*)_N_ *-m\** in the daughter cells, with a binomial probability 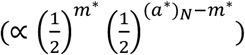 (**Figure 3B**).

The time scales pertaining to protein phosphorylation/de-phosphorylation occur on the order of seconds to minutes^4,5,36,37^, whereas mammalian cells usually duplicate in 24 hours^38^. In such cases, the time scales for progression in cell cycle states and signaling kinetics appear to be well-separated. However, the synthesis and degradation of some proteins can occur over longer time scales that can overlap with the above ranges, e.g., protein production can take anywhere from ½ hour to tens of hours^39^ and protein degradation via ubiquitylation can occur within ~1 hour to a few hours^40^. Further details regarding these time scales are provided in **Table 1** and **supplementary Table S1**. Furthermore, the protein synthesis and degradation rates can vary across cell cycles states, and CDK proteins can actively participate in post-translational modifications of signaling proteins – all of which could contribute to variations in the abundances of signaling proteins across cell cycle states. To investigate the roles of these different mechanisms, we considered two scenarios (*neutral* and *active*) regarding potential cell cycle dependence of synthesis, degradation, phosphorylation, and dephosphorylation rates of signaling proteins in our models. (1) *Active* case: In this scenario, these rates assume different values ({***θ***^*β*^}) for different cell cycle states β representing active participation of cell cycle processes in regulating signaling protein production and signaling kinetics. (2) *Neutral* case: In this case, the rates ***θ*** are the constant across all cell cycle states, representing no direct regulation of protein synthesis/ degradation/ signaling by cell cycle processes.

**Table 1:** Estimation of values of the model parameters.

| Parameter | Values | Notes |
| --- | --- | --- |
| $k_1, k_2, k_3, k_4$ | $k_1 \sim 0.07 \text{ hr}^{-1}$, $k_2 \sim 0.1 \text{ hr}^{-1}$, $k_3 \sim 0.2 \text{ hr}^{-1}$, $k_4 \sim 0.7 \text{ hr}^{-1}$. | NKL cells grow in the presence of IL-2 in vitro with a doubling time of its 24-48hrs<sup>8</sup>. Primary NK cells obtained from healthy adults expand ex-vivo in the presence of artificial antigen presenting cells and IL-2 with a doubling time of about 30 hours<sup>47</sup>. However, the cell cycle residence times in G1, S, G2, and M are not known. We used the cell cycle residence times of HeLa cells in G1, S, G2, and M states which are $\sim 10\text{hr}$, $6\text{hr}$, $3\text{hr}$, and $1\text{hr}$, respectively<sup>7</sup>, of the total cell cycle duration of 20 hours to evaluate the fractions of the cell cycle duration spent in each state and assumed cell cycle times in NK cells roughly follow the same fractions. If we assume $k_1$ to $k_4$ correspond to the durations of G1, S, G2, and M states, then the estimated rates $k_1$ to $k_4$ for NK cells are for the above case are $0.07 \text{ hr}^{-1}$, $0.1 \text{ hr}^{-1}$, $0.2 \text{ hr}^{-1}$, and $0.7 \text{ hr}^{-1}$. |
| $r$ | 10800 – 3240000 molecules $\text{hr}^{-1}$ | The values are based on measurement of protein turnover in activated CD4+ T cells<sup>48,49</sup>. The synthesis rates show large variations, e.g., CD69 synthesis rates is $\sim 180$ molecule number per min in activated CD4+ T cells whereas β-actin shows a high synthesis rate of 54000 molecule numbers/min. |
| $\delta$ | 0.03- 0.9 hr<sup>-1</sup> | Lysosome and proteasome mediated degradation occur with shorter time scales (half-lives 45min – few hours in HeLa cells) whereas loss of protein abundance due to dilution (or cell replication) occurs in cell cycle time scales (e.g., > 22hr for HeLa cells). Therefore, protein decay rates vary from 0.03-0.9 hr<sup>-1</sup> in HeLa cells<sup>50</sup>. In activated CD4+ T cells some proteins (e.g., $\beta_2$M) showed fast turn-over with a half-live ($t_{1/2}$) of 45 mins, proteins such as CD3ε turned over in $t_{1/2} \sim 11$ hours, and proteins like histones showed $t_{1/2} > 1000$ hour turn over times<sup>48,49</sup>. The half-lives for protein turnover reflect degradation time scales. |
| $r_p$ | 36-7200000 hr<sup>-1</sup> | The rates are $\sim 0.01$-2000 s<sup>-14,5,36,37</sup>. |
| $r_d$ | 36-7200000 hr<sup>-1</sup> | The rates are $\sim 0.01$-2000 s<sup>-1 4,5,36,37</sup> |
| $k_{on}$ | 10<sup>5</sup>-10<sup>8</sup> M<sup>-1</sup> s<sup>-1</sup> | The association range could be 10<sup>5</sup>-10<sup>8</sup> M<sup>-1</sup> s<sup>-1 51</sup>. In T cell, the association rate of ZAP70 to the ITAM is 5 μM<sup>-1</sup> s<sup>-1</sup> = 5 uM<sup>-1</sup> s<sup>-1</sup> = 29.90 μm<sup>3</sup> h<sup>-1</sup> per molecule<sup>52</sup>. |
| $k_{off}$ | 450-7200 hr<sup>-1</sup> | The rates are $\sim 0.125$-2 s<sup>-1</sup>. In T cell, ZAP70 unbinding rate from ITAM 0.125 s<sup>-1 52</sup>. SHP-1 unbinding rate from ITAM is 1.7 s<sup>-1 53</sup>. |

We set up a Master equation^28^ to describe the stochastic kinetics involving the cell signaling and cell cycle transitions as described above. The Master equation was then employed to analytically evaluate the kinetics of population-averaged protein abundances in each cell cycle stage (details in the **Supplementary Text 1**, sections 1-4). In parallel, we developed a stochastic simulation approach to numerically simulate the stochastic trajectories of single-cell abundances of A and A* as the cells progress through cell cycle stages. The details regarding the algorithm are provided in the Materials and Methods section. In the model, the total number of cells increases continuously with time due to cell division; however, the population-averaged number of cells in any given cell cycle state β relative to the total number of cells across all cell stages, denoted by *ρ*_*β*_ (see in Materials and Methods), reaches a stationary state after a sufficiently long period. The time evolution of *ρ*_*β*_ for the states β =1 to N in the model is described by a set of coupled non-linear ordinary differential equations (ODEs) (**Supplementary Text 1**, section 5)

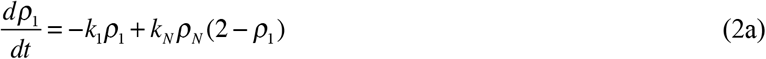

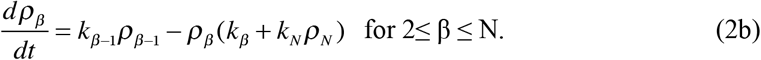

The above non-linear ODEs can be solved numerically. We compared the numerical solutions of the ODEs with the numerical simulations of the stochastic kinetics, which showed excellent agreement (**Figure 3C**). The steady states of the above ODEs can be analyzed analytically. The steady state of the above kinetics (see Materials and Methods section) shows that the longer the duration of a cell cycle state, the larger the fraction of cells residing in that state (**Supplementary Text 1**, section 6). The kinetics of the population-averaged abundance of the total amount of protein A = A^U^ + A* (= *a*_β_) and phosphoprotein 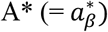 in any given cell cycle state β depends on the rates (***θ***^*β*^) of synthesis, degradation, phosphorylation, and de-phosphorylation in state β, as well as on the cell cycle transition rate from β − 1 → β, and the fraction of the cells in both the previous state (β-1 for β >1, and, N for β =1) and the current (state β) cell cycle state. The kinetics of *a*_β_ (**Supplementary Text 1**, section 5) is given by,

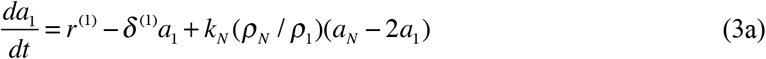

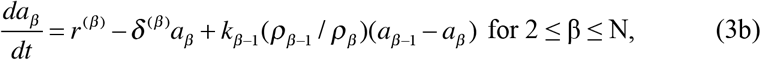

and the kinetics of 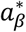 with full heritability of *a\** is given by,

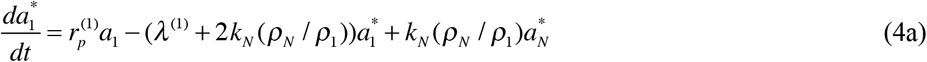

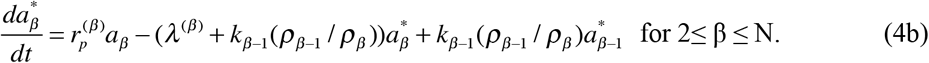

Where 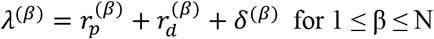.

Here, the rates of protein synthesis, degradation, phosphorylation, and dephosphorylation are denoted by 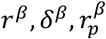 and 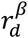, respectively, in state β. The above system of non-linear ODEs can be solved numerically, and the steady-state behavior can be analyzed analytically (see Materials and Methods, **Supplementary Text 1**, section 5). The numerical solution of the above ODEs showed excellent agreement with our stochastic simulations (**Figure 3D**). Below, we describe the dependencies of the steady-state average abundances of A and A* on the kinetic rates pertaining to the cell cycle, protein synthesis and degradation, and phosphorylation and dephosphorylation processes.

In the absence of any cell cycle transitions, the steady-state value of *a* is fully determined by the synthesis and degradation rates *r* and *δ*, respectively, *i*.*e*., 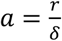 (**Supplementary Text 1**, section 9). In the presence of cell cycle transitions, the steady state average abundance *a*_*β*_ depends on the synthesis and degradation rates *r*^(β)^ and *δ*^(β)^, as well as on the steady state values of the average per-cell abundances of A in the preceding cell cycle state with a weight factor < 1 (see Materials and Methods, **Supplementary Text 1**, section 5, Eqn. 57). We can make few general observations from our analysis of the steady state of *a*. (i) *Neutral* case: For protein synthesis and degradation rates that do not change with cell cycle states, i.e., *r*^(*β*)^ = *r* and *δ* ^(*β*)^ =*δ*, the average abundance per cell *a*_*β*_ increases monotonically as the cell progresses through the cell cycle states when the protein degradation rate *δ* is slower than the cell cycle transition rates *k*_*β*_. However, when the rates of protein synthesis and degradation are substantially larger than the cell cycle transition rates, i.e., *δ* ≫ *k*_*β*_, the average protein abundances per cell *a*_*β*_ are roughly equal in all cell cycle states in this case. (ii) *Active* case: When the rates r and *δ* depend on the cell cycle states, there can be non-monotonic changes in the steady state-values of *a*_*β*_ with progressing cell cycle states.

Next, we analyzed the average abundances 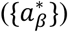 of the phosphorylated protein species A in the steady state. Using similar arguments as for {*a*_*β*_} we can make the following general observations.

#### 1. Neutral case

When the protein synthesis/degradation and phosphorylation/dephosphorylation rates do not change with cell cycle stages, i.e., 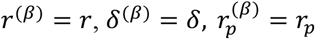, and, 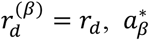 increases monotonically as the cell progresses through the cell cycle when the protein degradation rate *δ* is slower than the cell cycle transition rates *k*_*β*_, tracking the increase in *a*_*β*_ as the cells advance. If *a*_*β*_ remains unchanged with cell cycle stages, 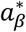 also remains unchanged.

#### 2. Active case

Any decrease or non-monotonic change in 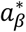 results from cell cycle-dependent rates of protein synthesis/degradation and/or phosphorylation/dephosphorylation. Therefore, any decrease or non-monotonic change in the average abundance of phosphorylated protein results from changes in protein synthesis/ degradation rates and/or phosphorylation/ dephosphorylation rates, which are influenced by cell cycle states. These variations in the rates could involve direct participation of proteins (e.g., CDKs) that regulate cell cycle transitions.

In the next section, we investigate coupling of a basic non-linear biochemical signaling reaction with cell cycle processes. The model cannot be solved analytically; therefore, we use stochastic simulations of the model for our investigation.

### Modeling coupling of non-linear biochemical signaling reactions with cell cycle processes

We considered phosphorylation of a substrate protein species A (e.g., Vav1) by a kinase E (e.g., Src kinase), where the kinase binds to the substrate via a second-order reaction and then phosphorylates it (**Figure 4A**). The phosphorylated substrate A* is dephosphorylated by phosphatases, which is modeled as a first-order reaction. The above reactions are common in many NK cell receptors including NKG2D receptor-initiated signaling. Because the propensity of a second-order reaction, which determines how frequently the reaction occurs, decreases with increasing cell volume (V) as ∝ 1/V, we considered the increase in the volume of a single cell in our stochastic simulation as it progresses through the cell cycle. We also considered for the production and degradation of the substrate A, and the kinase E in the different cell cycle stages, which are modeled by zero- and first-order reactions. The changes in the abundances of A, A*, E, and the intermediate complexes EA as cells transition through cell cycle stages are modeled in the same way as described in the previous section. Further details are provided in the Materials and Methods section.

**Figure 4:**
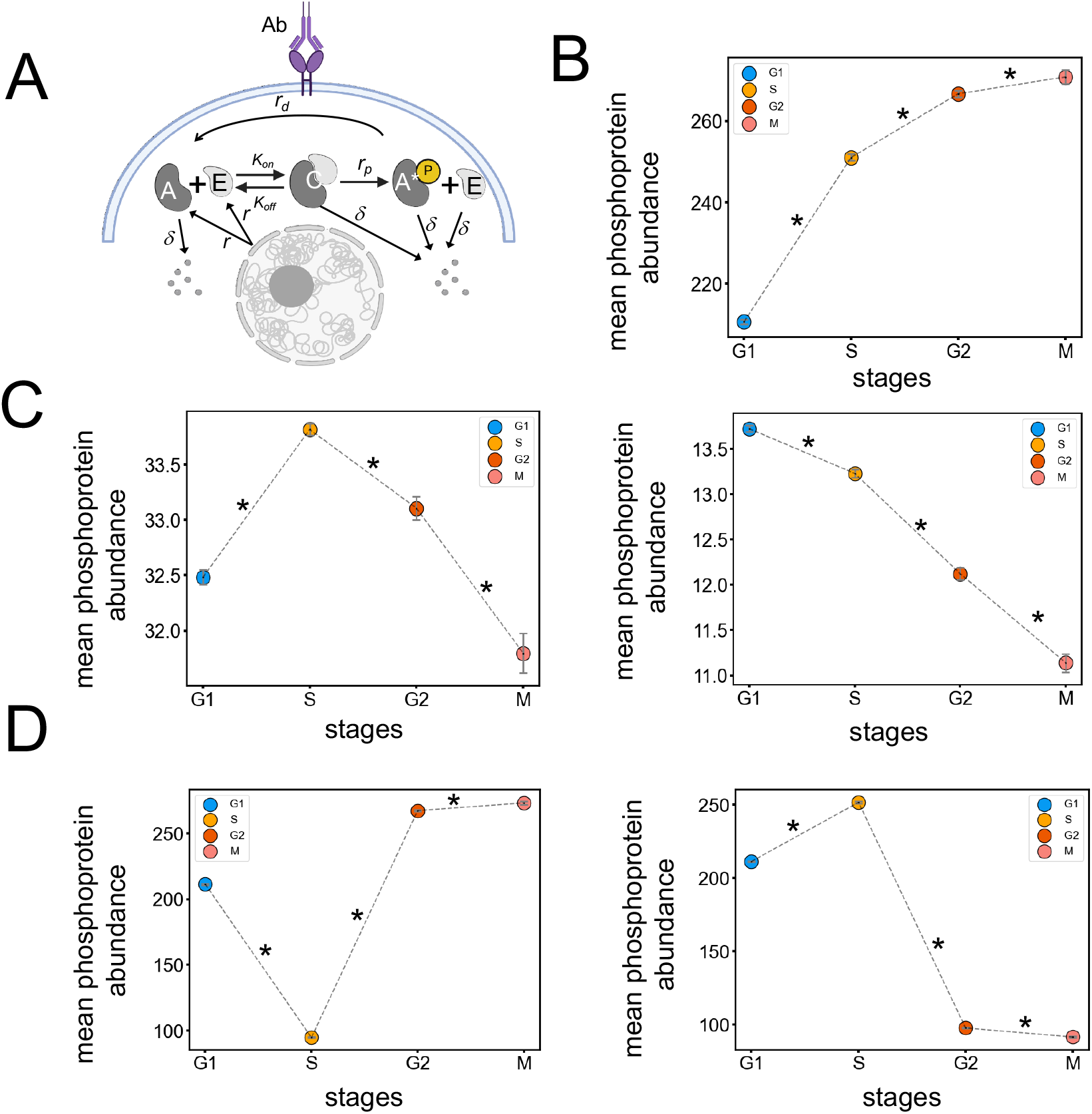
Kinetics of average protein abundances in cells residing in different cell cycle stages for a non-linear signaling reaction model. **(A)** Schematic representation of a non-linear signaling reaction model. Enzyme E and its substrate A bind to form a complex C, that gives rise to phosphorylated protein A* and the enzyme E is released. The proteins A, E, C and A* can be degraded, and the proteins A and E are produced in the cell with zeroth-order reactions. Rates associated with the reactions are shown above arrows. **(B-D)** Shows the average abundances of A* at the steady state for the signaling model in (A) at cell cycle stages G1, S, G2, and M. The results are shown for the *neutral* (**B** and **C**) and the *active* **(D)** scenarios. The *neutral* case can give rise to monotonic increase **(B)**, non-monotonic change (**C**, left panel), and monotonic decay (**C**, right panel) in the average abundance of A* for G1 → S → G2. The non-monotonic change (**C**, left) or the monotonic decay (**C**, right) can be observed in the *neutral* case within a narrow parameter range. In the *active case* the rate of protein phosphorylation is decreased 100-fold in S stage (**D**, left) which produces a non-monotonic change (decrease, increase) for G1 → S → G2 in the average abundance of A*. Increasing the rate of phosphorylation of A* 100 times in S stage and reducing it 100 times in G2 and M stages generates another type of non-monotonic change (increase, decrease) with progressing cell cycle stages for G1 → S → G2 (**D**, right). For **B-D**, the mean and the standard error of the mean (SEM) of the average abundance of A* are calculated from 20 simulation replicates. Asterisks represent the statistical significance in *t-*test with *P*-value < 0.05 which we calculated independently between G1 and S stage, S and G2 stage, and G2 and M stages with *n* = 20 samples of the average abundance of A* in each cell stage. Parameter values used and *P*-values calculated from *t-*tests in **B-D** are provided in **Supplementary Table S2**.

We then simulated the above reactions shown in **Figure 4A** to evaluate the variation of the cell population-averaged abundances of 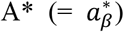 across cell cycle stages and the underlying mechanisms. We investigated both the *neutral* and the *active* scenarios as described previously for the first-order signaling model, where the rates of protein synthesis, degradation and the rates for biochemical signaling reactions (i.e., phosphorylation/dephosphorylation/association/dissociation) are neutral or depend actively on cell cycle states. We observed the following patterns in the variations of the per cell average abundances 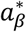 in steady state as cells progress through cell cycle states (details in **Supplementary Text 1**, section 11 and **Supplementary Table S3**).

#### (a) Monotonic increase

In the *neutral* case, the average abundance of 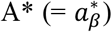 increases from G1→ S → G2 + M when the time scales for protein degradation (∝ 1/*δ*) is similar or larger than that of the cell cycle transition time scales, i.e., 1/*δ* ≥ {1/*k*_*β*_}. In this situation, the total abundances of the substrate, intermediate complex and product (A_total_ = A + A* + EA), and the enzyme (E_total_ = E + EA) increase as the cell progresses through cycle states, with each cell cycle state accumulating the abundances from previous state. Consequently, as the substrate and the enzyme abundances increase, this leads to greater abundances of A* in successive cell cycle stages compared to the previous one (**Figure 4B**). In the *active* case, we can achieve this behavior for a wider range of rate parameters, e.g., when the rates of protein synthesis/phosphorylation increase with increasing cell cycle states, the kinetics can generate a monotonic increase regardless of the degradation rate.

#### (b) Monotonic decrease

This pattern occurs in the *neutral* case when the rate of protein degradation is much faster compared to the cell cycle transitions (*δ* ≫{*k*_*β*_}). In this scenario, the average per cell abundances of A_total_ and E_total_ reach a steady state at each cell cycle state, and thus do not vary across cell cycle states. However, as the cell volume increases with cell cycle progression, the propensity for the second-order reaction producing the intermediate complex, EA, decreases, leading to the decrease of the abundance of A* (**Figure 4C**). For the cell volume increases by a factor of approximately two before division, the monotonic decrease in the abundance of A* observed in **Figure 4C** is ~ 0.23 fold across all cell cycle stages. This point is further considered in the **Supplementary Text 1**, section 11. For the *active* case, a decrease (or increase) in the synthesis (or degradation) rate or a decrease (or increase) in protein phosphorylation (or dephosphorylation) rate with cell cycle progression can give rise to a substantially large monotonic decrease.

#### (c) Non-monotonic changes

This behavior can occur in the *neutral* case for a narrow range of parameter values within the ranges provided in **Supplementary Table S1**. For instance, this situation can arise when the increase in the propensity of the second-order reaction (e.g., protein binding: E + A → EA) yielding the intermediate complex EA due to the increase in the total abundances of E (E_total_) and A(A_total_) with progressing cell cycle stages is counteracted by the decrease in propensity caused by the increasing cell volume (see **Supplementary Text 1**, section 11 for further details). For specific values of r, δ, and the biochemical signaling rates, we can obtain an increasing abundance of A* from G1→ S, and then decrease from S → G2 + M (**Figure 4C**). However, the magnitudes of protein variation are small ~ 0.06 fold **(Figure 4C)**. The non-monotonic changes are more easily obtained in the *active* case by cells introducing cell cycle-dependent rates of protein synthesis, or degradation, or biochemical signaling rates (**Figure 4D**).

#### (d) Unchanged abundances

This behavior can be obtained in the *neutral* case when the degradation of E and A are faster compared to the cell cycle transitions (*δ* ≫{*k*_*β*_}) that leads to the constant per cell abundances of A* across cell cycle stages when cell volume does not change across these stages. When the growing cell volume is taken into consideration, a high binding affinity between the E and A (or low dissociation constant *K*_*D*_) reduces the effect of increasing cell volume on the reaction propensity and consequently on the abundances of A*. As a result, the protein abundances remain unchanged across cell cycle stages (**Supplementary Text 1**, section 11, **Supplementary Table S3**). Similarly, in the *active* case, if the cell cycle dependencies of the synthesis/degradation/biochemical signaling rates are tuned such that those counteract the increases or decreases in abundances of A* between consecutive cell cycle stages obtained in either the *active* or the *neutral* case as described before.

To quantify the proportion and robustness of different types of phosphoprotein variation (monotonic vs. non-monotonic) in the *neutral* scenario, we initially performed numerical simulations within a specific range of parameters (~ 20-30 fold variation in k_on_ and *δ*, see **Supplementary Table S2C**). These simulations resulted in ~26% non-monotonic cases and ~2% monotonic decrease cases, with the monotonic decay and non-monotonic variation being at most ~ 0.2 fold and ~ 0.1 fold, respectively. Furthermore, we developed approximate analytical expressions to estimate variations of the abundances with cell cycle states across parameter ranges. Details on the maximum variation in protein abundances in the *neutral* scenario are provided in **Supplementary Text 1**, Section 11 (**Supplementary Figure S9)**.

## Discussion

NK cells proliferate in response to NKR and cytokine stimulations. As proliferating NK cells progress through cell cycle states from G1→ S→ G2→ M, many proteins including those involved in NKR and cytokine signaling, are synthesized acquiring different abundances in each cell cycle state. In addition, physical characteristics of dividing cells, such the cell volume that can affect the propensity of signaling reactions and the abundances of signaling products increase with cell cycle progression. Our investigation of the variations in the abundances of NKG2D signaling proteins with cell cycle states, using CyTOF experiments in IL-2-treated primary CD56^dim^ NK cells showed monotonic or semi-monotonic increases, non-monotonic changes, and no-changes in the population-averaged per cell abundances of phosphorylated signaling proteins with progressing cell cycle states. The majority (~ 89%) of the 31 measured proteins (e.g., CD107a, pCrkL, and pPLCγ2) at various timepoints in the NKG2D-stimulated primary CD56^dim^ NK cells showed monotonic or semi-monotonic increases. Whereas a small proportion (~ 9%) of the proteins, such as pVav1 and pNFκB, exhibited non-monotonic changes at specific time points during NKG2D signaling. NKG2D stimulation of human NKL cells constitutively expressing NKG2D receptor by plate-bound antibodies showed a higher proportion (~15%) of the proteins showing non-monotonic changes such as CD69 and pS6. The abundances of these proteins show a higher frequency of semi-monotonic behavior in primary NK cells compared to NKL cells. Although the fraction of cells expressing high abundances of CD107a was larger in NKL cells compared to primary NK cells (**Supplementary Figure S6**), the variation in CD107a abundances with cell cycle stages was qualitatively similar between NKG2D stimulated primary NK cells and NKL cells. For Ly49H-stimulated NKL cells, the proportions of non-monotonic changes increased (~17%) compared to the primary NK cells; CD107a or CD69 abundances showed no changes across cell cycle stages in contrast to NKG2D-stimulated NKL cells. The increased instances of non-monotonic changes as well as differences in cell cycle dependence of particular signaling proteins (e.g., pS6) in the NKL cells suggest increased influence of cell cycle driving proteins on NK cell signaling kinetics. In addition, the specific proteins that show qualitative differences in changes in protein abundances with cell cycle stages in primary CD56^dim^ NK cells post-NKG2D stimulation and the NKL cells post-Ly49H stimulation, which could potentially be due to differences in NKR signaling (e.g., DAP10 versus DAP12 adapter signaling).

We explored mechanisms that can make the NK cell signaling kinetics dependent on cell cycle states, specifically, when the time scales for cell cycle transition are about an order of magnitude longer than those of the signaling kinetics. To investigate this, we developed minimal models that can describe enzymatic chemical modifications (e.g., phosphorylation or dephosphorylation) of signaling proteins. Using both analytical calculations and simulations, we examined the coupling of biochemical signaling kinetics and the physiochemical changes that are associated with cell cycle transitions.

Our analyses for the *neutral* case revealed that when the degradation rates of enzyme (e.g., kinase and phosphatases) are much smaller than the cell cycle transition rates, the abundances of the phosphorylated forms of the substrate increased monotonically as the cell cycle progresses following the increase in the abundance of enzyme and substrate. In contrast, when protein degradation rates exceed cell cycle transition rates, the changes in the abundances of the phosphorylated signaling protein (product) with cell cycle progression is negligible if the cell volume remains unchanged with cell cycle progression. However, as the cell volume approximately doubles during cell cycle progression, the rate of binding of the enzyme and the substrate effectively decreases by a factor of approximately two with cell cycle stages resulting in the decrease of the abundance of the phosphoprotein. This can lead to a monotonic decay (decrease-decrease) in the abundance of the phosphorylated substrate. In the situation when the degradation rates are comparable to the cell cycle transition rates, the abundances of the enzyme and substrate increase progressively with cell cycle transition. This can overcome for the effective decrease in the rate of binding by the volume increase for limited range of parameter values (see **Supplementary Table S2**) and lead to an increase in the abundance of the phosphoprotein. A non-monotonic variation (e.g., increase-decrease) in the abundance of the phosphoprotein can arise when the above compensation takes place for G1→ S but not for S → G2. Our modeling within a specific parameter range (**Supplementary Table S2C**) showed that such non-monotonic protein variation occurs in about 26% of cases, indicating that it less likely to occur in nature compared to monotonic or semi-monotonic protein variation cases. In contrast, for the *active* case, we found that the active influence of cell cycle regulating proteins generate non-monotonic variations in phosphoproteins in NK cell signaling for a wide range of parameters and variations of the proteins across cell cycle stages are much larger (~ several folds). Thus, the presence of such non-monotonicity is likely to indicate active modulation of signaling kinetics by cell cycle processes.

The time scale NK cell proliferation is about an order of magnitude slower than that (~ 1/mins to 1/secs) of the signaling kinetics – thus the signaling kinetics in a cell cycle state was mainly influenced by the protein synthesis and degradation rates in the cell cycle state. However, for lymphocytes such as T cells, the cell division can occur in much shorter (~ 2-4 hours duplication time^41^) time scale in the expansion phase. In such cases the cell cycle transition and signaling kinetic time scales can be less separated in magnitude, and the protein synthesis/degradation rates as well as the abundances of phosphorylated forms of the proteins in several preceding states can influence the signaling kinetics in the current state. This will be an interesting future direction. We analyzed simple enzymatic signaling reactions in our model. However, certain signaling reactions can involve positive feedbacks where the product promotes its own production and display digital response to stimulations, e.g., Ras activation in T- and B-cells showing a digital response depending on abundances of RasGEFs such as SOS1 and Rasgrp1^42^. As the abundances of substrates and enzymes can vary across cell cycle states, it would of interest to investigate if such digital response occurs in specific cell cycle states.

### The limitation of the study

For primary NK cells, expressions of some of the cell cycle marker proteins (e.g., pH3) were low which made it difficult for us to separate G2 and M stages in gating the cell populations using UMAP embeddings. We manually gated cell populations into different cell cycle stages on the UMAP plane, which resulted in the loss of a small fraction of cell population at the borders between two cell cycle gates. In addition, some the proteins and the timepoints where non-monotonic (or monotonic) changes with progressing cell cycles occurred showed dependence on the choice of the parameters used in CyTOF data analysis (see Materials and Methods and **Supplementary Figure S7**). Thus, confirmation of the underlying mechanisms regarding the cell cycle dependence of the protein abundances requires further experimental validation.

We considered four cell cycle stages in simulating the non-linear signaling kinetics to keep the computation time feasible. This makes the duration of the cell cycle states in the model distributed exponentially; however, the durations of cell cycle states have been observed to follow an Erlang distribution with substantially less variations than the exponential distribution. Since the cell cycle durations are almost an order of magnitude longer than the cell signaling time scales, we reasoned the variations in the durations caused by the exponential distributions will not affect the qualitative conclusions drawn from the model.

## Materials and Methods

### Cell preparation and NKR stimulation

Human peripheral blood NK cells were isolated from blood obtained from healthy adult donors, cultured in IL-2 for 24 hours, and then stimulated with plate-bound agonist anti-human NKG2D mAb as described previously^43^. The IL-2-treated NK cells were stimulated with anti-human NKG2D mAb (5 µg/ml, clone 1D11, BioLegend) for 2, 4, 8, 16, 32, 64, and 256 min. The NKL cells used in this study were generated and maintained as described previously ^44^. The human NKL cell line, which endogenously expresses NKG2D, was transduced with pMX-puro retroviral vectors containing the mouse Ly49H and positive NKL cells were isolated by using flow cytometry sorting^8^. The NKL cells were stimulated with plate-bound agonist anti-Ly49H (5 µg/ml, clone 3D10, eBioscience) mAbs) or anti-NKG2D (5 µg/ml, clone 1D11, Biolegend) mAbs as described previously ^44^ for 8, 32, 64, 128, and 256 min.

### Mass cytometry

Cell staining and mass cytometry (i.e., CyTOF) was performed as described previously^43^. The mass cytometry Ab panels used for the analysis of primary human NK cells and NKL cells are shown in **Supplementary Table S4** and **Table S5**, respectively.

### Gating strategy

Gating of live CD56^dim^ NK cells (i.e., CD45^+^ cisplatin^-^ CD3^-^ CD235^-^ CD61^-^ CD15^-^CD56^dim^ CD16^+^) and live NKL cells (i.e., CD45^+^ cPARP^-^ cisplatin^-^) was done using Cytobank software. Gating of CD56^dim^ NK cell and NKL populations residing in different cell cycle stages was performed using UMAP (nearest neighbor = 15, min distance = 0.01) based on the cell cycle markers Idu, pRb, Cyclin B, pH3, and Ki67, and cell viability marker cisplatin in Cytobank software. We first eliminated dead cells by removing the cisplatin^+^pRb^-^Idu^-^pH3^-^ population. The cell population in the G0 stage was gated based on pRb^-^ Idu^-^ CyclinB1^low^ pH3^-^ expression (**Supplementary Figure S1**). The cell populations in the G1, S, and G2 stages were gated based on pRb^+^ Idu^-^ CyclinB1^low^ pH3^-^ expression, Idu^+^ pH3^-^ expression, and Idu^-^ CyclinB1^high^ pH3^-^ expression, respectively (**Supplementary Figure S1**). In NKG2D-stimulated primary NK cells, we did not see a clear distinction of the pH3^+^ expression; therefore, we could not separate the cell population in the M stage from the G2 stage and designated as G2-M stage. (**Supplementary Figure S1**). Whereas in NKL cells, we identified distinct populations of M-stage cells that are pH3^+^pS6^+^pMAPKAPK2^+22^, although their percentage is very small. Here, we combined these two populations G2 and M into a single G2+M stage and similarly designated it as the G2-M stage.

### Data processing

We exported the untransformed mass cytometry (.fcs) data after splitting it into cell cycle stages. Our gating yielded cell numbers that varied between ~ 400-900 for the G0, S, and G2-M stages and ~ 3000-4000 for the G1 stage. Next, we examined the distribution of intensities corresponding to the expression of signaling proteins in cell populations residing in different cell cycle stages at various time points. The distributions of intensities are asymmetric and contained expressions at negative values. To compare the protein behavior between cell cycle stages (**Figure 2E, Supplementary Figure S3 and S4**) or compute the geometric mean of the protein expression (**Figure 2G, Supplementary Figure S2)** we transformed the intensities to a natural log scale (log_e_). We assigned a small constant value ϵ (10^-11^), less than the minimum positive intensity (~10^-^ _5_), to the negative and zero values as the logarithm of these values cannot be computed (**Supplementary Figure S5**). We checked sensitivity of the protein variation with cell cycle states to the variation of ϵ and the significance level α used in the *t*-test for primary NK cells vs. NKL cells. The qualitative nature of the protein variations in **Figure 2E** did not change for ϵ <10^-11^, but changed at certain time points for those proteins when ϵ was chosen to be higher (~10^-9-^10^-6^) (see **Supplementary Figure S7** for details). In addition, decreasing to α < 0.05 led to qualitative changes in some protein variations compared to that shown in **Figure 2E**.

### Statistical tests

To determine whether the log-transformed intensities of a signaling protein increased, decreased, or did not change between cell cycle stages (**Figure 2E, Supplementary Figure S3 and S4**), we sorted single cells into cell cycle stages. We then performed a one-sided *t-*test comparing the log-transformed intensities of single cells between two cell cycle stages (e.g., G1 and S) to assess whether an increase occurred between the stages (H_1_: μ_2_ > μ_1_). We additionally performed a one-sided *t-*test to determine whether the protein expressions decreased between the cell cycle stages (H_1_: μ_2_ < μ_1_). If neither of these tests achieved the significance level α = 0.05, we classified the protein as having “no change” between the stages. For our stochastic simulation results presented in **Figure 4**, we used similar *t-*tests to compare the phosphoprotein abundances at steady state between cell cycle stages.

### Kinetic equations for average cell populations and protein abundances

We evaluated the kinetics of the total numbers of cells (*n*_*β*_) in any cell cycle state *β*, as well as the total numbers of A (or *A*_*β*_) and A* (or 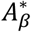) in the cell population residing in the state *β*, using the Master equation^45^ describing the stochastic kinetics for the transitions in Eqns. (5-7). The details of the derivation are provided in the **Supplementary Text 1** (sections 1-4). Here, we present the ODEs describing the above kinetics which show the couplings between the processes regulating the cell cycle and phosphorylation/dephosphorylation kinetics.

The kinetics for {n_*β*_} are described by,

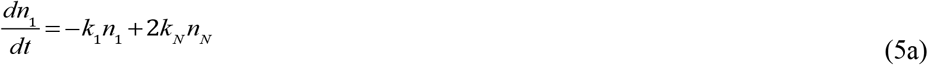

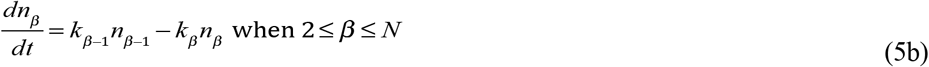

The kinetics for A_*β*_ are given by,

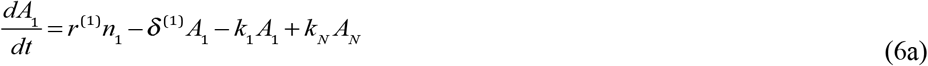

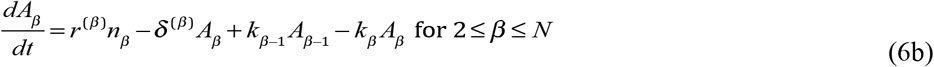

The kinetics for 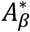 when there is *no heritibility* of A* during cell division are given by,

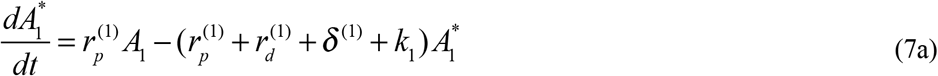

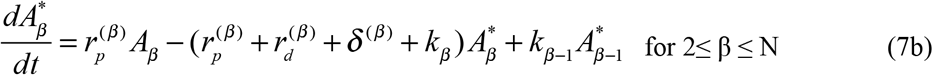

For the case of *full heritibility*, the kinetics of 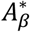 for *β* ≥ 2 is same as that above, and for *β* =1 it is changed to,

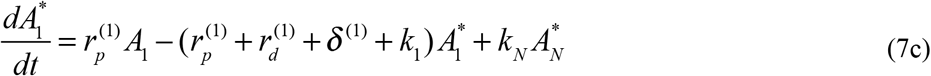

In CyTOF experiments, single cells can be sorted based on expression levels of specific proteins to determine the cell cycle state that single cell is residing in. Abundances of different protein species can be then evaluated in these sorted cells. However, cytometry experiments do not track changes in the total number of cells that can occur due to cell division. Therefore, the relative numbers of cells in different cell cycle states, i.e., *ρ*_*β*_ = *n*_*β*_/(∑_*β*_ *n*_*β*_), and the average abundance of protein species A in a cell cycle state β, given by *a*_*β*_ = *A*_*β*_/*n*_*β*_, can be evaluated from the CyTOF data. The number of cells in each cell cycle state, {n_*β*_}, increase exponentially with time, therefore, the total number of proteins in the cell population, {*A*_*β*_} and, 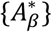 increase exponentially with time as well. However, the relative numbers of cells in any state β, given by 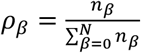 approaches a steady state. The kinetics of {ρ_*β*_}is given by (see **Supplementary Text 1**, section 5.1 for details),

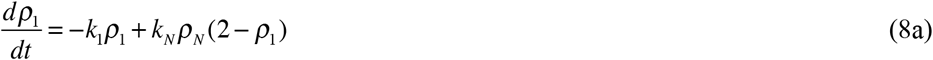

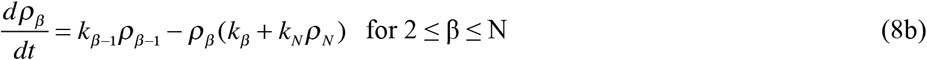

The steady state values are given by,

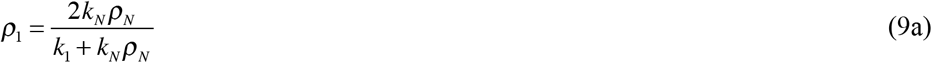

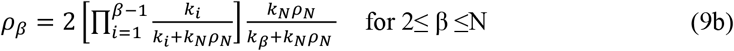

The average abundances of A and A* in a single cell {*a*_*β*_} and {*a*\*_*β*_}, respectively, approach steady states as well. The kinetics of these variables are given by (see **Supplementary Text 1**, section 5.2 for details),

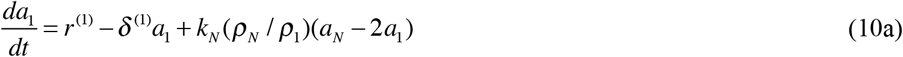

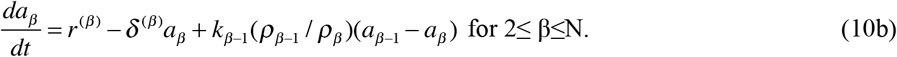

and,

For no heritability the equations for 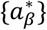 are (see **Supplementary Text 1**, section 5.3 for details),

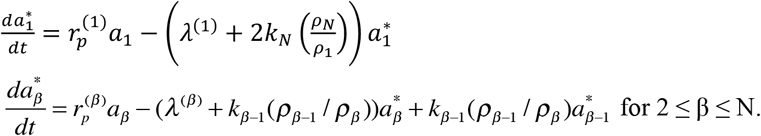

For full heritability for 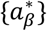 are,

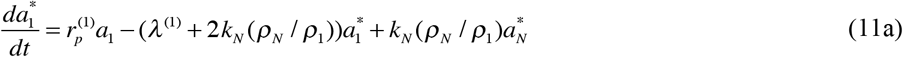

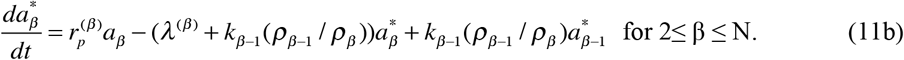

Here, 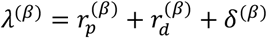 for 1≤ *β* ≤ N.

The steady state values of {*a*_β_} as functions of the scaled 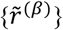 and dimensionless 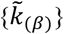 (i.e., ≤ 1) variables are given by (see **Supplementary Text 1**, section 5.2 for details),

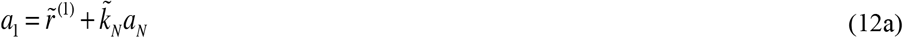

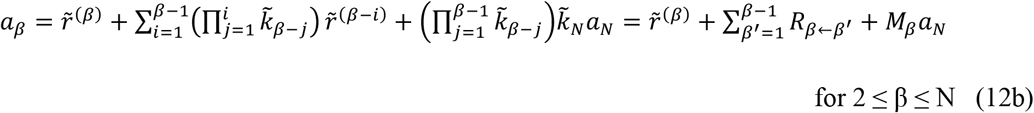

and,

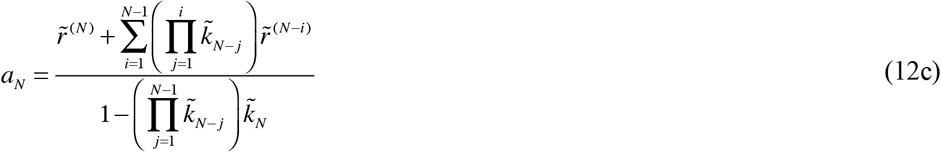

where,

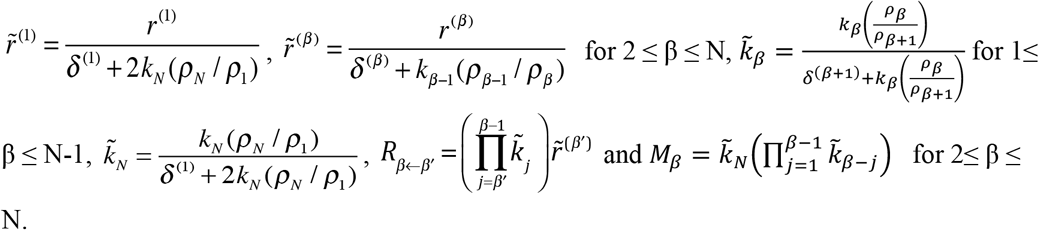

The steady state values for 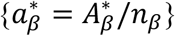 functions of dimensionless 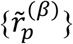 and 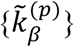 variables are given by (see **Supplementary Text 1**, section 5.3 for details),

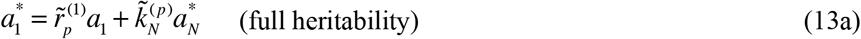

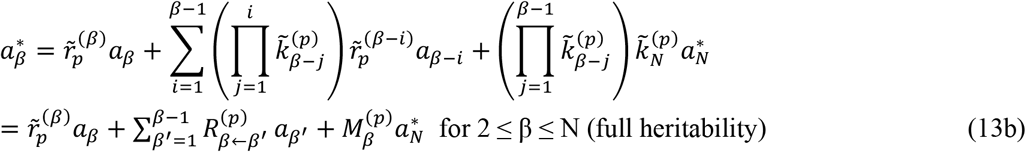

for 2 ≤ *β* ≤ N,

where,

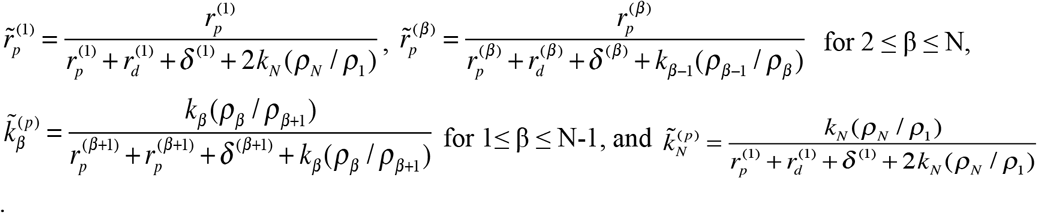

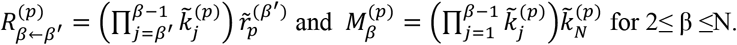

### Stochastic simulations

Multi-scale stochastic simulations were performed in MATLAB. The code can be found at https://github.com/indraniny/NK_cell_cycle_and_signaling, and a descriptive illustration is provided in **Supplementary Figure S8**. The simulation was seeded with 400 cells for the linear model (or 20 cells for the non-linear model), initialized with the abundances of each protein and a cell cycle stage designation. See “Initial Condition Generation” in Methods for details. Then, for each cell, a Gillespie simulation is performed^46^, where the cell can perform any protein-level reaction, or progress to the next stage of cell cycle. If the cell reaches the final stage (i.e. M stage) and advances, it divides, generating a second daughter cell. Proteins from the parent cell are randomly split between the two daughter cells according to a binomial distribution. Each cell also retains the “time” advanced from the previous timesteps in the Gillespie simulation for that cell. The process continues, with each cell independently selecting reactions to perform via the Gillespie simulation. When a cell is about to complete a reaction that would advance it past the endpoint time of interest, it is removed from the simulation and placed in the output. The simulation continues until all cells have been removed. The resulting output is a collection of cells that would be observed at the endpoint time of interest, originating from the input cells. Finally, we calculated the average behavior of protein abundances from the collections of cells obtained at the endpoint time for each cell cycle stage.

For non-linear simulations that require a volume correction (i.e., for a second-order reaction that governs binding of two proteins), the propensity for the reaction to occur decreases as the volume increases, which is correlated with progression through the cell cycle. To account for this, we assume that the propensity of the reaction to occur in each stage is given by:

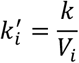

Where *V*_*i*_ = 1, 1.3, 1.6, 2.0 represents the volume factors in G1, S, G2, and M, respectively. This allows the volumes of each stage to take values between initial volume and twice the initial volume, mimicking a monotonic increase in cell volume up to a doubling of the cell volume. The simulations for the linear model, shown in **Figure 3**, took approximately one hour. The simulations for the non-linear model, which produced the results shown in **Figure 4**, had runtimes ranging from approximately 1 to 35 hours.

### Initial Condition Generation

For the non-linear model, we began with a population of 20 cells, each containing 80 molecules of the enzyme (E) and 80 molecules of the substrate (A) (**Supplementary Tables S1 and S2**). Initially, all cells were assumed to be in the G1 state of the cell cycle. Over time, these cells progress through the subsequent stages—S state, G2 state, and M state —before undergoing division to produce two daughter cells. The signaling pathway involves the binding of the enzyme (E) to the substrate (A), forming a complex (EA) that is further phosphorylated to create the product (A*), as described in **Figure 4A** (details in results section).We then let the system of reactions run for a long period (~ 5 generations of cells or 170 hours) until the mean protein concentrations of all species achieve steady state. At this point, we calculated the average protein abundances per cell at cell cycle stages G1, S, G2, and M. Because stage M has the shortest duration (~ 1.4 h), the number of cells in this stage is relatively low. In contrast, stage G1 has the longest duration, resulting in a higher number of cells (~14 h) (**Supplementary Table S1 and S2)**. Consequently, the fluctuation in average protein abundance is more pronounced during stage M, while it is less variable in the G1 state. We independently ran 20 simulation replicates of identical initial configurations to calculate the standard error of average protein abundance (SEM) in **Figures 4B-D**.

### Metric for Protein variation in sensitivity analysis

To quantify protein variation across cell cycle stages, we used the formula: (max−min)/min for fold change. Where max and min represent the average protein abundance per cell across the G1, S, G2, and M stages.

## Code Availability

The codes for this manuscript are written in MATLAB and Python, can be accessed https://github.com/indraniny/NK_cell_cycle_and_signaling.

## Acknowledgements

This work was supported by NIH grant R01AI146581 to J.D. and NIH S10 Instrumentation Grant S10 1S10OD018040-01 to L.L.L. We acknowledge to Cytobank software for performing CyTOF data analysis. We thank to HPC facility Franklin at the Nationwide Children’s Hospital for providing computational resources. We acknowledge Ohio Supercomputer Center for usage of MATLAB. BioRender is used to create the schematic figures.

## Author Contributions

Darren Wethington: Data processing; computational methods, investigation, data analysis; Figures; writing – original draft. Indrani Nayak: Data processing; computational methods, investigation, data analysis; Figures; writing – original draft. Helle Jensen: CyTOF experiments, Data curation, data analysis. Oscar A. Aguilar: Review. Gregory K. Behbehani: methodology, review. Lewis L. Lanier: Conceptualization, funding acquisition; investigation; methodology; project administration; supervision; writing – original draft, review and editing. Jayajit Das: Conceptualization, funding acquisition; investigation; methodology; project administration; supervision; Figures; writing – original draft, review and editing.

## Data Availability Statement

CyTOF data are available from the corresponding authors upon request, except for identifying donor information.

