## Supplementary Figures and Tables for "Natural Killer Cell Receptor Signaling and Activation Depend on Cell Cycle Stages"

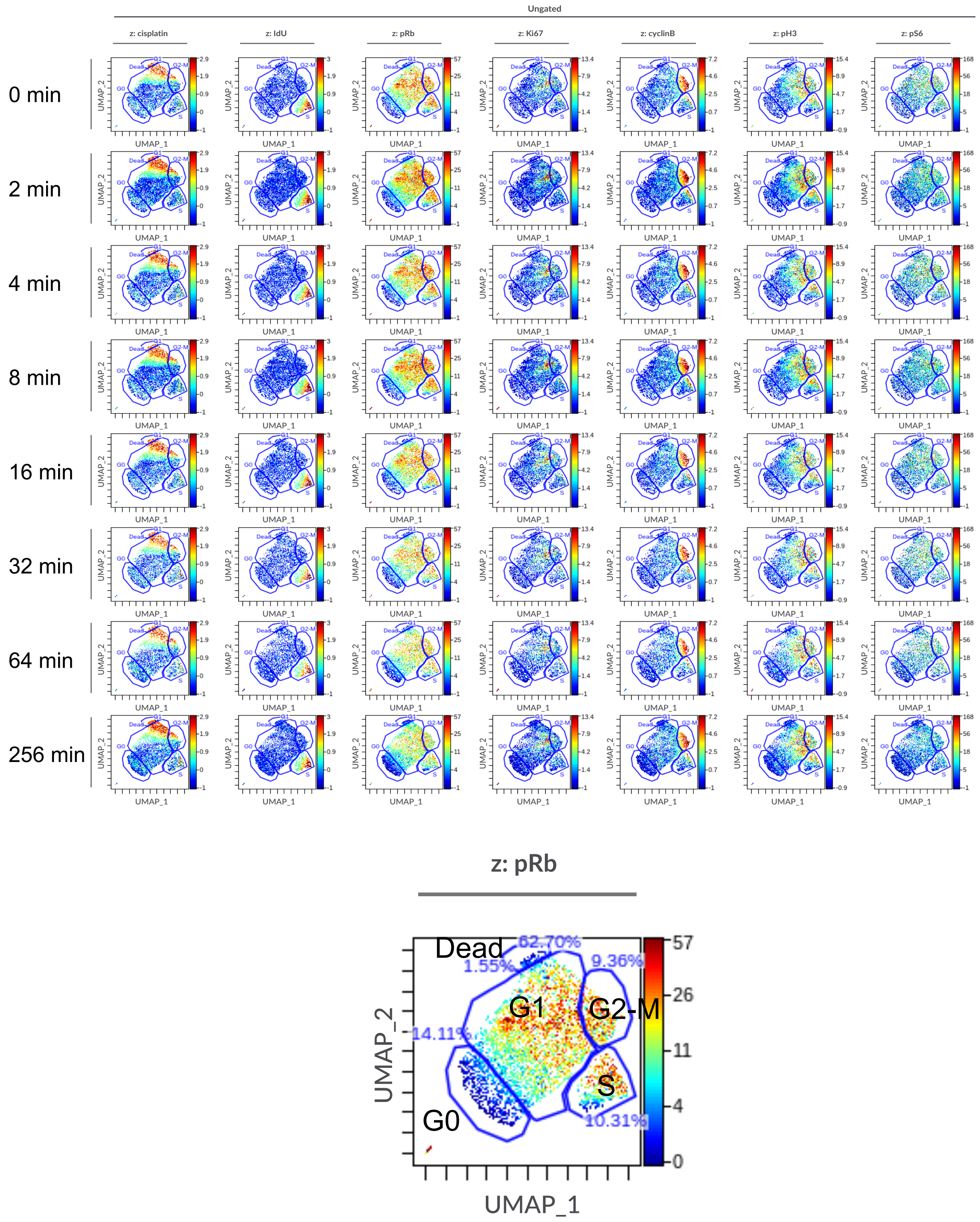


**Supplementary Figure S1: UMAP visualization of various markers across cell cycle stages for IL-2 treated NKG2D-stimulated primary NK cells.** CD56$^{dim}$ NK cell populations in dead, G0, S, and G2+M cell cycle stages were gated on UMAP using Cytobank software as described in

Materials and Methods. Data show the intensities for markers used for cell cycle gating as well as representative signaling proteins in the UMAP1-UMAP2 plane for unstimulated CD56$^{dim}$ NK cells (t = 0 min) and at 2, 4, 8, 16, 32, 64, and 256 min post-NKG2D antibody stimulation.

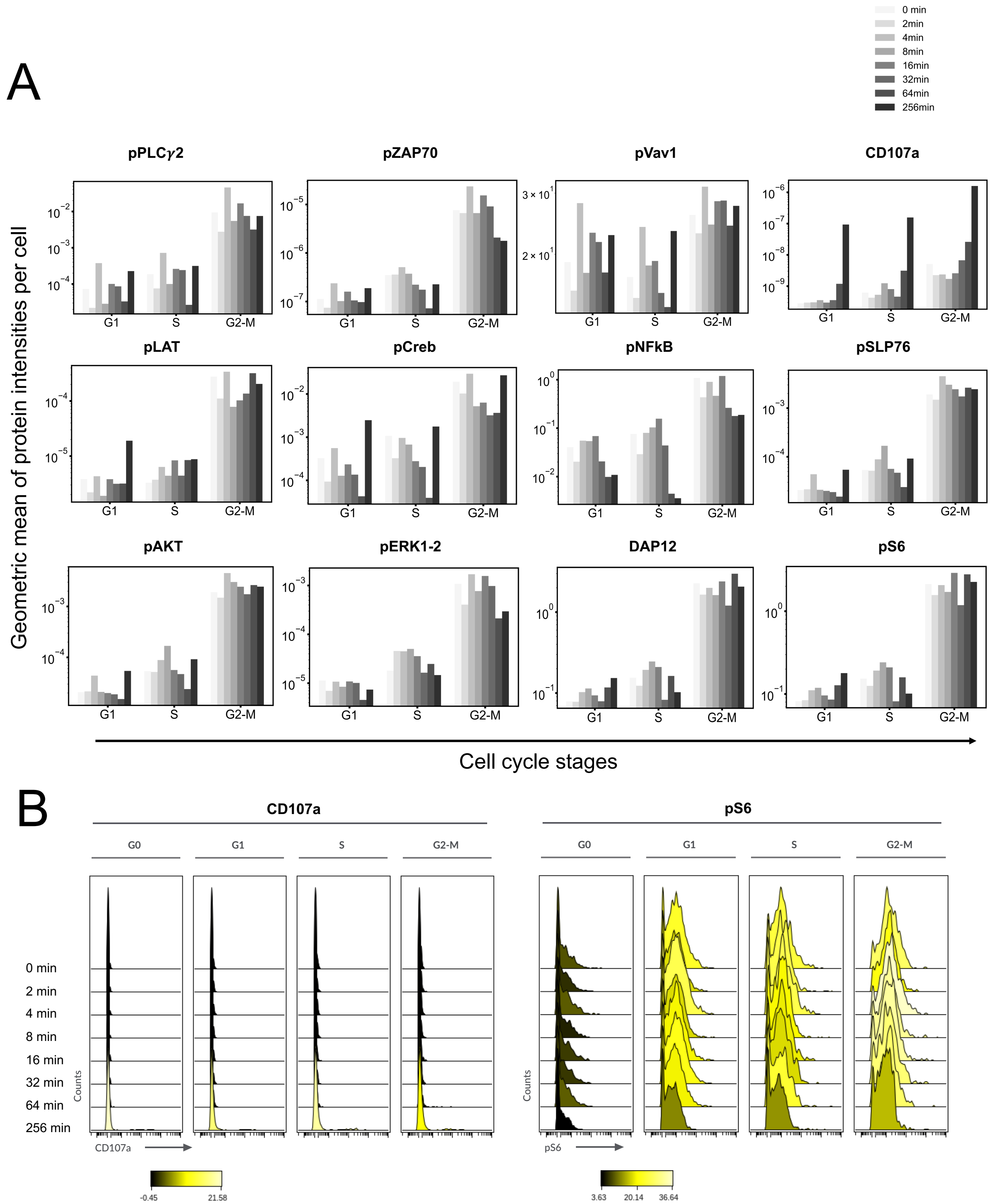

**Supplementary Figure S2: The mean intensities of various signaling proteins across cell cycle stages for IL2-treated NKG2D-stimulated primary NK cells. (A)** Geometric means for different proteins at multiple times post-NKG2D antibody stimulation is shown at cell cycle stages, G1, S, and G2+M. Majority of the phospho-proteins have low values of intensities except pVav1, pAKT and pS6. **(B)** Shows histograms of protein intensities for untransformed (raw) data at different time points across cell cycle stages. The colors of the histograms represent the arithmetic mean. The histograms were generated in Cytobank.

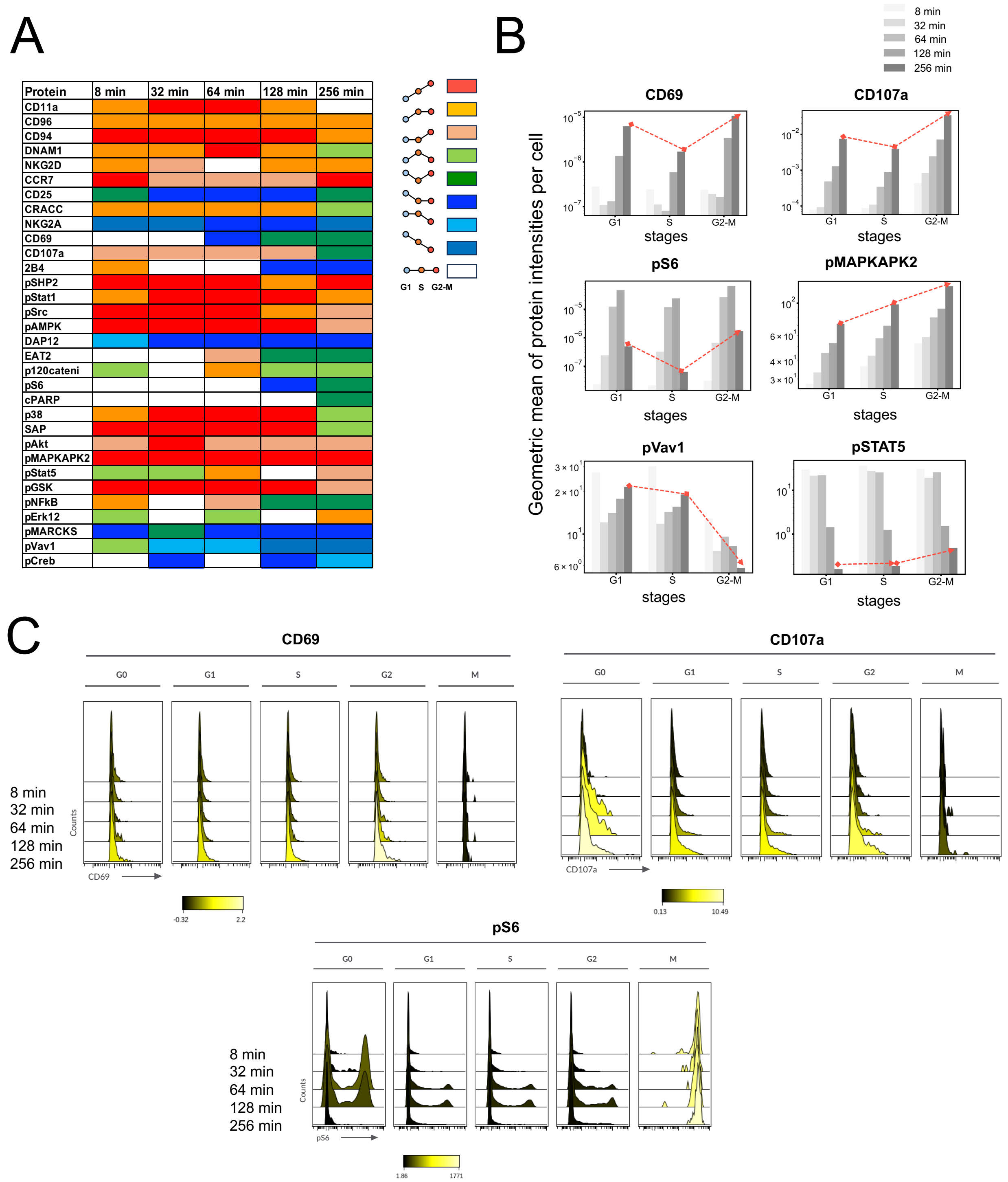


**Supplementary Figure S3: Variation of the abundances of signaling proteins in NKG2D-stimulated NKLs residing in different cell cycle stages.** The human NK cell line, NKL, constitutively expresses the activating human NKG2D receptor (see Materials and Methods for details). **(A)** Cell populations were gated from UMAPs into G1, S and G2+M for different time points using cell cycle marker pH3, Idu, cyclinB, pRb and cell viability marker cisplatin. The variations of the mean $\log_e$ (Intensity) of signaling proteins across cell cycle transitions, G1→ S and S → G2+M, are shown in terms of nine different types of variations. The nine types are characterized by the changes in (G1→ S, S → G2+M) as follows: (increase, increase), (increase, no-change), (no-change, increase), (increase, decrease), (decrease, increase), (decrease, no-change), (no-change, decrease), (decrease, decrease), (no-change, no-change). The variations are tabulated for time points 8, 32, 64, 128, and 256 min post-NKG2D antibody stimulation. The geometric means of the intensities are compared between cell cycle stages with one sided *t-test* (*P*-value < 0.05). **(B)** Geometric averages in intensities are shown as a function of cell cycle stages for different time points. **(C)** Shows histograms of protein intensities for untransformed (raw) data at different time points across cell cycle stages. The colors of the histograms represent the arithmetic mean. The histograms were generated in Cytobank.

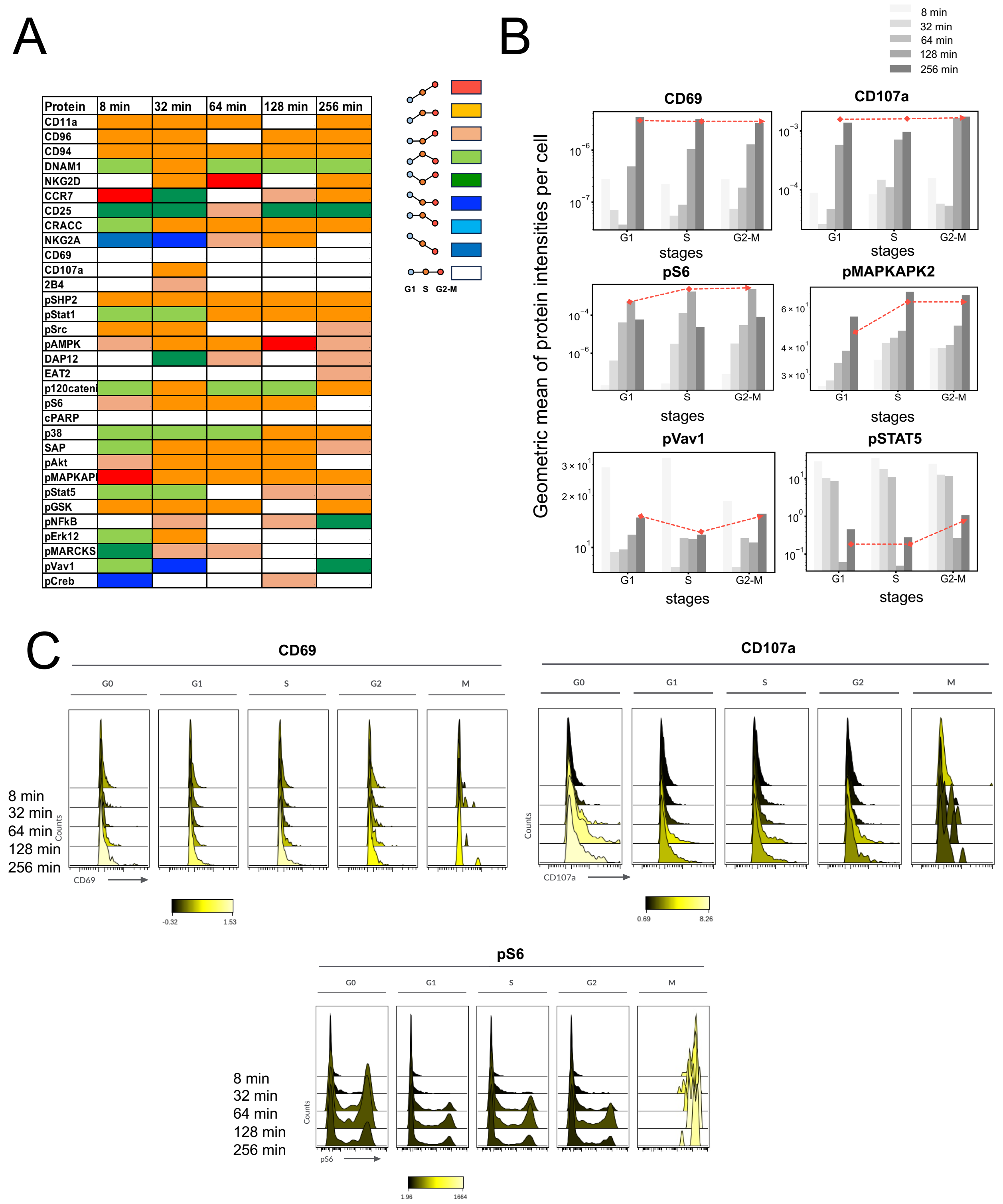


**Supplementary Figure S4: Variation of the abundances of signaling proteins in Ly49H-stimulated NKLs residing in different cell cycle stages.** The human NK cell line, NKL, was stably transduced to express the activating mouse Ly49H receptor (see Materials and Methods for details). **(A)** Cell populations were gated from UMAPs into G1, S and G2+M for different time points using cell cycle marker pH3, Idu, cyclinB, pRb and cell viability marker cisplatin. The variations of the mean $\log_e$ (Intensity) of signaling proteins across cell cycle transitions, G → S and S → G2+M, are shown in terms of nine different types of variations. The nine types are characterized by the changes in (G1→ S, S → G2+M) as follows: (increase, increase), (increase, no-change), (no-change, increase), (increase, decrease), (decrease, increase), (decrease, no-change), (no-change, decrease), (decrease, decrease), (no-change, no-change). The variations are tabulated for time points 8, 32, 64, 128, and 256 min post-Ly49H antibody stimulation. The geometric means of the intensities are compared between cell cycle stages with one sided *t-test* (*P*-value $< 0.05$). **(B)** Geometric averages in intensities are shown as a function of cell cycle stages for different time points. **(C)** Shows histograms of protein intensities for untransformed (raw) data at different time points across cell cycle stages. The colors of the histograms represent the arithmetic mean. The histograms were generated in Cytobank.

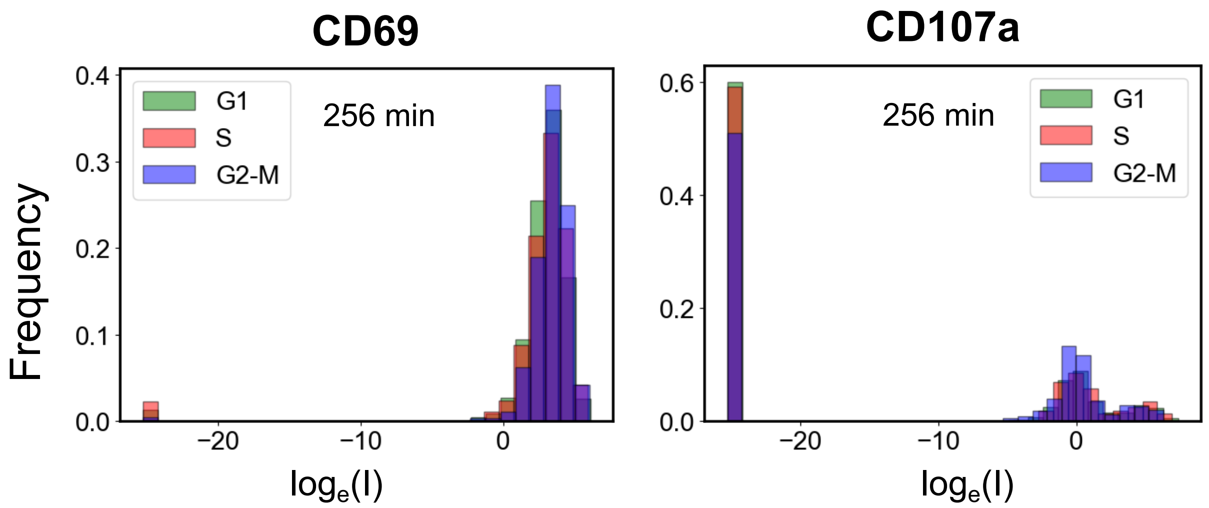


**Supplementary Figure S5: Normalized histograms of the natural log-transformed intensities (I) for the activation markers for IL2-treated NKG2D-stimulated primary NK cells.** (left) and (right) show the histograms for CD69 and CD107a in cell cycle stages G1, S and G2 at 256 min. (right) The double peak around 0 represents two distinct NK cell population having different expressions of CD107a. Population around the right peak (small height) indicates the NK cell population with higher degranulation.

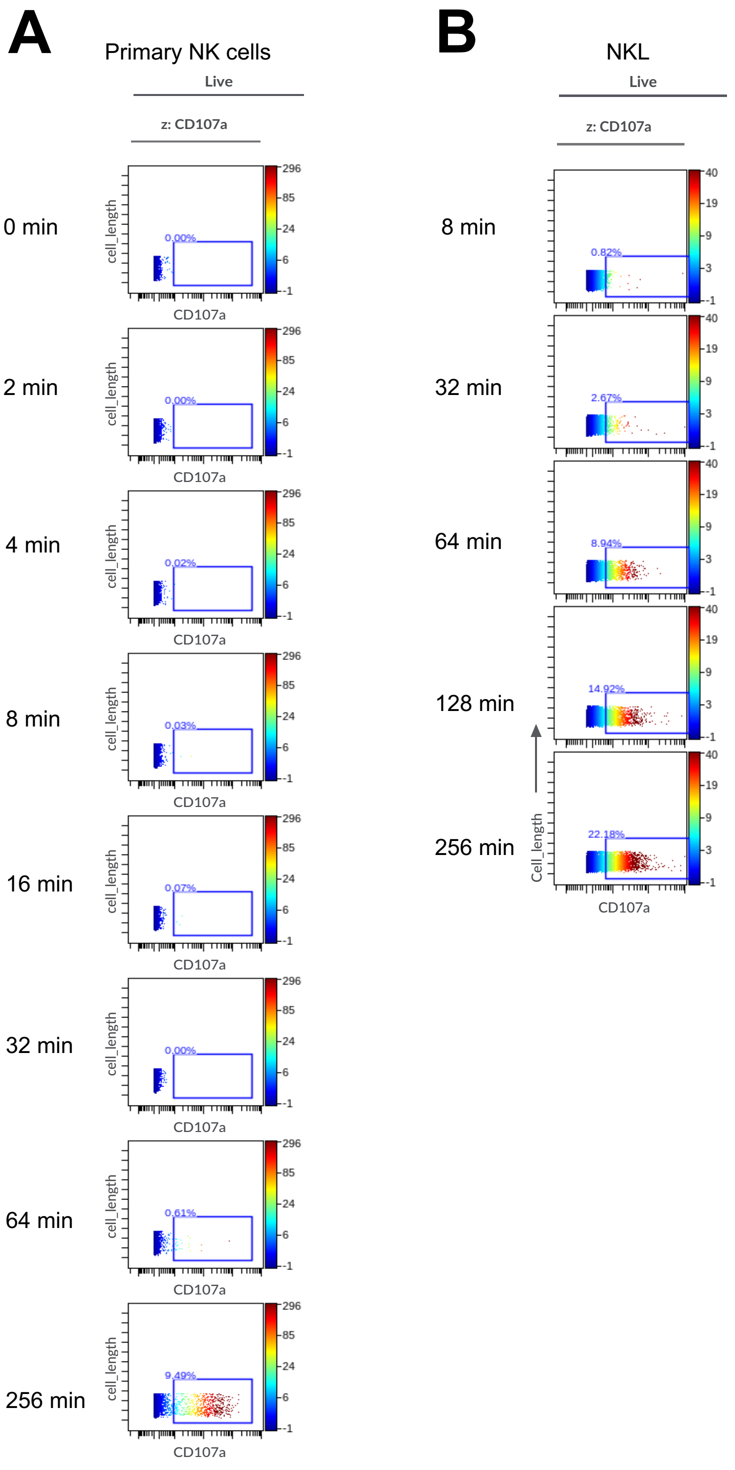


**Supplementary Figure S6: Degranulation in primary NK cells vs. NKL cells.** Dot plots depict high CD107a expressing cells following NKG2D stimulation in **(A)** primary NK cells and **(B)** NKL cells.

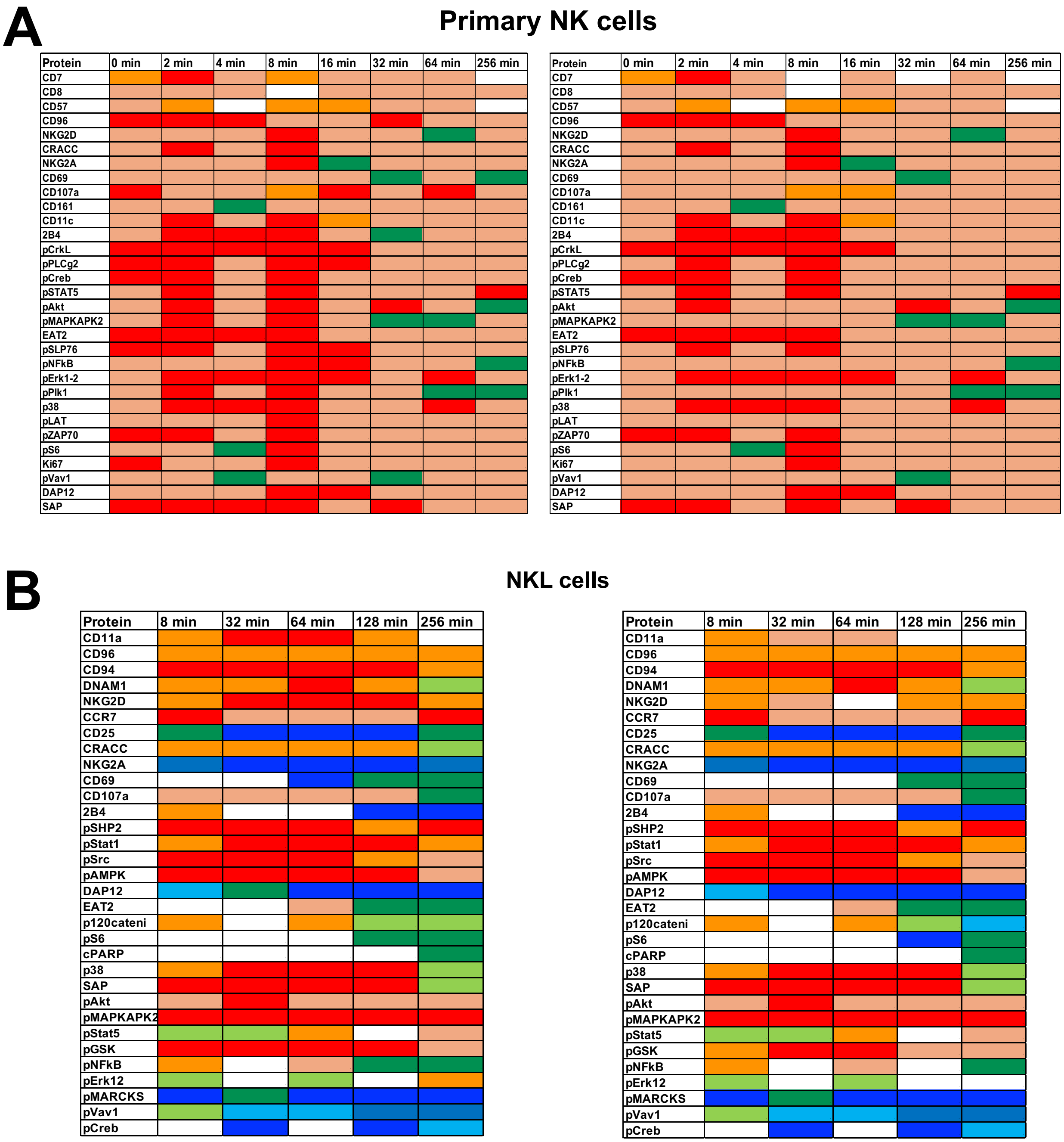


**Supplementary Figure S7: Sensitivity in variation of the abundances of signaling proteins in NKG2D-stimulated (A) primary NK cells and (B) NKLs residing in different cell cycle stages.** The analyses were performed varying the values of $\epsilon$ (see Data Processing section in Methods) and significance levels ($\alpha$) in the *t*-test of natural-log transformed protein intensities. **(A)** For primary NK cells, the left panel was generated considering $\epsilon = 10^{-7}$ and $\alpha = 0.05$, while the right panel was generated considering $\epsilon = 10^{-11}$ and $\alpha = 0.03$. **(B)** For NKL cells, the left panel was generated considering $\epsilon = 10^{-11}$ and $\alpha = 0.05$, while the right panel was generated considering $\epsilon = 10^{-11}$ and $\alpha = 0.03$.

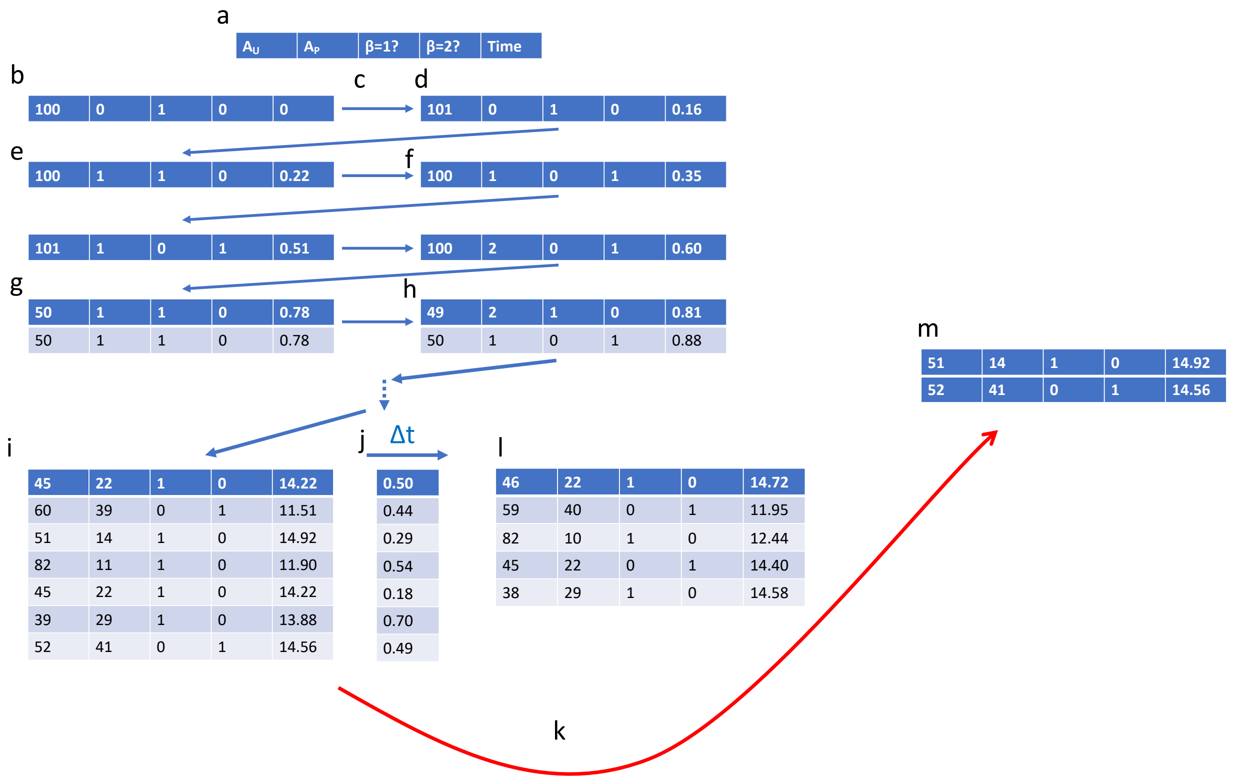


**Supplementary Figure S8: A descriptive illustration of multi-scale stochastic simulation.** For this representation, we consider the signaling system where a single protein $A$ is synthesized, degraded, and phosphorylated, while the phosphorylated form $A*$ is degraded and dephosphorylated. For this example, cells have only two cell cycle stages. We aim to profile the cell population at t=15. **(a)** Each column of the matrix used to represent single cells refers to 1) amount of unphosphorylated protein in the cell, 2) amount of phosphorylated protein in the cell, 3) a Boolean indicating whether the cell is in the first cell cycle stage, 4) a Boolean indicating whether the cell is in the second cell cycle stage. **(b)** The simulation is initialized with a single cell with 100 units of unphosphorylated protein in the first cell cycle stage. **(c)** Propensities for each possible reaction are evaluated, and a time step is randomly sampled according to the Gillespie algorithm. The time step Δt for this reaction is 0.16. **(d)** The randomly chosen reaction for the first step is synthesis of unphosphorylated $A$. This cell now has 101 units of $A$. **(e)** The next chosen reaction is phosphorylation of $A$, leading to one unit of $A*$ and 100 units of $A$. The Δt for this reaction was 0.06, leading to a ∑Δt of 0.22. **(f)** The next chosen reaction is progression to the second stage of the cell cycle. The Boolean values indicating which stage the cell is in are updated. Discrete reactions continue to occur. **(g)** This chosen reaction is division of the cell. A second row is added, indicating that there are now two cells in the simulation. Each cell is given $p$ and $n$-$p$ unphosphorylated and phosphorylated protein, respectively, according to binomial sampling. Both cells return to the first stage of the cell cycle and have progressed the same amount of time in the simulation (∑Δt=0.78). **(h)** We now evaluate a reaction for each cell in the simulation. The first has phosphorylation of protein, while the second advances to the second stage of the cell cycle. Δt is 0.03 and 0.10 for the first and second cell, respectively. The cells are now being simulated asynchronously. Many more discrete reactions occur, including cell divisions. **(i)** Many cells with varying protein expressions and ∑Δt values exist in the simulation. **(j)** Each cell is assigned a value of Δt as with each step before. The value ∑Δt must not cross the end time of t=15. The 3rd and last cell (rows) will cross t=15 with the given Δt values. **(k)** The two cells (3rd and last) have reached t=15 and are removed from the simulation. **(l)** The remaining cells continue to update according to the signaling and cell cycle processes until the reach t=15. When they do, they will be added to the group in **(m)**. This finalized group can then be analyzed.

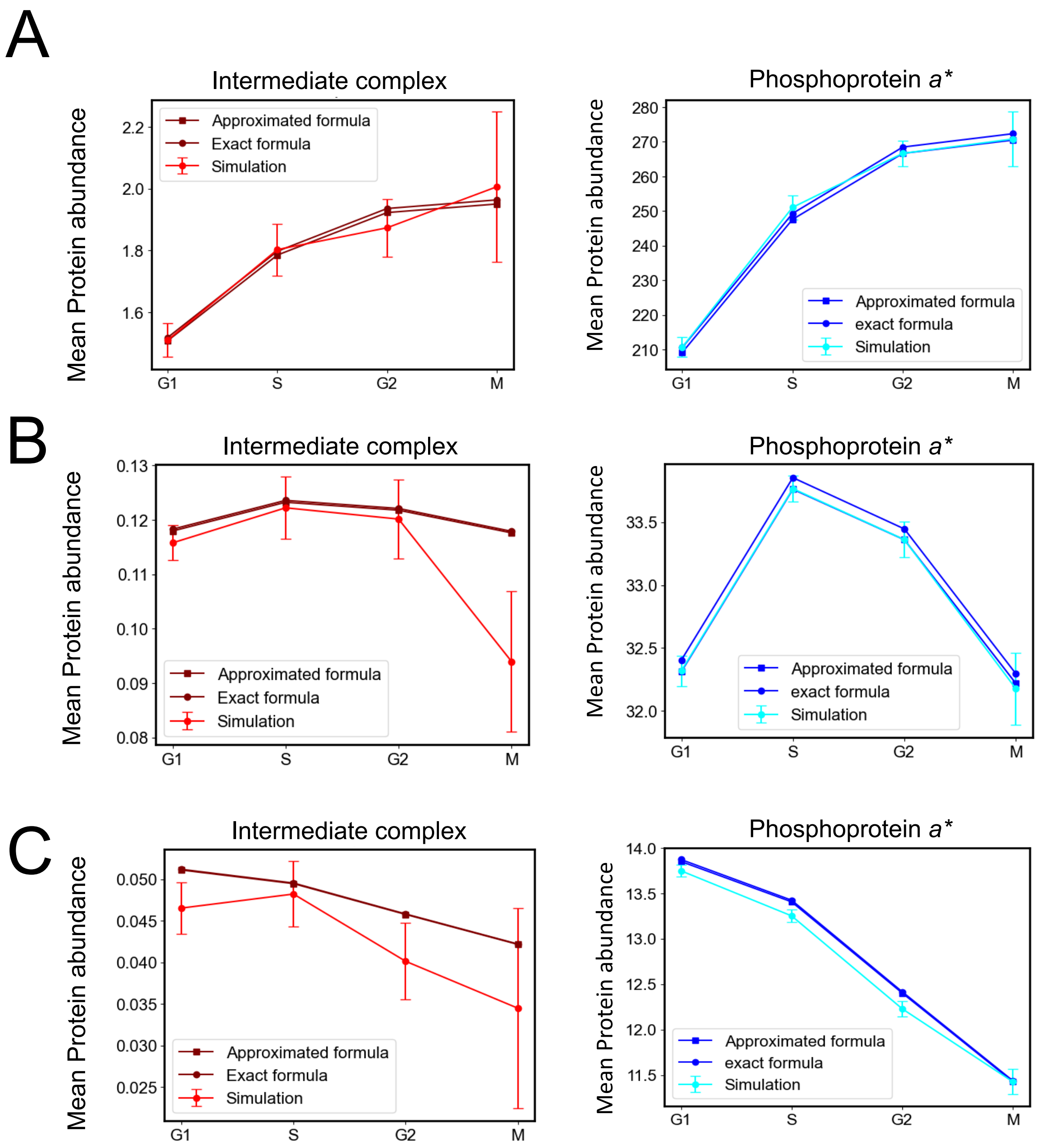


**Supplementary Figure S9: Comparison of the mean abundances of the intermediate complex and phosphoprotein (*a**) between *in silico* simulation, the approximated formula and exact formula for the non-linear protein model at steady state, related to Figure 4C in the main text.** The simulations in **(A-C)** show the abundances of the complex (red) and the phosphoprotein (cyan) obtained from Figures 4B, 4C (left) and 4C (right), respectively, in the main text. The approximated formulas in **(A-C)** show the abundances of the complex (maroon square) and the phosphoprotein (blue square), calculated using the approximated formulas (see Supplementary Text 1, Eqns. 92 and Eqn. 94). The exact formulas in **(A-C)** show the abundances of the complex (maroon circle) and the phosphoprotein (blue square), calculated using the exact formulas (see Supplementary Text 1, Eqns. 93 and Eqn. 94). For all exact and approximate formulas in **(A-C)**, we used steady-state values (at 170 hours) of abundances total A ($A^{S}_{tot}$ ) and total E ($E^{S}_{tot}$) for the G1, S, G2, and M phases from the simulation, along with the corresponding parameter values given in Supplementary Table 2.

**Supplementary Table S1: Parameter values used in the non-linear in-silico model (rates used here are from Table 1)**

| **Parameters used in simulation** | **Parameter values used in the simulation** | **Remarks** |
|---|---|---|
| Enzyme (E) | 80 molecules | Assumption. The cytosolic volume ($V_{cyto}$) of NK cell is ~ 80 um$^3$ assuming cell and nucleus radii to be 3 um and 2 ums, respectively. For simulation, we assumed a fraction of cytosolic cell volume as a cell to reduce computational cost. |
| Substrate (A) | 80 molecules | Assumption. The cytosolic volume of NK cell. $V_{cyto}$ is ~ 80 um$^3$ assuming cell and nucleus radii to be 3 um and 2 ums, respectively. For simulation, we assumed a fraction of cytosolic cell volume as a cell to reduce computational cost. |
| Cell volume ($V_e$) | 0.132 um$^3$ | Since inside the cell the protein density is ~1 uM or 602 molecules per um$^3$ (e.g., ZAP70 density in NK cell), the simulation volume ($V_e$) to accommodate 80 copies is ~ 0.132 um$^3$. |
| Cell cycle transition rate | [0.07, 0.1, 0.2, 0.7] h$^{-1}$ | For primary NK cell, the cell cycle takes around 30 hrs. How the time durations are distributed into G1, S, G2 and M are not known. We have calculated the cell cycle transition rates in primary NK cell using the proportion of duration of cell cycle stages in Hela cell. |

| | | |
|---|---|---|
| Simulation time | 150 h -170 h | Assuming avg. protein per cell will reach to a steady state value after 170 hours or ~ 5 complete cycles. |
| Enzyme or Substrate synthesis rate (r) | 18-5350 molecules $h^{-1}$ | This is a zeroth order reaction rule. Because we have considered a fraction of cell volume for simulation, we scaled the realistic synthesis rate (10800-3240000 copies per hour) from literature by a multiplicative factor ($V_e/V_{cyto}$). The calculation of multiplicative factor is shown in Supplementary Text **1,** section 10. |
| Protein degradation rate ($\delta$) | 0.03-0.9 $h^{-1}$ | This is a first-order reaction. Thus, it remains same for any cell volume. |
| Complex formation rate ($k_{on}$) | 4.53-4530 $h^{-1}$ per molecule | This is a second order reaction rule. The association range could be $10^5$-$10^8$ $M^{-1}$ $s^{-1}$ . For example, ZAP70 binds to ITAM with rate 5 $uM^{-1}$ $s^{-1}$ = 29.90 $\mu m^3$ $h^{-1}$ per copy. Because we have considered a fraction of cell volume for simulation, we scaled the realistic $k_{on}$ from literature by a multiplicative factor (1/ $V_e$) to calculate the propensity which gives 226 $h^{-1}$ per copy. The calculation of multiplicative factor is shown in Supplementary Text **1,** section 10. |
| Complex dissociation rate ($k_{off}$) | 450-7200 $h^{-1}$ | In T cell, ZAP70 unbinding rate from ITAM 0.125 $s^{-1}$. SHP1 unbinding rate from ITAM 2 $s^{-1}$. |
| Protein phosphorylation rate ($r_p$) | 36-7200000 $h^{-1}$ | This is a first-order reaction. Thus, it remains same for any cell volume. |

| | | |
|---|---|---|
| Protein dephosphorylation rate ($r_d$) | 36-7200000 $h^{-1}$ | This is a first-order reaction. Thus, it remains same for any cell volume. |

**Supplementary Table S2A: Parameter values used in Figure 4**

| Figure index | r (molecules $h^{-1}$) | δ ($h^{-1}$) | $k_{on}$ ($h^{-1}$ per molecule) | $k_{off}$ ($h^{-1}$) | $r_p$ ($h^{-1}$) | $r_d$ ($h^{-1}$) | Volume factor at G1, S, G2, M |
|---|---|---|---|---|---|---|---|
| **B** | 18 | 0.05 | 15 | 3000 | 5000 | 36 | [1, 1.3, 1.6, 2] |
| **C (left)** | 20 | 0.5 | 10 | 3000 | 10000 | 36 | [1, 1.3, 1.6, 2] |
| **C (right)** | 20 | 0.9 | 5 | 5000 | 10000 | 36 | [1, 1.3, 1.6, 2] |
| **D (left)** | 18 | 0.05 | 15 | 3000 | [5000, 50, 5000, 5000] | 36 | [1, 1.3, 1.6, 2] |
| **D (right)** | 18 | 0.05 | 15 | 3000 | [5000, 5000, 50, 50] | 36 | [1, 1.3, 1.6, 2] |

**Other common parameters for Figures 4B-4D**

| Enzyme (molecules per cell) | Substrate (molecules per cell) | Cell transition rates ($h^{-1}$) | Simulation time (h) | Cell volume ($\mu m^3$ ) |
|---|---|---|---|---|
| 80 | 80 | [0.07, 0.1, 0.2, 0.7] | 150-170 | 0.132 |

**Supplementary Table S2B: *P*-values are tabulated for *t*-tests to compare the avg. protein abundances between cell cycle stages in Figure 4**

| | G1-S | S-G2 | G2-M |
|---|---|---|---|
| **Fig. B** | $1.14\times10^{-31}$ | $2.32\times10^{-16}$ | 0.04 |
| **Fig. C (left)** | $3.21\times10^{-17}$ | $2.19\times10^{-6}$ | $8.20\times10^{-6}$ |
| **Fig. C (right)** | $5.73\times10^{-10}$ | $2.87\times10^{-14}$ | $6.04\times10^{-9}$ |
| **Fig. D (left)** | $1.37\times10^{-48}$ | $1.97\times10^{-45}$ | $3.04\times10^{-4}$ |
| **Fig. D (right)** | $5.26\times10^{-34}$ | $1.78\times10^{-47}$ | $2.2\times10^{-8}$ |

**Supplementary Table S2C: Parameter ranges used to obtain protein variation for sensitivity analysis**

| r (molecules $h^{-1}$) | δ ($h^{-1}$) | $k_{on}$ ($h^{-1}$ per molecule) | $k_{off}$ ($h^{-1}$) | $r_p$ ($h^{-1}$) | $r_d$ ($h^{-1}$) | Volume factor at G1, S, G2, M |
|---|---|---|---|---|---|---|
| 20 | [0.03,0.9] | [5,100] | 3000 | 10000 | 36 | [1, 1.3, 1.6, 2] |

*parameter values are chosen randomly within the range of δ and $k_{on}$.

**Supplementary Table S3: Mechanism of protein abundances across cell cycle stages in silico model for realistic parameter ranges given in Supplementary Table S1**

Assuming cell cycle transition rates $k_\beta$ depends on the cell cycle stage $\beta$. See the approximated calculation in Supplementary Text **1,** section 11.

| Protein Behavior G1→ S → G2 | *Neutral* Case with const. signaling rates | Condition | Protein behavior along G1 →S →G2 | | | |
|---|---|---|---|---|---|---|
| | | | $E_{tot}$ | $A_{tot}$ | C | P |
| Monotonic increase (increase, increase) | $r, \delta, k_{on}, k_{off}, r_p, r_d$ | $\delta < k_\beta$ | Monotonic rise | Monotonic rise | Monotonic rise | Monotonic rise |
| Non-monotonic (increase, decrease) | | $x$ small, Competing $(f_1 \times f_2)$ Where $f_1, f_2$ are increasing and decreasing functions, respectively. | Monotonic rise | Monotonic rise | Non-monotonic peak | Non-monotonic peak |
| Monotonic decrease (decrease, decrease) | | $x$ small, $\delta \gg k_\beta$ Competing $(f_1 \times f_2)$ Where $f_1, f_2$ are constant and decreasing functions, respectively. | No change | No change | Monotonic decay | Monotonic decay |
| Unchanged (no-change, no-change) | | $\delta \gg k_\beta$ $K_D' \ll 1$ | No change | No change | No change | No change |

$$K'_D = \frac{K_{off}+\delta+r_p}{K_{on}}, E_{tot} = A_{tot} = \frac{r}{\delta} = \text{total concentrations of E and A}$$

$$f_1 = E_{tot} \times A_{tot}, \quad f_1 \text{ is an increasing or constant function with cell cycle stages.}$$

$$f_2 = \frac{1}{A_{tot}+\beta E_{tot}+K'_D}, \quad f_2 \text{ is a decreasing function as cell volume grows with cell cycle stages.}$$

$$\beta = 1 + \frac{r_p}{\delta+r_d}, \text{ range } [1, \infty]$$

$$x = 4\beta f_1 f_2^2$$

**Supplementary Table S4: Mass cytometry Ab panel used for analysis of primary NK cells**

| | Antibody marker | Metal | Ab Clone | Vendor |
|---|---|---|---|---|
| Surface | CD7 | 113In | CD7-6B7 | BioLegend |
| | CD45 | 115In | HI30 | BioLegend |
| | CD57 | 139La | HCD57 | BioLegend |
| | CD3+CD235+CD61 +CD15 | 141Pr | UCHT1+HIR2+VI-PL2 +W6D3 | BioLegend |
| | CD96 | 142Nd | NK92.39 | BioLegend |
| | CD8 | 147Sm | RTA-T8 | BioLegend |
| | CD16 | 148Nd | 3G8 | Fluidigm |
| | CD107a | 151Eu | H4A3 | Fluidigm |
| | NKG2D | 154Sm | 1D11 | BioLegend |
| | CRACC | 155Gd | 162.1 | BioLegend |
| | CD161 | 158Gd | 191B8 | Beckman Coulter |
| | CD11c | 159Tb | Bu15 | Fluidigm |
| | 2B4 | 160Gb | PP35 | eBioscience |
| | CD69 | 162Dy | FN50 | Fluidigm |
| | NKG2A | 174Yb | Z199 | Beckman Coulter |
| | CD56 | 176Yb | NCAM16.2 | BD |
| Intracellular | pCrkL(Y207) | 143Nd | Polyclonal | Fluidigm |
| | pHistone H3(S28) | 145Nd | HTA28 | BioLegend |
| | pPLCγ2(Y759) | 146Nd | K86-689.37 | BD |
| | pCreb(S133) | 149Sm | 87G3 | Cell Signaling |
| | pSTAT5(Y694) | 150Nd | 47 | BD |
| | pAkt(S473) | 152Sm | D9E | Fluidigm |
| | pMAPKAPK2(T334) | 153Eu | 27B7 | Cell Signaling |
| | Cyclin B | 156Gd | GNS-1 | BD |
| | SAP | 157Gd | XLP-1D12 | Cell Signaling |
| | DAP12 | 161Dy | 406288 | R&D Systems |
| | EAT2 | 163Dy | Polyclonal | Abgent |
| | pSLP76(Y128) | 164Dy | J141-668.36.58 | BD |
| | pNFκB(S529) | 165Ho | K10-895.12.50 | BD |
| | pRb(S807/S811) | 166Er | J112-906 | BD |
| | pErk1/2(T202/Y204) | 167Er | D13-14-4E | Fluidigm |
| | pPlk1(T210) | 168Er | K50-483 | BD |
| | p38(T180/Y182) | 169Tm | 36/p38 | BD |
| | pLAT(Y226) | 170Er | J96-1238.58.93 | BD |
| | pZAP70(Y319/Y352) | 171Yb | 17a | Fluidigm |

| | | | | |
|---|---|---|---|---|
| | pS6(S235/S236) | 172Yb | N7-548 | Fluidigm |
| | Ki67 | 173Yb | SolA15 | eBioscience |
| | pVav1(Y160) | 175Lu | Polyclonal | Life Technology |

**Supplementary Table S5: Mass cytometry Ab panel used for analysis of NKL cells**

| | Antibody marker | Metal | Ab Clone | Vendor |
|---|---|---|---|---|
| Surface | CD45 | 115In | HI30 | BioLegend |
| | CD11a | 139La | HI111 | BD |
| | CD96 | 142Nd | NK92.39 | BioLegend |
| | CD94 | 146Nd | DX22 | BioLegend |
| | DNAM1 | 149Sm | 11A8 | BioLegend |
| | CD107a | 151Eu | H4A3 | Fluidigm |
| | NKG2D | 154Sm | 1D11 | BioLegend |
| | CRACC | 155Gd | 162.1 | BioLegend |
| | CCR7 | 159Tb | 150503 | R&D Systems |
| | 2B4 | 160Gb | PP35 | eBioscience |
| | CD69 | 162Dy | FN50 | Fluidigm |
| | CD25 | 169Tm | BC96 | BioLegend |
| | NKG2A | 174Yb | Z199 | Beckman Coulter |
| Intracellular | pSHP2(Y580) | 141Pr | polycloncal | Cell Signaling |
| | pSTAT1(Y701) | 143Nd | 4a | BD |
| | pSrc(Y418) | 144Nd | K98-37 | BD |
| | pAMPK(T172) | 145Nd | 40H9 | Cell Signaling |
| | pGSK-3a/b(S21/9) | 147Sm | 37F11 | Cell Signaling |
| | pSTAT5(Y694) | 150Nd | 47 | BD |
| | pAkt(S473) | 152Sm | D9E | Fluidigm |
| | pMAPKAPK2(T334) | 153Eu | 27B7 | Cell Signaling |
| | p38(T180/Y182) | 156Gd | 36/p38 | BD |
| | SAP | 157Gd | XLP-1D12 | Cell Signaling |
| | Ki67 | 158Gd | SolA15 | eBioscience |
| | DAP12 | 161Dy | 406288 | R&D Systems |
| | EAT2 | 163Dy | Polyclonal | Abgent |
| | Cyclin B | 164Dy | GNS-1 | BD |
| | pNFκB(S529) | 165Ho | K10-895.12.50 | BD |
| | pRb(S807/S811) | 166Er | J112-906 | BD |
| | pErk1/2(T202/Y204) | 167Er | D13-14-4E | Fluidigm |
| | pHistone H3(S28) | 168Er | HTA28 | BioLegend |
| | pMARCKS(S152/56) | 170Er | polyclonal | Cell Signaling |
| | p120 catenin(T916) | 171Yb | 1/catenin | BD |
| | pS6(S235/S236) | 172Yb | N7-548 | Fluidigm |
| | cPARP | 173Yb | F21-852 | BD |
| | pVav1(Y160) | 175Lu | Polyclonal | Life Technology |

| | | | | |
|---|---|---|---|---|
| | pCreb(S133) | 176Yb | 87G3 | Cell Signaling |
