## Supplementary Text 1 for "Natural Killer Cell Receptor Signaling and Activation Depend on Cell Cycle Stages"

February 12, 2025

Consider a simple model where single cells can be in $N$ cell cycle states: $\beta = 1, .., N$ where, the rate of transition from one cell cycle state to next cell cycle stage $\beta \to \beta + 1$ is denoted by $k_\beta$. We have considered a simple model where within each cell, a protein $A$ can be synthesized ($\phi \to A$) with rate $r$ and degraded ($A \to \phi$) with rate $\delta$. Protein $A$ can further be phosphorylated ($A \to A^*$) with phosphorylation rate $r_p$. The phosphorylated protein $A^*$ can be dephosphorylated ($A^* \to A$) by rate $r_d$. For these linear signaling reaction model to calculate the average protein behavior $(A, A^*)$, we wrote down the Master equation as follows:

$$\frac{dP(\{..., n_{a,a*,\beta}, ...\}, t)}{dt} = (\text{contributions from cell cycle transitions} + \text{contributions from signaling reactions} + \text{contribution from protein synthesis-degradation}) \quad (1)$$

Where, $P(\{..., n_{a,a*,\beta}, ...\}, t)$ represents the probability of getting number of cells $n$ at cell cycle stage $\beta$, having $a$ number of protein ($A$) and $a^*$ number of phosphorylated protein ($A^*$) at time $t$. In the following, we have written down the individual contributions from cell cycle transitions, signaling reactions and protein synthesis-degradation (RHS of Eq. (1)).

#### 1 Contributions from cell cycle transitions

##### Differentiation or $\beta \to \beta + 1$ $(1 \le \beta \le (N-1))$ transition :

Here, cell is transitioning from one cell cycle stage ($\beta$) to the next cell cycle stage ($\beta + 1$)

$$\begin{aligned} T_\beta^{cell\ cycle} &= \sum_{a=0}^{\infty} \sum_{a^*=0}^{a} k_\beta (n_{a,a^*,\beta} + 1) P(\{..., n_{a,a^*,\beta} + 1, n_{a,a^*,\beta+1} - 1, ...\}, t) \\ &- \sum_{a=0}^{\infty} \sum_{a^*=0}^{a} k_\beta (n_{a,a^*,\beta}) P(\{..., n_{a,a^*,\beta}, ...\}, t) \end{aligned} \quad (2)$$

##### Division or $N \to 1$ transition (no heritability for $A^*$) :

Mother cell ($\beta = N$) is dividing into the two daughter cells ($\beta = 1$). No phosphorylated protein ($A^*$) of mother cells is transferred to the daughter cells upon cell division.

$$\begin{aligned} T_N^{cell\ cycle} &= \sum_{a=0}^{\infty} \sum_{a^*=0}^{a} \sum_{m=0}^{a} k_N (n_{a,a^*,N} + 1) (^aC_m 2^{-a}) P(\{..., n_{a,a^*,N} + 1, n_{m,0,1} - 1, n_{a-m,0,1} - 1, ...\}, t) \\ &- \sum_{a=0}^{\infty} \sum_{a^*=0}^{a} \sum_{m=0}^{a} k_N (n_{a,a^*,N}) (^aC_m 2^{-a}) P(\{..., n_{a,a^*,N}, n_{m,0,1}, n_{a-m,0,1} ...\}, t) \end{aligned} \quad (3)$$

##### or, division or $N \to 1$ transition (full heritability for $A^*$):

Mother cell is dividing into the two daughter cells. All phosphorylated proteins of mother cells are transferred to the daughter cells upon cell division. The phosphorylated $A^*$ molecules in a cell at state $N$ are partitioned with a binomial probability distribution into the daughter cells. For example, if a cell in state $N$ with $a$ and $a^*$ molecules produce daughter cells with $m$ and $a-m$ number of $A$ molecules, the $a^*$ molecules can be partitioned as, $(0,a^*),(1,a^*-1),(2,a^*-2),(m^*,a^*-m^*),..,(a^*,0)$ when $m \geq a^*$ and, $(0,a^*),(1,a^*-1),(2,a^*-2),(m^*,a^*-m^*),..,(m,a^*-m)$ when $m < a^*$. The $1^{st}$ and the $2^{nd}$ place in the parenthesis refers to the number of $A^*$ in the two daughter cells having $m$ and $(a-m)$ number of $A$ molecules. The probability of choosing any $(m^*,a^*-m^*)$ is proportional to ${}^{a^*}C_{m^*}(1/2)^{m^*}(1/2)^{a^*-m^*}$, where $m^*=1,...,a^*$ for $m >= a^*$ and $m^*=1,...,m$ for $m < a*$.

$$
\begin{aligned}
T_N^{cell\ cycle} = &\sum_{a=0}^{\infty}\sum_{a^*=0}^{a}\sum_{m=0}^{a}\sum_{m^*=0}^{\tilde{a}^*} k_N(n_{a,a^*,N}+1)({}^{a}C_m 2^{-a})({}^{a^*}C_{m^*}2^{-a^*}) \\
&P(\{...,n_{a,a^*,N}+1,n_{m,m^*,1}-1,n_{a-m,a^*-m^*,1}-1,...\},t) \\
&-\sum_{a=0}^{\infty}\sum_{a^*=0}^{a}\sum_{m=0}^{a}\sum_{m^*=0}^{\tilde{a}^*} k_N(n_{a,a^*,N})({}^{a}C_m 2^{-a})({}^{a^*}C_{m^*}2^{-a^*})P(\{...,n_{a,a^*,N},n_{m,m^*,1},n_{a-m,a^*-m^*,1}...\},t)
\end{aligned} \tag{4}
$$

$\tilde{a}^* = a^*$ when $m \geq a^*$, and $\tilde{a}^* = m$ when $m < a^*$.

### 2 Contributions from signaling reactions

#### Phosphorylation ($A \rightarrow A^*$; rate $r_p$) at state$-\beta$:

At cell cycle stage $\beta$, a protein ($A$) is phosphorylating with rate $r_p$ within a cell.

$$
\begin{aligned}
T_\beta^{phospho} = &\sum_{a=0}^{\infty}\sum_{a^*=0}^{a} r_p(a-(a^*-1)(n_{a,a^*-1,\beta}+1)P(\{...,n_{a,a^*-1,\beta}+1,n_{a,a^*,\beta}-1,...\},t) \\
&-\sum_{a=0}^{\infty}\sum_{a^*=0}^{a} r_p(a-a^*)(n_{a,a*,\beta})P(\{...,n_{a,a^*-1,\beta},n_{a,a^*,\beta},...\},t)
\end{aligned} \tag{5}
$$

#### Dephosphorylation ($A^* \rightarrow A$; rate $r_d$) at state$-\beta$:

At cell cycle stage $\beta$, a phospho-protein ($A^*$) is de-phosphorylating with rate $r_d$ within a cell.

$$
\begin{aligned}
T_\beta^{dephospho} = &\sum_{a=0}^{\infty}\sum_{a^*=0}^{a} r_d(a^*+1)(n_{a,a^*+1,\beta}+1)P(\{...,n_{a,a^*+1,\beta}+1,n_{a,a^*,\beta}-1,...\},t) \\
&-\sum_{a=0}^{\infty}\sum_{a^*=0}^{a} r_d(a^*)(n_{a,a^*,\beta})P(\{...,n_{a,a^*+1,\beta},n_{a,a^*,\beta},...\},t)
\end{aligned} \tag{6}
$$

### 3 Contributions from synthesis-degradation of a signaling protein

At cell cycle stage $\beta$, a protein ($A$) is produced with zeroth-order reaction rate $r$ within a cell.

#### Production of A ( $\phi \rightarrow A$ ; rate $r$) at state$-\beta$:

$$
\begin{aligned}
T_\beta^{synthesis} = &\sum_{a=0}^{\infty}\sum_{a^*=0}^{a} r(n_{a-1,a^*,\beta}+1)P(\{...,n_{a-1,a^*,\beta}+1,n_{a,a^*,\beta}-1,...\},t) \\
&-\sum_{a=0}^{\infty}\sum_{a^*=0}^{a} r(n_{a,a^*,\beta})P(\{...,n_{a-1,a^*,\beta},n_{a,a^*,\beta},...\},t)
\end{aligned} \tag{7}
$$

##### Degradation of unphosphorylated-protein $A^U$ ($A^U \to \phi$ ; rate $\delta$) at state$-\beta$:

At cell cycle stage $\beta$, an unphosphorylated-protein ($A^U$) is degraded with first-order reaction rate $\delta$ within a cell.

$$\begin{aligned} T_{\beta}^{degradation} &= \sum_{a=0}^{\infty}\sum_{a^*=0}^{a} \delta(a+1-a^*)(n_{a+1,a^*,\beta}+1)P(\{...,n_{a+1,a^*,\beta}+1, n_{a,a^*,\beta}-1,...\},t) \\ &- \sum_{a=0}^{\infty}\sum_{a^*=0}^{a} \delta(a-a^*)(n_{a,a^*,\beta})P(\{...,n_{a+1,a^*,\beta}, n_{a,a^*,\beta},...\},t) \end{aligned} \tag{8}$$

##### Degradation of phospho-protein $A^*$ ($A^* \to \phi$ ; rate $\delta$) at state$-\beta$:

$$\begin{aligned} T_{\beta}^{degradation} &= \sum_{a=0}^{\infty}\sum_{a^*=0}^{a} \delta(a^*+1)(n_{a+1,a^*+1,\beta}+1)P(\{...,n_{a+1,a^*+1,\beta}+1, n_{a,a^*,\beta}-1,...\},t) \\ &- \sum_{a=0}^{\infty}\sum_{a^*=0}^{a} \delta a^*(n_{a,a^*,\beta})P(\{...,n_{a+1,a^*+1,\beta}, n_{a,a^*,\beta},...\},t) \end{aligned} \tag{9}$$

#### 4 Calculating average cell number and protein abundances

The average number of single cells at state -$\beta$ is,

$$n_{\beta} = \sum_{a=0}^{\infty}\sum_{a^*=0}^{a}\sum_{\{n_{a,a^*,\beta}\}} n_{a,a^*,\beta}P(\{...,n_{a,a^*,\beta},...\},t) \tag{10}$$

The average number of total protein abundance $A$ in $n$ number of single cells at state $\beta$ is,

$$A_{\beta} = \sum_{a=0}^{\infty}\sum_{a^*=0}^{a}\sum_{\{n_{a,a^*,\beta}\}} (a n_{a,a^*,\beta})P(\{...,n_{a,a^*,\beta},...\},t) \tag{11}$$

The average number of total phospho-protein abundance $A^*$ in $n$ number of single cells at state $\beta$ is,

$$A^*_{\beta} = \sum_{a=0}^{\infty}\sum_{a^*=0}^{a}\sum_{\{n_{a,a^*,\beta}\}} (a^* n_{a,a^*,\beta})P(\{...,n_{a,a^*,\beta},...\},t) \tag{12}$$

Where $\sum_{\{n_{a,a^*,\beta}\}}$ represents the sum over various cell populations.

##### Time evolution for the mean observables :

The time-dependent Ordinary differential Equations (ODE) of average number of single cells at state -$\beta$ is,

$$\frac{dn_{\beta}}{dt} = \sum_{a=0}^{\infty}\sum_{a^*=0}^{a}\sum_{\{n_{a,a^*,\beta}\}} (n_{a,a^*,\beta})\frac{dP(\{...,n_{a,a^*,\beta},...\},t)}{dt} \tag{13}$$

The time-dependent ODE of the average number of total protein abundance $A$ in $n$ number of single cells at state $\beta$ is,

$$\frac{dA_{\beta}}{dt} = \sum_{a=0}^{\infty}\sum_{a^*=0}^{a}\sum_{\{n_{a,a^*,\beta}\}} (a n_{a,a^*,\beta})\frac{dP(\{...,n_{a,a^*,\beta},...\},t)}{dt} \tag{14}$$

The time-dependent ODE of the average number of total phospho-protein abundance $A^*$ in $n$ number of single cells at state $\beta$ is,

$$\frac{dA^*_{\beta}}{dt} = \sum_{a=0}^{\infty}\sum_{a^*=0}^{a}\sum_{\{n_{a,a^*,\beta}\}} a^*(n_{a,a^*,\beta})\frac{dP(\{...,n_{a,a^*,\beta},...\},t)}{dt} \tag{15}$$

where for Eq.(13)-(15),

$$\frac{dP(\{...,n_{a,a^*,\beta},...\},t)}{dt} = \sum_{\beta=1}^{N}[T_\beta^{cell\ cycle} + T_\beta^{signalling} + T_\beta^{synthesis} + T_\beta^{degradation}] \tag{16}$$

Where, $T_\beta^{signalling} = [T_\beta^{phospho} + T_\beta^{dephospho}]$

#### 4.1 Calculating RHS from possible reactions for Eq.(13)-(15) :

##### 4.1.1 Contribution in $\frac{dn_1}{dt}$ from cell cycle state transition from state $1 \to 2$

$$\begin{aligned}\frac{dn_1}{dt} &= \sum_{a=0}^{\infty}\sum_{a^*=0}^{a}\sum_{\{n_{a,a^*,\beta}\}}(n_{a,a^*,1})\frac{dP(\{...,n_{a,a^*,1},...\},t)}{dt} = \sum_{a=0}^{\infty}\sum_{a^*=0}^{a}\sum_{\{n_{a,a^*,\beta}\}}(n_{a,a^*,1})\big[T_1^{cell\ cycle}\big] \\ &= \sum_{a=0}^{\infty}\sum_{a^*=0}^{a}\sum_{\{n_{a,a^*,\beta}\}}(n_{a,a^*,1})\big[\sum_{a'=0}^{\infty}\sum_{a'^*=0}^{a'}k_1(n_{a',a'^*,1}+1)P(\{...,n_{a',a'^*,1}+1,n_{a',a'^*,2}-1,...\},t) \\ &\qquad - \sum_{a'=0}^{\infty}\sum_{a'^*=0}^{a'}k_1(n_{a',a'^*,1})P(\{...,n_{a',a'^*,1},...\},t)\big]\end{aligned} \tag{17}$$

Defining
$n_{a',a'^*,1}+1 = n'_{a',a'^*,1}$ and $n_{a',a'^*,2}-1 = n'_{a',a'^*,2}$ the above sum becomes

$$\begin{aligned}(I_1 - I_2) = k_1\sum_{a=0}^{\infty}\sum_{a^*=0}^{a}\sum_{\{n_{a,a^*,\beta}\}}\sum_{a'=0}^{\infty}\sum_{a'^*=0}^{a'}&\big[(n_{a,a^*,1})(n'_{a',a'^*,1})P(\{...,n'_{a',a'^*,1},n'_{a',a'^*,2},...\},t) \\ &-(n_{a,a^*,1})(n_{a',a'^*,1})P(\{...,n_{a',a'^*,1},...\},t)\big]\end{aligned} \tag{18}$$

For $a = a'$ and $a^* = a'^*$ :

$$\begin{aligned}(I_1 - I_2) &= k_1\sum_{a'=0}^{\infty}\sum_{a'^*=0}^{a'}\sum_{\{n_{a',a'^*,\beta}\}}\big[(n_{a',a'^*,1})(n'_{a',a'^*,1})P(\{...,n'_{a',a'^*,1},n'_{a',a'^*,2},...\},t) \\ &\qquad -(n_{a',a'^*,1})(n_{a',a'^*,1})P(\{...,n_{a',a'^*,1},...\},t)\big] \\ &= k_1\sum_{a'=0}^{\infty}\sum_{a'^*=0}^{a'}\sum_{\{n_{a,a^*,\beta}\}}\big[(n'_{a',a'^*,1}-1)(n'_{a',a'^*,1})P(\{...,n'_{a',a'^*,1},n'_{a',a'^*,2},...\},t) \\ &\qquad -(n_{a',a'^*,1})(n_{a',a'^*,1})P(\{...,n_{a',a'^*,1},...\},t)\big] = -k_1 n_1\end{aligned} \tag{19}$$

Where $n_1 = \sum_{a'=0}^{\infty}\sum_{a'^*=0}^{a'}\sum_{\{n_{a',a'^*,\beta}\}}(n'_{a',a'^*,1})P(\{...,n'_{a',a'^*,1},n'_{a',a'^*,2},...\},t)$
For $a \neq a'$ and $a^* \neq a'^*$, the terms of $I_1$ and $I_2$ cancel each other.

Therefore, the contribution of differentiation to state $1 \to 2$ is $-k_1 n_1$ for $\frac{dn_1}{dt}$.
Similarly, the terms corresponding to $\beta \to \beta+1$ will give $-k_\beta n_\beta$ for $\frac{dn_\beta}{dt}$.

##### 4.1.2 Contribution in $\frac{dn_1}{dt}$ from cell cycle state transition from state $N \to 1$:

We have derived here the contribution for the no heritability case, as both this case and the full heritability case will generate identical contributions. The contribution of $T_N^{cell\ cycle}$ to $\frac{dn_1}{dt}$ can be described in terms of the sums below:

$$\begin{aligned}(I_1 - I_2) = k_N\sum_{a=0}^{\infty}\sum_{a^*=0}^{a}\sum_{\{n_{a,a^*,\beta}\}}\sum_{a'=0}^{\infty}\sum_{a'^*=0}^{a'}\sum_{m=0}^{a'}& \\ \big[(n_{a,a^*,1})(n_{a',a'^*,N}+1)({}^{a'}C_m 2^{-a'})P(\{...,n_{a',a'^*,N}+1,n_{m,0,1}-1,n_{a'-m,0,1}-1,...\},t)& \\ -(n_{a,a^*,1})(n_{a',a'^*,N})({}^{a'}C_m 2^{-a'})P(\{...,n_{a',a'^*,N},n_{m,0,1},n_{a'-m,0,1}...\},t)\big]&\end{aligned} \tag{20}$$

Defining
$n_{a',a'^*,N}+1=n'_{a',a'^*,N}$,
$n_{m,0,1}-1=n'_{m,0,1}$
$n_{a'-m,0,1}-1=n'_{a'-m,0,1}$
Substituting above we get,

$$\begin{split}(I_1-I_2)=k_N\sum_{a=0}^{\infty}\sum_{a^*=0}^{a}\sum_{\{n_{a,a^*,\beta}\}}\sum_{a'=0}^{\infty}\sum_{a'^*=0}^{a'}\sum_{m=0}^{a'}\\ \big[(n_{a,a^*,1})(n'_{a',a'^*,N})({}^{a'}C_m2^{-a'})P(\{...,n'_{a',a'^*,N},n'_{m,0,1},n'_{a'-m,0,1},...\},t)\\ -(n_{a,a^*,1})(n_{a',a'^*,N})({}^{a'}C_m2^{-a'})P(\{...,n_{a',a'^*,N},n_{m,0,1},n_{a'-m,0,1}...\},t)\big]\end{split} \tag{21}$$

Expanding $(I_1-I_2)$ for $a=m,a^*=0$ and $a=(a'-m),a^*=0$ we get :

$$\begin{split}(I_1-I_2)=k_N\sum_{a'=0}^{\infty}\sum_{a'^*=0}^{a'}\sum_{m=0}^{a'}\sum_{\{n_{a',a'^*,\beta}\}}\big[(n_{m,0,1}+n_{a'-m,0,1})(n'_{a',a'^*,N})({}^{a'}C_m2^{-a'})P(\{...,n'_{a',a'^*,N},n'_{m,0,1},n'_{a'-m,0,1},...\},t)\\ -(n_{m,0,1}+n_{a'-m,0,1})(n_{a',a'^*,N})({}^{a'}C_m2^{-a'})P(\{...,n_{a',a'^*,N},n_{m,0,1},n_{a'-m,0,1}...\},t)\big]\\ =k_N\sum_{a'=0}^{\infty}\sum_{a'^*=0}^{a'}\sum_{m=0}^{a'}\sum_{\{n_{a',a'^*,\beta}\}}({}^{a'}C_m2^{-a'})\big[(n'_{m,0,1}+1+n'_{a'-m,0,1}+1)(n'_{a',a'^*,N})P(\{...,n'_{a',a'^*,N},n'_{m,0,1},n'_{a'-m,0,1},...\},t)\\ -(n_{m,0,1}+n_{a'-m,0,1})(n_{a',a'^*,N})P(\{...,n_{a',a'^*,N},n_{m,0,1},n_{a'-m,0,1}...\},t)\big]\\ =2k_N\sum_{a'=0}^{\infty}\sum_{a'^*=0}^{a'}\sum_{\{n_{a',a'^*,\beta}\}}[\sum_{m=0}^{a'}({}^{a'}C_m2^{-a'})](n'_{a',a'^*,N})P(\{...,n'_{a',a'^*,N},n'_{m,0,1},n'_{a'-m,0,1},...\},t)\\ =2k_N\sum_{a'=0}^{\infty}\sum_{a'^*=0}^{a'}(2^{a'}2^{-a'})]\overline{(n'_{a',a'^*,N})}=2k_Nn_N\end{split} \tag{22}$$

For $a\neq m,a^*\neq 0$ and $a\neq(a'-m),a^*\neq 0$, the terms of $I_1$ and $I_2$ cancel each other.

Therefore, the contribution of $N\to 1$ to $\frac{dn_1}{dt}$ is $2k_Nn_N$.

##### 4.1.3 Contribution in $\frac{dA_1}{dt}$ from cell cycle state transition from state $1\to 2$:

The contribution in $\frac{dA_1}{dt}$ from the term $T_1^{cell\ cycle}$ is,

$$\begin{split}\frac{dA_1}{dt}=\sum_{a=0}^{\infty}\sum_{a^*=0}^{a}\sum_{\{n_{a,a^*,\beta}\}}(an_{a,a^*,1})\frac{dP(\{...,n_{a,a^*,1},n_{a,a^*,2},...\},t)}{dt}\\ =\sum_{a=0}^{\infty}\sum_{a^*=0}^{a}\sum_{\{n_{a,a^*,\beta}\}}(an_{a,a^*,1})\Big[\sum_{a'=0}^{\infty}\sum_{a'^*=0}^{a'}k_1(n_{a',a'^*,1}+1)P(\{...,n_{a',a'^*,1}+1,n_{a',a'^*,2}-1,...\},t)\\ -\sum_{a'=0}^{\infty}\sum_{a'^*=0}^{a'}k_1(n_{a',a'^*,1})P(\{...,n_{a',a'^*,1},n_{a',a'^*,2},...\},t)\Big]\end{split} \tag{23}$$

Defining
$n_{a',a'^*,1}+1=n'_{a',a'^*,1}$
$n_{a',a'^*,2}-1=n'_{a',a'^*,2}$

$$\begin{split}=k_1\sum_{a=0}^{\infty}\sum_{a^*=0}^{a}\sum_{\{n_{a,a^*,\beta}\}}\sum_{a'=0}^{\infty}\sum_{a'^*=0}^{a'}\big[(an_{a,a^*,1})(n'_{a',a'^*,1})P(\{...,n'_{a',a'^*,1},n'_{a',a'^*,2},...\},t)\\ -(an_{a,a^*,1})(n_{a',a'^*,1})P(\{...,n_{a',a'^*,1},n_{a',a'^*,2},...\},t)\big]=(I_1-I_2)\end{split} \tag{24}$$

For $a \neq a', a^* \neq a'^*$, the terms of $I_1$ and $I_2$ cancel each other.
Expanding $(I_1 - I_2)$ for $a = a'$ and $a^* = a'^*$ we get :

$$\begin{aligned}
&= k_1 \sum_{a'=0}^{\infty} \sum_{a'^*=0}^{a'} \sum_{\{n_{a',a'^*,\beta}\}} \big[(a' n_{a',a'^*,1})(n'_{a',a'^*,1}) P(\{..., n'_{a',a'^*,1}, n'_{a',a'^*,2}, ...\}, t) \\
&\qquad - (a' n_{a',a'^*,1})(n_{a',a'^*,1}) P(\{..., n_{a',a'^*,1}, n_{a',a'^*,2}, ...\}, t)\big] \\
&= k_1 \sum_{a'=0}^{\infty} \sum_{a'^*=0}^{a'} \sum_{\{n_{a',a'^*,\beta}\}} \big[(a'[n'_{a',a'^*,1} - 1])(n'_{a',a'^*,1}) P(\{..., n'_{a',a'^*,1}, n'_{a',a'^*,2}, ...\}, t) \\
&\qquad - (a' n_{a',a'^*,1})(n_{a',a'^*,1}) P(\{..., n_{a',a'^*,1}, n_{a',a'^*,2}, ...\}, t)\big] \\
&= -k_1 \sum_{a'=0}^{\infty} \sum_{a'^*=0}^{a'} \sum_{\{n_{a',a'^*,\beta}\}} \big[(a' n'_{a',a'^*,1}) P(\{..., n'_{a',a'^*,1}, n'_{a',a'^*,2}, ...\}, t) \\
&= -k_1 \sum_{a'=0}^{\infty} \sum_{a'^*=0}^{a'} [(a' \overline{n'_{a',a'^*,1}}) = -k_1 A_1
\end{aligned} \tag{25}$$

Similarly, contribution of a transition $\beta \to \beta + 1$ to $\frac{dA_\beta}{dt}$ and $\frac{dA_{\beta+1}}{dt}$ are $-k_\beta A_\beta$ and $+k_\beta A_\beta$, respectively.

##### 4.1.4 Contribution in $\frac{dA_1}{dt}$ from cell cycle state transition from state $N \to 1$:

The contribution in $\frac{dA_1}{dt}$ from the term $T_N^{cell\ cycle}$ in no heritability case is,

$$\begin{aligned}
\frac{dA_1}{dt} &= \sum_{a=0}^{\infty} \sum_{a^*=0}^{a} \sum_{\{n_{a,a^*,\beta}\}} (a n_{a,a^*,1}) \frac{dP(\{..., n_{a,a^*,N}, n_{m,0,1}, n_{a-m,0,1}, ...\}, t)}{dt} \\
&= \sum_{a=0}^{\infty} \sum_{a^*=0}^{a} \sum_{\{n_{a,a^*,\beta}\}} (a n_{a,a^*,1}) k_N \sum_{a'=0}^{\infty} \sum_{a'^*=0}^{a} \sum_{m=0}^{a'} ({}^{a'}C_m 2^{-a'}) \big[(n_{a',a'^*,N} + 1) \\
&\qquad P(\{..., n_{a',a'^*,N} + 1, n_{m,0,1} - 1, n_{a'-m,0,1} - 1, ...\}, t) \\
&\qquad - (n_{a',a'^*,N}) P(\{..., n_{a',a'^*,N}, n_{m,0,1}, n_{a'-m,0,1} ...\}, t)\big] = (I_1 - I_2)
\end{aligned} \tag{26}$$

Defining
$n_{a',a'^*,N} + 1 = n'_{a',a'^*,N}$,
$n_{m,0,1} - 1 = n'_{m,0,1}$
$n_{a'-m,0,1} - 1 = n'_{a'-m,0,1}$
we get

$$\begin{aligned}
(I_1 - I_2) = \sum_{a=0}^{\infty} \sum_{a^*=0}^{a} \sum_{\{n_{a,a^*,\beta}\}} \sum_{a'=0}^{\infty} \sum_{a'^*=0}^{a'} \sum_{m=0}^{a'} (a n_{a,a^*,1}) k_N ({}^{a'}C_m 2^{-a'}) \big[(n'_{a',a'^*,N}) P(\{..., n'_{a',a'^*,N}, n'_{m,0,1}, n'_{a'-m,0,1}, ...\}, t) \\
- (n_{a',a'^*,N}) P(\{..., n_{a',a'^*,N}, n_{m,0,1}, n_{a'-m,0,1} ...\}, t)\big]
\end{aligned} \tag{27}$$

For $a \neq m, a^* \neq 0$ and $a \neq (a' - m), a^* \neq 0$, the terms of $I_1$ and $I_2$ cancel each other.

Expanding $(I_1 - I_2)$ for $a = m, a^* = 0$ and $a = (a' - m), a^* = 0$ we get :

$$
\begin{aligned}
&(I_1 - I_2) \\
&= k_N \sum_{a'=0}^{\infty} \sum_{a'^*=0}^{a'} \sum_{\{n_{a',a'^*,\beta}\}} \sum_{m=0}^{a'} ({}^{a'}C_m 2^{-a'}) \big[(mn_{m,0,1} + (a'-m)n_{a'-m,0,1})(n'_{a',a'^*,N})P(\{..., n'_{a',a'^*,N}, n'_{m,0,1}, n'_{a'-m,0,1}, ...\}, t) \\
&\qquad - (mn_{m,0,1} + (a'-m)n_{a'-m,0,1})(n_{a',a'^*,N})P(\{..., n_{a',a'^*,N}, n_{m,0,1}, n_{a'-m,0,1}...\}, t)\big] \\
&= k_N \sum_{a'=0}^{\infty} \sum_{a'^*=0}^{a'} \sum_{\{n_{a',a'^*,\beta}\}} \big[(m(n'_{m,0,1} + 1) + (a'-m)(n'_{a'-m,0,1} + 1)](n'_{a',a'^*,N})P(\{..., n'_{a',a'^*,N}, n'_{m,0,1}, n'_{a'-m,0,1}, ...\}, t) \\
&\qquad - (mn_{m,0,1} + (a'-m)n_{a'-m,0,1})(n_{a',a'^*,N})P(\{..., n_{a',a'^*,N}, n_{m,0,1}, n_{a'-m,0,1}...\}, t)\big] \\
&= k_N \sum_{a'=0}^{\infty} \sum_{a'^*=0}^{a} \sum_{\{n_{a',a'^*,\beta}\}} a'(n'_{a',a'^*,N})P(\{..., n'_{a',a'^*,N}, n'_{m,0,1}, n'_{a'-m,0,1}, ...\}, t) = k_N \sum_{a'=0}^{\infty} \sum_{a'^*=0}^{a} a'\overline{(n'_{a',a'^*,N})} = k_N A_N
\end{aligned} \tag{28}
$$

Therefore, the contribution of $T_N^{cell\ cycle}$ in the kinetics for $A_1$ is $k_N A_N$ . Full heritability for $A^*$ The contribution $T_N^{cell\ cycle}$ is the same as that for the no heritability case.

###### 4.1.5 Contribution of protein production $\phi \to A$ in $\frac{dA_1}{dt}$ at state-$\beta$:

$$
\begin{aligned}
&\frac{dA_1}{dt} = \sum_{a=0}^{\infty} \sum_{a^*=0}^{a} \sum_{\{n_{a,a^*,\beta}\}} (an_{a,a^*,1}) \frac{dP(\{..., n_{a,a^*,1}, ...\}, t)}{dt} \\
&= \sum_{a=0}^{\infty} \sum_{a^*=0}^{a} \sum_{\{n_{a,a^*,\beta}\}} a(n_{a,a^*,1}) \big[ \sum_{a'=0}^{\infty} \sum_{a'^*=0}^{a'} r(n_{a'-1,a'^*,1} + 1)P(\{..., n_{a'-1,a'^*,1} + 1, n_{a',a'^*,1} - 1, ...\}, t) \\
&\qquad - \sum_{a'=0}^{\infty} \sum_{a'^*=0}^{a'} r(n_{a',a'^*,1})P(\{..., n_{a'-1,a'^*,1}, n_{a',a'^*,1}, ...\}, t)\big] = (I_1 - I_2)
\end{aligned} \tag{29}
$$

Defining
$n_{a'-1,a'^*,1} + 1 = n'_{a'-1,a'^*,1}$,
$n_{a',a'^*,1} - 1 = n'_{a',a'^*,1}$

we get

$$
\begin{aligned}
(I_1 - I_2) = r \sum_{a=0}^{\infty} \sum_{a^*=0}^{a} \sum_{\{n_{a,a^*,\beta}\}} \sum_{a'=0}^{\infty} \sum_{a'^*=0}^{a'} &\big[(an_{a,a^*,1})(n'_{a'-1,a'^*,1})P(\{..., n'_{a'-1,a'^*,1}, n'_{a',a'^*,1}, ...\}, t) \\
&- (an_{a,a^*,1})(n_{a',a'^*,1})P(\{..., n_{a'-1,a'^*,1}, n_{a',a'^*,1}, ...\}, t)\big]
\end{aligned} \tag{30}
$$

Expanding $I_1$ in the Eqn. (30) for $a = (a' - 1), a = a'$ and $a \neq (a' - 1)$ or $a'$, we get the following

$$\begin{aligned}
I_1 = \big[r \sum_{\{n_{a',a'^*,\beta}\}} \sum_{a'=0}^{\infty} \sum_{a'^*=0}^{a'} (a'-1)(n_{a'-1,a'^*,1})(n'_{a'-1,a'^*,1})P(\{...,n'_{a'-1,a'^*,1},n'_{a',a'^*,1},...\},t)\big]+ \\
\big[r \sum_{\{n_{a',a'^*,\beta}\}} \sum_{a'=0}^{\infty} \sum_{a'^*=0}^{a'} \big[(a')(n_{a',a'^*,1})(n'_{a'-1,a'^*,1})P(\{...,n'_{a'-1,a'^*,1},n'_{a',a'^*,1},...\},t)\big]+ \\
\big[r \sum_{a\neq(a'-1),a'}^{\infty} \sum_{a^*=0}^{a} \sum_{\{n_{a,a^*,\beta}\}} \sum_{a'=0}^{\infty} \sum_{a'^*=0}^{a'} \big[(an_{a,a^*,1})(n'_{a'-1,a'^*,1})P(\{...,n'_{a'-1,a'^*,1},n'_{a',a'^*,1},...\},t)\big] \\
= r \sum_{\{n_{a',a'^*,\beta}\}} \sum_{a'=0}^{\infty} \sum_{a'^*=0}^{a'} \big[(a'-1)(n'_{a'-1,a'^*,1}-1)(n'_{a'-1,a'^*,1})P(\{...,n'_{a'-1,a'^*,1},n'_{a',a'^*,1},...\},t)+ \\
(a')(n'_{a',a'^*,1}+1)(n'_{a'-1,a'^*,1})P(\{...,n'_{a'-1,a'^*,1},n'_{a',a'^*,1},...\},t)+ \\
\sum_{a\neq(a'-1),a'}^{\infty} \sum_{a^*=0}^{a} (an_{a,a^*,1})(n'_{a'-1,a'^*,1})P(\{...,n'_{a'-1,a'^*,1},n'_{a',a'^*,1},...\},t)\big] \\
= r \sum_{\{n_{a',a'^*,\beta}\}} \sum_{a'=0}^{\infty} \sum_{a'^*=0}^{a'} \big[(a'-1)(n'_{a'-1,a'^*,1})(n'_{a'-1,a'^*,1})P(\{...,n'_{a'-1,a'^*,1},n'_{a',a'^*,1},...\},t)+ \\
(n'_{a'-1,a'^*,1})P(\{...,n'_{a'-1,a'^*,1},n'_{a',a'^*,1},...\},t)+ \\
(a')(n'_{a',a'^*,1})(n'_{a'-1,a'^*,1})P(\{...,n'_{a'-1,a'^*,1},n'_{a',a'^*,1},...\},t)+ \\
\sum_{a\neq(a'-1),a'}^{\infty} \sum_{a^*=0}^{a} (an_{a,a^*,1})(n'_{a'-1,a'^*,1})P(\{...,n'_{a'-1,a'^*,1},n'_{a',a'^*,1},...\},t)\big]
\end{aligned} \tag{31}$$

We have

$$I_2 = r \sum_{a=0}^{\infty} \sum_{a^*=0}^{a} \sum_{\{n_{a,a^*,\beta}\}} \sum_{a'=0}^{\infty} \sum_{a'^*=0}^{a'} \big[(an_{a,a^*,1})(n_{a',a'^*,1})P(\{...,n_{a'-1,a'^*,1},n_{a',a'^*,1},...\},t)\big] \tag{32}$$

Note that $n$ and $n'$ are dummy indices. In the first term, $(a' - 1)$ can be considered as $a'$ as limit of $a'$ starts from 0. Subtracting $I_2$ in Eq. (32) from $I_1$ in Eq. (31) all the square terms $a'n'_{a'}n'_{a'}$, cross-square terms $a'n'_{a'}n'_{(a'-1)}$ and $an_a n'_{(a'-1)}$ are canceled with the terms of $an_a n_{a'}$ in $I_2$ giving,

$$(I_1 - I_2) = r \sum_{\{n_{a',a'^*,\beta}\}} \sum_{a'=0}^{\infty} \sum_{a'^*=0}^{a'} \big[(n'_{a'-1,a'^*,1})P(\{...,n'_{a'-1,a'^*,1},n'_{a',a'^*,1},...\},t) = r \sum_{a'=0}^{\infty} \sum_{a'^*=0}^{a'} \overline{(n'_{a'-1,a'^*,1})} = rn_1 \tag{33}$$

Thus, the contribution of protein production is $rn_1$ to $dA_1/dt$. Similarly, it can be shown that the contribution is $rn_\beta$ to $dA_\beta/dt$.

##### 4.2 Contribution of unphosphorylated-protein degradation $A^U \rightarrow \phi$ in $\frac{dA^U_\beta}{dt}$ at state-$\beta$ :

$$\frac{dA^U_\beta}{dt} = \sum_{a=0}^{\infty} \sum_{a^*=0}^{a} \sum_{\{n_{a,a^*,\beta}\}} (an_{a,a^*,\beta}) \frac{dP(\{...,n_{a,a^*,\beta},...\},t)}{dt} = \sum_{a=0}^{\infty} \sum_{a^*=0}^{a} \sum_{\{n_{a,a^*,\beta}\}} (an_{a,a^*,\beta})[T^{degradation}_\beta]$$

$$\begin{aligned}
= \sum_{a=0}^{\infty} \sum_{a^*=0}^{a} \sum_{\{n_{a,a^*,\beta}\}} (an_{a,a^*,\beta})\delta \sum_{a'=0}^{\infty} \sum_{a'^*=0}^{a'} \big[(a'+1-a'^*)(n_{a'+1,a'^*,\beta}+1)P(\{...,n_{a'+1,a'^*,\beta}+1,n_{a',a'^*,\beta}-1,...\},t) \\
-(a'-a'^*)(n_{a',a'^*,\beta})P(\{...,n_{a'+1,a'^*,\beta},n_{a',a'^*,\beta},...\},t)\big] = I_1 - I_2
\end{aligned} \tag{34}$$

Defining
$n_{a'+1,a'^*,\beta}+1 = n'_{a'+1,a'^*,\beta}$,
$n_{a',a'^*,\beta}-1 = n'_{a',a'^*,\beta}$
we get

$$I_1 - I_2 = \sum_{a=0}^{\infty}\sum_{a^*=0}^{a}\sum_{\{n_{a,a^*,\beta}\}}(an_{a,a^*,\beta})\delta\sum_{a'=0}^{\infty}\sum_{a'^*=0}^{a'}\Big[(a'+1-a'^*)(n'_{a'+1,a'^*,\beta})P(\{...,n'_{a'+1,a'^*,\beta},n'_{a',a'^*,\beta},...\},t) \\ -(a'-a'^*)(n_{a',a'^*,\beta})P(\{...,n_{a'+1,a'^*,\beta},n_{a',a'^*,\beta},...\},t)\Big] \tag{35}$$

Expanding $I_1$ for $a = a'+1$, $a^* = a'^*$, and $a = a'$, $a^* = a'^*$ and $a \neq a'+1$ or $a'$, $a^* \neq a'^*$, we get

$$I_1 = \Big[\sum_{\{n_{a',a'^*,\beta}\}}\sum_{a'=0}^{\infty}\sum_{a'^*=0}^{a'}(a'+1)(n_{a'+1,a'^*,\beta})\delta\big[(a'+1-a'^*)(n'_{a'+1,a'^*,\beta})P(\{...,n'_{a'+1,a'^*,\beta},n'_{a',a'^*,\beta},...\},t)\big] \\ +\Big[\sum_{\{n_{a',a'^*,\beta}\}}\sum_{a'=0}^{\infty}\sum_{a'^*=0}^{a'}(a')(n_{a',a'^*,\beta})\delta\big[(a'+1-a'^*)(n'_{a'+1,a'^*,\beta})P(\{...,n'_{a'+1,a'^*+1,\beta},n'_{a',a'^*,\beta},...\},t)\big]+ \\ \Big[\sum_{a\neq(a'-1),a'}^{\infty}\sum_{a^*\neq a'^*}^{a}\sum_{\{n_{a,a^*,\beta}\}}\sum_{a'=0}^{\infty}\sum_{a'^*=0}^{a'}(an_{a,a^*,\beta})\delta\big[(a'+1-a'^*)(n'_{a'+1,a'^*,\beta})P(\{...,n'_{a'+1,a'^*,\beta},n'_{a',a'^*,\beta},...\},t)\big] \\ = \delta\sum_{\{n_{a',a'^*,\beta}\}}\sum_{a'=0}^{\infty}\sum_{a'^*=0}^{a'}\Bigg[[(a'+1)(n'_{a'+1,a'^*,\beta}-1)\big[(a'+1-a'^*)(n'_{a'+1,a'^*,\beta})P(\{...,n'_{a'+1,a'^*,\beta},n'_{a',a'^*,\beta},...\},t)\big] \\ +\big[(a')(n'_{a',a'^*,\beta}+1)\big[(a'+1-a'^*)(n'_{a'+1,a'^*,\beta})P(\{...,n'_{a'+1,a'^*+1,\beta},n'_{a',a'^*,\beta},...\},t)\big]+ \\ \Big[\sum_{a\neq(a'-1),a'}^{\infty}\sum_{a^*\neq a'^*}^{a}(an_{a,a^*,\beta})\big[(a'+1-a'^*)(n'_{a'+1,a'^*,\beta})P(\{...,n'_{a'+1,a'^*,\beta},n'_{a',a'^*,\beta},...\},t)\big]\Bigg] \\ = \delta\sum_{\{n_{a',a'^*,\beta}\}}\sum_{a'=0}^{\infty}\sum_{a'^*=0}^{a'}\Bigg[[(a'+1)(n'_{a'+1,a'^*,\beta})\big[(a'+1-a'^*)(n'_{a'+1,a'^*,\beta})P(\{...,n'_{a'+1,a'^*,\beta},n'_{a',a'^*,\beta},...\},t)\big] \\ -\big[(a'+1-a'^*)(n'_{a'+1,a'^*,\beta})P(\{...,n'_{a'+1,a'^*,\beta},n'_{a',a'^*,\beta},...\},t)\big] \\ +\big[(a')(n'_{a',a'^*,\beta})\big[(a'+1-a'^*)(n'_{a'+1,a'^*,\beta})P(\{...,n'_{a'+1,a'^*+1,\beta},n'_{a',a'^*,\beta},...\},t)\big]+ \\ \Big[\sum_{a\neq(a'-1),a'}^{\infty}\sum_{a^*\neq a'^*}^{a}(an_{a,a^*,\beta})\big[(a'+1-a'^*)(n'_{a'+1,a'^*,\beta})P(\{...,n'_{a'+1,a'^*,\beta},n'_{a',a'^*,\beta},...\},t)\big]\Bigg] \tag{36}$$

we have

$$I_2 = \sum_{a=0}^{\infty}\sum_{a^*=0}^{a}\sum_{\{n_{a,a^*,\beta}\}}(an_{a,a^*,\beta})\delta\sum_{a'=0}^{\infty}\sum_{a'^*=0}^{a'}\big[(a'-a'^*)(n_{a',a'^*,\beta})P(\{...,n_{a'+1,a'^*,\beta},n_{a',a'^*,\beta},...\},t)\big] \tag{37}$$

Subtracting $I_2$ from $I_1$ canceled all the terms except

$$I_1 - I_2 = -\sum_{\{n_{a,a^*,\beta}\}}\delta\sum_{a'=0}^{\infty}\sum_{a'^*=0}^{a'}\big[(a'+1-a'^*)(n'_{a'+1,a'^*,\beta})P(\{...,n'_{a'+1,a'^*,\beta},n'_{a',a'^*,\beta},...\},t)\big] = -\delta(A_\beta - A^*_\beta) = -\delta A^U_\beta \tag{38}$$

To obtain the time-dependent variation in the total protein $A_\beta$, we added up the effect from $A^U_\beta$ and $A^*_\beta$ as $dA_\beta/dt = (dA^U_\beta/dt + dA^*_\beta/dt)$.

#### 4.3 Contribution to the phosphorylated protein number $A^*$ :

##### 4.3.1 Contribution of cell cycle transition from $\beta \to \beta+1$ :

Following a similar derivation pertaining to Eq. (25) it can be shown the contribution to $dA^*_\beta/dt$ and $dA^*_{\beta+1}/dt$ are $-k_\beta A^*_\beta$ and $k_\beta A^*_\beta$, respectively.

###### 4.3.2 Contribution of cell cycle transition from state $N \to 1$ :

**No heritability condition:**
The contribution to $dA_1^*/dt$ vanishes as all the $A^*$ molecules are dephosphorylated upon $N \to 1$ transition.
**Full heritability condition:** The contribution of $T_N^{cell\ cycle}$ to the kinetics of $A_1^*$ is,

$$\begin{aligned}\frac{dA_1^*}{dt} &= \sum_{a=0}^{\infty}\sum_{a^*=0}^{a}\sum_{\{n_{a,a^*,\beta}\}}(a^* n_{a,a^*,1})\frac{dP(\{...,n_{a,a^*,1},...\},t)}{dt} = \sum_{a=0}^{\infty}\sum_{a^*=0}^{a}\sum_{\{n_{a,a^*,\beta}\}}(a^* n_{a,a^*,1})T_N^{cell\ cycle}\\ &= \sum_{a=0}^{\infty}\sum_{a^*=0}^{a}\sum_{\{n_{a,a^*,\beta}\}}(a^* n_{a,a^*,1})\sum_{a'=0}^{\infty}\sum_{a'^*=0}^{a'}\sum_{m=0}^{a'}\sum_{m^*=0}^{\tilde{a}'^*}k_N({}^{a'}C_m 2^{-a'})({}^{a'^*}C_{m^*}2^{-a'^*}) \\ &\quad \big[(n_{a',a'^*,N}+1)P(\{...,n_{a',a'^*,N}+1,n_{m,m^*,1}-1,n_{a'-m,a'^*-m^*,1}-1,...\},t)\\ &\quad -(n_{a',a'^*,N})P(\{...,n_{a',a'^*,N},n_{m,m^*,1},n_{a'-m,a'^*-m^*,1}...\},t)\big]\end{aligned} \quad (39)$$

Defining
$n_{a',a'^*,N}+1 = n'_{a',a'^*,N}$,
$n_{m,m^*,1}-1 = n'_{m,m^*,1}$
$n_{a'-m,a'^*-m^*,1}-1 = n'_{a'-m,a'^*-m^*,1}$
we get

$$\begin{aligned}(I_1 - I_2) &= \sum_{a=0}^{\infty}\sum_{a^*=0}^{a}\sum_{\{n_{a,a^*,\beta}\}}(a^* n_{a,a^*,1})\sum_{a'=0}^{\infty}\sum_{a'^*=0}^{a'}\sum_{m=0}^{a'}\sum_{m^*=0}^{a'^*}k_N({}^{a'}C_m 2^{-a'})({}^{a'^*}C_{m^*}2^{-a'^*})\\ &\quad \big[(n'_{a',a'^*,N})P(\{...,n'_{a',a'^*,N},n'_{m,m^*,1},n'_{a'-m,a'^*-m^*,1},...\},t)\\ &\quad -(n_{a',a'^*,N})P(\{...,n_{a',a'^*,N},n_{m,m^*,1},n_{a'-m,a'^*-m^*,1}...\},t)\big]\end{aligned} \quad (40)$$

For the terms $a = m,\ a^* = m^*$ and $a = a' - m,\ a^* = a'^* - m^*$ we get,

$$\begin{aligned}(I_1 - I_2) &= \sum_{\{n_{a',a'^*,\beta}\}}\sum_{a'=0}^{\infty}\sum_{a'^*=0}^{a'}\sum_{m=0}^{a'}\sum_{m^*=0}^{\tilde{a}'^*}(m^* n_{m,m^*,1}+(a'^*-m^*)n_{a'-m,a'^*-m^*,1})k_N({}^{a'}C_m 2^{-a'})({}^{a'^*}C_{m^*}2^{-a'^*})\\ &\quad \big[(n'_{a',a'^*,N})P(\{...,n'_{a',a'^*,N},n'_{m,m^*,1},n'_{a'-m,a'^*-m^*,1},...\},t)\\ &\quad -(n_{a',a'^*,N})P(\{...,n_{a',a'^*,N},n_{m,m^*,1},n_{a'-m,a'^*-m^*,1}...\},t)\big]\\ &= \sum_{\{n_{a',a'^*,\beta}\}}\sum_{a'=0}^{\infty}\sum_{a'^*=0}^{a'}\sum_{m=0}^{a'}\sum_{m^*=0}^{\tilde{a}'^*}k_N({}^{a'}C_m 2^{-a'})({}^{a'^*}C_{m^*}2^{-a'^*})\\ &\quad \big[(m^*(n'_{m,m^*,1}+1)+(a'^*-m^*)(n'_{a'-m,a'^*-m^*,1}+1)(n'_{a',a'^*,N})P(\{...,n'_{a',a'^*,N},n'_{m,m^*,1},n'_{a'-m,a'^*-m^*,1},...\},t)\\ &\quad -(m^* n_{m,m^*,1}+(a'^*-m^*)n_{a'-m,a'^*-m^*,1})(n_{a',a'^*,N})P(\{...,n_{a',a'^*,N},n_{m,m^*,1},n_{a'-m,a'^*-m^*,1}...\},t)\big]\\ &= \sum_{\{n_{a',a'^*,\beta}\}}\sum_{a'=0}^{\infty}\sum_{a'^*=0}^{a'}\sum_{m=0}^{a'}\sum_{m^*=0}^{\tilde{a}'^*}k_N({}^{a'}C_m 2^{-a'})({}^{a'^*}C_{m^*}2^{-a'^*})\\ &\quad \big[(a'^*)(n'_{a',a'^*,N})P(\{...,n'_{a',a'^*,N},n'_{m,m^*,1},n'_{a'-m,a'^*-m^*,1},...\},t)\big]\\ &= \sum_{a'=0}^{\infty}\sum_{a'^*=0}^{a'}\big[\sum_{m=0}^{a'}\sum_{m^*=0}^{\tilde{a}'^*}k_N({}^{a'}C_m 2^{-a'})({}^{a'^*}C_{m^*}2^{-a'^*})(a'^*)\overline{(n'_{a',a'^*,N})}\big] = k_N A_N^*\end{aligned} \quad (41)$$

Thus, the contribution to $dA_1^*/dt$ is $k_N A_N^*$.

###### 4.3.3 Contribution of phosphorylation $A \to A^*$ at state $\beta$:

Total number of phosphorylated proteins $A^*$ in state-$\beta$ is,

$$\frac{dA^*_\beta}{dt} = \sum_{a=0}^{\infty}\sum_{a^*=0}^{a}\sum_{\{n_{a,a^*,\beta}\}}(a^* n_{a,a^*,\beta})\frac{dP(\{...,n_{a,a^*,\beta},...\},t)}{dt} = \sum_{a=0}^{\infty}\sum_{a^*=0}^{a}\sum_{\{n_{a,a^*,\beta}\}}(a^* n_{a,a^*,\beta})[T^{phospho}_\beta]$$

$$= \sum_{a=0}^{\infty}\sum_{a^*=0}^{a}\sum_{\{n_{a,a^*,\beta}\}}(a^* n_{a,a^*,\beta})\big[\sum_{a'=0}^{\infty}\sum_{a'^*=0}^{a'} r_p(a' - (a'^* - 1))(n_{a',a'^*-1,\beta}+1)P(\{...,n_{a',a'^*-1,\beta}+1, n_{a',a'^*,\beta}-1,...\},t)$$
$$-\sum_{a'=0}^{\infty}\sum_{a'^*=0}^{a'} r_p(a' - a'^*)(n_{a',a'^*,\beta})P(\{...,n_{a',a'^*-1,\beta}, n_{a',a'^*,\beta},...\},t)\big] \tag{42}$$

Defining
$n_{a',a'^*-1,\beta}+1 = n'_{a',a'^*-1,\beta}$,
$n_{a',a'^*,\beta}-1 = n'_{a',a'^*,\beta}$

we get

$$I_1 - I_2 = \sum_{a=0}^{\infty}\sum_{a^*=0}^{a}\sum_{\{n_{a',a'^*,\beta}\}}(a^* n_{a,a^*,\beta})\sum_{a'=0}^{\infty}\sum_{a'^*=0}^{a'} r_p\big[(a' - (a'^* - 1))(n'_{a',a'^*-1,\beta})P(\{...,n'_{a',a'^*-1,\beta}, n'_{a',a'^*,\beta},...\},t)$$
$$-(a' - a'^*)(n_{a',a'^*,\beta})P(\{...,n_{a',a'^*-1,\beta}, n_{a',a'^*,\beta},...\},t)\big] \tag{43}$$

For the terms $a = a',\ a^* = a'^* - 1$, we get the terms of $I_1$,

$$I_1^{(1)} = \sum_{a'=0}^{\infty}\sum_{a'^*=0}^{a'}\sum_{\{n_{a',a'^*,\beta}\}}((a'^* - 1)n_{a',a'^*-1,\beta})r_p(a' - (a'^* - 1))(n'_{a',a'^*-1,\beta})P(\{...,n'_{a',a'^*-1,\beta}, n'_{a',a'^*,\beta},...\},t)$$
$$= \sum_{a'=0}^{\infty}\sum_{a'^*=0}^{a'}\sum_{\{n_{a',a'^*,\beta}\}}((a'^* - 1)(n'_{a',a'^*-1,\beta} - 1)r_p(a' - (a'^* - 1))(n'_{a',a'^*-1,\beta})P(\{...,n'_{a',a'^*-1,\beta}, n'_{a',a'^*,\beta},...\},t) \tag{44}$$

For the terms $a = a',\ a^* = a'^*$ we get the terms of $I_1$,

$$I_1^{(2)} = \sum_{a'=0}^{\infty}\sum_{a'^*=0}^{a'}\sum_{\{n_{a',a'^*,\beta}\}}(a'^* n_{a',a'^*,\beta})r_p(a' - (a'^* - 1))(n'_{a',a'^*-1,\beta})P(\{...,n'_{a',a'^*-1,\beta}, n'_{a',a'^*,\beta},...\},t)$$
$$= \sum_{a'=0}^{\infty}\sum_{a'^*=0}^{a'}\sum_{\{n_{a',a'^*,\beta}\}} a'^*(n'_{a',a'^*-1,\beta}+1)r_p(a' - (a'^* - 1))(n'_{a',a'^*,\beta})P(\{...,n'_{a',a'^*-1,\beta}, n'_{a',a'^*,\beta},...\},t) \tag{45}$$

For the terms $a \neq a',\ a^* \neq (a'^* - 1)$ and $a \neq a',\ a^* \neq a'^*$ we get the terms of $I_1$,

$$I_1^{(3)} = \sum_{a'=0}^{\infty}\sum_{a'^*=0}^{a'}\sum_{\{n_{a',a'^*,\beta}\}}(a^* n_{a,a^*,\beta})r_p(a' - (a'^* - 1))(n'_{a',a'^*-1,\beta})P(\{...,n'_{a',a'^*-1,\beta}, n'_{a',a'^*,\beta},...\},t) \tag{46}$$

$$I_2 = \sum_{a=0}^{\infty}\sum_{a^*=0}^{a}\sum_{a'=0}^{\infty}\sum_{a'^*=0}^{a'}\sum_{\{n_{a',a'^*,\beta}\}}(a^* n_{a,a^*,\beta})r_p(a' - a'^*)(n_{a',a'^*,\beta})P(\{...,n_{a',a'^*-1,\beta}, n_{a',a'^*,\beta},...\},t) \tag{47}$$

$$
\begin{aligned}
I_1 - I_2 &= I_1^{(1)} + I_1^{(2)} + I_1^{(3)} - I_2 \\
= \sum_{a=0}^{\infty}\sum_{a^*=0}^{a}\sum_{\{n_{a',a'^*,\beta}\}}\sum_{a'=0}^{\infty}\sum_{a'^*=0}^{a'} &-(a'^* - 1)r_p(a' - (a'^* - 1))(n'_{a',a'^*-1,\beta})P(\{..., n'_{a',a'^*-1,\beta}, n'_{a',a'^*,\beta}, ...\}, t) \\
&+ a'^* r_p(a' - (a'^* - 1))(n'_{a',a'^*,\beta})P(\{..., n'_{a',a'^*-1,\beta}, n'_{a',a'^*,\beta}, ...\}, t) \\
&= \sum_{a=0}^{\infty}\sum_{a^*=0}^{a}\sum_{\{n_{a',a'^*,\beta}\}}\sum_{a'=0}^{\infty}\sum_{a'^*=0}^{a'} r_p(a' - (a'^* - 1))(n'_{a',a'^*,\beta}) \\
&P(\{..., n'_{a',a'^*-1,\beta}, n'_{a',a'^*,\beta}, ...\}, t) = r_p(A_\beta - A^*_\beta)
\end{aligned}
\tag{48}
$$

### 5 Steady State Solutions :

The relative number of cells in a cell cycle state $\beta$ : $\rho_\beta = \eta_\beta/(\sum_\beta \eta_\beta)$
and the average number of $A$ in single cells residing in cell cycle state $\beta$, $a_\beta = A_\beta/n_\beta$ can be estimated.

#### 5.1 Steady State solution for the relative cell number $\rho_\beta$ at state $\beta$ : $n_1 \to n_2 \to n_N$

Calculation for $\rho_1$ :

$$
\begin{aligned}
\frac{d\rho_1}{dt} &= \frac{d(n_1/n)}{dt} = \frac{1}{n}\frac{d(n_1)}{dt} - \frac{n_1}{n^2}\frac{d(n)}{dt} \\
&= -k_1\rho_1 + 2k_N\rho_N - \frac{\rho_1}{n}\frac{d(n_1 + n_2 + n_N)}{dt} \\
&= -k_1\rho_1 + 2k_N\rho_N - \frac{\rho_1}{n}(-k_1n_1 + 2k_Nn_N + k_1n_1 - k_2n_2 + k_2n_2 - k_Nn_N)) \\
&= -k_1\rho_1 + 2k_N\rho_N - \frac{\rho_1}{n}(2k_Nn_N - k_Nn_N) \\
&= -k_1\rho_1 + 2k_N\rho_N - k_N\rho_1\rho_N \\
&= -k_1\rho_1 + (2 - \rho_1)k_N\rho_N
\end{aligned}
\tag{49}
$$

Steady State value of $\rho_1^{(S)}$ :

$$
\begin{aligned}
\frac{d\rho_1^{(s)}}{dt} = 0 = -k_1\rho_1^{(s)} + (2 - \rho_1^{(s)})k_N\rho_N \\
\boxed{\rho_1^{(s)} = \frac{2k_N\rho_N}{(k_1 + k_N\rho_N)}}
\end{aligned}
\tag{50}
$$

Calculation for $\rho_\beta$ :

$$
\begin{aligned}
\frac{d\rho_2}{dt} &= \frac{d(n_2/n)}{dt} = \frac{1}{n}\frac{d(n_2)}{dt} - \frac{n_2}{n^2}\frac{d(n)}{dt} \\
&= k_1\rho_1 - k_2\rho_2 - \frac{\rho_2}{n}\frac{d(n_1 + n_2 + n_N)}{dt} \\
&= k_1\rho_1 - k_2\rho_2 - \frac{\rho_2}{n}(-k_1n_1 + 2k_Nn_N + k_1n_1 - k_2n_2 + k_2n_2 - k_Nn_N)) \\
&= k_1\rho_1 - k_2\rho_2 - \frac{\rho_2}{n}(2k_Nn_N - k_Nn_N) \\
&= k_1\rho_1 - k_2\rho_2 - k_N\rho_2\rho_N \\
\frac{d\rho_\beta}{dt} &= k_{\beta-1}\rho_{\beta-1} - \rho_\beta(k_\beta + k_N\rho_N)
\end{aligned}
\tag{51}
$$

Steady State value of $\rho_\beta^{(S)}$ :

$$\frac{d\rho_\beta^{(s)}}{dt} = 0 = k_{\beta-1}\rho_{\beta-1}^{(s)} - \rho_\beta^{(s)}(k_\beta + k_N\rho_N^{(s)})$$

$$\text{Assuming } \beta{=}3,\ \rho_3^{(s)} = \frac{k_2\rho_2^{(s)}}{(k_3 + k_N\rho_N^{(s)})} = \frac{k_2 k_1\rho_1^{(s)}}{(k_3 + k_N\rho_N^{(s)})(k_2 + k_N\rho_N^{(s)})}$$

$$\begin{aligned}\rho_\beta^{(s)} &= \frac{k_{\beta-1}k_{\beta-2}...k_1\rho_1^{(s)}}{(k_\beta + k_N\rho_N^{(s)})...(k_2 + k_N\rho_N^{(s)})} \\ &= \frac{k_{\beta-1}k_{\beta-2}...k_1(2k_N\rho_N)}{(k_\beta + k_N\rho_N^{(s)})...(k_2 + k_N\rho_N^{(s)})(k_1 + k_N\rho_N^{(s)})} \\ &\boxed{\rho_\beta^{(s)} = 2\Big[\prod_{i=1}^{\beta-1}\frac{k_i}{(k_i + k_N\rho_N^{(s)})}\Big]\frac{(k_N\rho_N)}{(k_\beta + k_N\rho_N^{(s)})}}\end{aligned} \tag{52}$$

Where $\rho_N$ satisfies

$$\Big[\prod_{i=1}^{N-1}\frac{k_i}{(k_i + k_N\rho_N^{(s)})}\Big]\frac{(k_N)}{(k_N + k_N\rho_N^{(s)})} = \frac{1}{2} \tag{53}$$

#### 5.2 Steady State solution for the average number of proteins $a_\beta = A_\beta/n_\beta$ at state $\beta$ :

Calculation for $a_1$ :

$$\begin{aligned}\frac{da_1}{dt} &= \frac{d(A_1/n_1)}{dt} = \frac{1}{n_1}\frac{d(A_1)}{dt} - \frac{A_1}{n_1^2}\frac{d(n_1)}{dt} \\ &= \frac{1}{n_1}(r^{(1)}n_1 - \delta^{(1)}A_1 - k_1A_1 + k_NA_N) - \frac{A_1}{n_1^2}(-k_1n_1 + 2k_Nn_N) \\ &= (r^{(1)} - \delta^{(1)}a_1 - k_1a_1 + k_Na_N\frac{n_N}{n_1}) - a_1(-k_1 + 2k_N\frac{n_N}{n_1}) \\ &= r^{(1)} - \delta^{(1)}a_1 + (a_N - 2a_1)k_N\frac{\rho_N}{\rho_1}\end{aligned} \tag{54}$$

Steady-state value for $a_1^{(s)}$ :

$$\begin{aligned}\frac{da_1^{(s)}}{dt} &= 0 = r^{(1)} - \delta^{(1)}a_1 + (a_N - 2a_1)k_N\frac{\rho_N}{\rho_1} \\ a_1^{(s)} &= \frac{r^{(1)} + a_Nk_N(\frac{\rho_N}{\rho_1})}{\delta^{(1)} + 2k_N(\frac{\rho_N}{\rho_1})} \\ &\boxed{a_1^{(s)} = \tilde{r}^{(1)} + a_N\tilde{k}_N}\end{aligned} \tag{55}$$

Where, $\tilde{r}^{(1)} = \frac{r^{(1)}}{\delta^{(1)}+2k_N(\frac{\rho_N}{\rho_1})}$ and $\tilde{k}_N = \frac{k_N(\frac{\rho_N}{\rho_1})}{\delta^{(1)}+2k_N\frac{\rho_N}{\rho_1}}$.

Calculation for $a_\beta$ :

$$\begin{aligned}\frac{da_\beta}{dt} &= \frac{d(A_\beta/n_\beta)}{dt} = \frac{1}{n_\beta}\frac{d(A_\beta)}{dt} - \frac{A_\beta}{n_\beta^2}\frac{d(n_\beta)}{dt} \\ &= \frac{1}{n_\beta}(r^{(\beta)}n_\beta - \delta^{(\beta)}A_\beta + k_{\beta-1}A_{\beta-1} - k_\beta A_\beta) - \frac{A_\beta}{n_\beta^2}(k_{\beta-1}n_{\beta-1} - k_\beta n_\beta) \\ &= r^{(\beta)} - \delta^{(\beta)}a_\beta + k_{\beta-1}a_{\beta-1}\frac{\rho_{\beta-1}}{\rho_\beta} - k_\beta a_\beta - a_\beta(k_{\beta-1}\frac{\rho_{\beta-1}}{\rho_\beta} - k_\beta) \\ &= r^{(\beta)} - \delta^{(\beta)}a_\beta + k_{\beta-1}\frac{\rho_{\beta-1}}{\rho_\beta}(a_{\beta-1} - a_\beta)\end{aligned} \tag{56}$$

Steady-state value for $a_\beta^{(s)}$ :

$$\begin{aligned}\frac{da_\beta^{(s)}}{dt} =& 0 = r^{(\beta)} - \delta^{(\beta)}a_\beta + k_{\beta-1}\frac{\rho_{\beta-1}}{\rho_\beta}(a_{\beta-1}^{(s)} - a_\beta^{(s)})\\ a_\beta^{(s)} =& \frac{r^{(\beta)} + k_{\beta-1}(\frac{\rho_{\beta-1}}{\rho_\beta})a_{\beta-1}^{(s)}}{\delta^{(\beta)} + k_{\beta-1}(\frac{\rho_{\beta-1}}{\rho_\beta})}\\ =& \tilde{r}^{(\beta)} + \tilde{k}_{\beta-1}a_{\beta-1}^{(s)}\\ =& \tilde{r}^{(\beta)} + \tilde{k}_{\beta-1}[\tilde{r}^{(\beta-1)} + \tilde{k}_{\beta-2}a_{\beta-2}^{(s)}]\\ =& \tilde{r}^{(\beta)} + \tilde{k}_{\beta-1}\tilde{r}^{(\beta-1)} + \tilde{k}_{\beta-1}\tilde{k}_{\beta-2}a_{\beta-2}^{(s)}\\ =& \tilde{r}^{(\beta)} + \tilde{k}_{\beta-1}\tilde{r}^{(\beta-1)} + \tilde{k}_{\beta-1}\tilde{k}_{\beta-2}\tilde{r}^{(\beta-2)} + \tilde{k}_{\beta-1}\tilde{k}_{\beta-2}\tilde{k}_{\beta-3}a_{\beta-3}^{(s)}\end{aligned} \tag{57}$$

Assuming $\beta = 4$ we get, $a_4^{(s)} = \tilde{r}^{(4)} + \tilde{k}_3\tilde{r}^{(3)} + \tilde{k}_3\tilde{k}_2\tilde{r}^{(2)} + \tilde{k}_3\tilde{k}_2\tilde{k}_1\tilde{a}_1^{(s)}$

$$= \tilde{r}^{(4)} + \tilde{k}_3\tilde{r}^{(3)} + \tilde{k}_3\tilde{k}_2\tilde{r}^{(2)} + \tilde{k}_3\tilde{k}_2\tilde{k}_1(\tilde{r}^{(1)} + a_N\tilde{k}_N)$$

$$\boxed{a_\beta^{(s)} = \tilde{r}^{(\beta)} + \sum_{i=1}^{\beta-1}(\prod_{j=1}^{i}\tilde{k}_{\beta-j})\tilde{r}^{(\beta-i)} + (\prod_{j=1}^{\beta-1}\tilde{k}_{\beta-j})\tilde{k}_N a_N}$$

where $\tilde{r}^{(\beta)} = \frac{r^{(\beta)}}{\delta^{(\beta)}+k_{\beta-1}(\frac{\rho_{\beta-1}}{\rho_\beta})}$ for $2 \le \beta \le N$, $\tilde{k}_\beta = \frac{k_\beta(\frac{\rho_\beta}{\rho_{\beta+1}})}{\delta^{\beta+1}+k_\beta(\frac{\rho_\beta}{\rho_{\beta+1}})}$ for $1 \le \beta \le N-1$, $\tilde{k}_N = \frac{k_N(\frac{\rho_N}{\rho_1})}{\delta^{(1)}+2k_N\frac{\rho_N}{\rho_1}}$ and for $\beta = N$, we get $a_N = \frac{\tilde{r}^{(N)}+\sum_{i=1}^{N-1}\left(\prod_{j=1}^{i}\tilde{k}_{N-j}\right)\tilde{r}^{(N-i)}}{1-\left(\prod_{j=1}^{N-1}\tilde{k}_{N-j}\right)\tilde{k}_N}$.

##### 5.3 Steady State solution for the average number of phosphorylated proteins $a_\beta^* = A_\beta^*/n_\beta$ at state $\beta$ :

Calculation for $a_1^*$ (No heritability case) :

$$\begin{aligned}\frac{da_1^*}{dt} &= \frac{d(A_1^*/n_1)}{dt} = \frac{1}{n_1}\frac{d(A_1^*)}{dt} - \frac{A_1^*}{n_1^2}\frac{d(n_1)}{dt}\\ &= \frac{1}{n_1}(r_p^{(1)}A_1 - (r_p^{(1)} + r_d^{(1)} + \delta^{(1)} + k_1)A_1^*) - a_1^*(-k_1 + 2k_N\frac{n_N}{n_1})\\ &= r_p^{(1)}a_1 - (r_p^{(1)} + r_d^{(1)} + \delta^{(1)} + k_1)a_1^* - a_1^*(-k_1 + 2k_N\frac{\rho_N}{\rho_1})\\ &= r_p^{(1)}a_1 - (\lambda^{(1)} + 2k_N\frac{\rho_N}{\rho_1})a_1^*\end{aligned} \tag{58}$$

Calculation for $a_1^*$ (Full heritability case) :

$$\begin{aligned}\frac{da_1^*}{dt} &= \frac{d(A_1^*/n_1)}{dt} = \frac{1}{n_1}\frac{d(A_1^*)}{dt} - \frac{A_1^*}{n_1^2}\frac{d(n_1)}{dt}\\ &= \frac{1}{n_1}(r_p^{(1)}A_1 - (r_p^{(1)} + r_d^{(1)} + \delta^{(1)} + k_1)A_1^* + k_N A_N^*)) - a_1^*(-k_1 + 2k_N\frac{n_N}{n_1})\\ &= r_p^{(1)}a_1 - (r_p^{(1)} + r_d^{(1)} + \delta^{(1)} + k_1)a_1^* + k_N a_N^*(\frac{\rho_N}{\rho_1}) - a_1^*(-k_1 + 2k_N\frac{\rho_N}{\rho_1})\\ &= r_p^{(1)}a_1 - (\lambda^{(1)} + 2k_N\frac{\rho_N}{\rho_1})a_1^* + k_N(\frac{\rho_N}{\rho_1})a_N^*\end{aligned} \tag{59}$$

Steady State values for $a_1^{*(s)}$ (Full heritability case) :

$$\begin{aligned}\frac{da_1^{*(s)}}{dt} &= 0 = r_p^{(1)}a_1 - (\lambda^{(1)} + 2k_N\frac{\rho_N}{\rho_1})a_1^* + k_N(\frac{\rho_N}{\rho_1})a_N^*\\ &\boxed{a_1^{*(s)} = \frac{r_p^{(1)}a_1 + k_N(\frac{\rho_N}{\rho_1})a_N^*}{\lambda^{(1)} + 2k_N(\frac{\rho_N}{\rho_1})} = \tilde{r}_p^{(1)}a_1 + \tilde{k}_N^{(p)}a_N^*}\end{aligned} \tag{60}$$

Where $\tilde{r}_p^{(1)} = \frac{r_p^{(1)}}{\lambda^{(1)}+2k_N(\frac{\rho_N}{\rho_1})}$ and $\tilde{k}_N^{(p)} = \frac{k_N(\frac{\rho_N}{\rho_1})}{\lambda^{(1)}+2k_N(\frac{\rho_N}{\rho_1})} = \tilde{r}_p^{(1)} a_1 + \tilde{k}_N^{(p)} a_N^*$

Calculation for $a_\beta^*$ (No or full heritability case depends on the value of $a_1^{*(s)}$) :

$$\begin{aligned}
\frac{da_\beta^*}{dt} &= \frac{d(A_\beta^*/n_\beta)}{dt} = \frac{1}{n_\beta}\frac{d(A_\beta^*)}{dt} - \frac{A_\beta^*}{n_\beta^2}\frac{d(n_\beta)}{dt} \\
&= \frac{1}{n_\beta}(r_p^{(\beta)} A_\beta - (r_p^{(\beta)} + r_d^{(\beta)} + \delta^{(\beta)} + k_\beta) A_\beta^* + k_{\beta-1} A_{\beta-1}^*) - \frac{a_\beta^*}{n^\beta}(k_{\beta-1} n_{\beta-1} - k_\beta n_\beta) \\
&= r_p^{(\beta)} a_\beta - (r_p^{(\beta)} + r_d^{(\beta)} + \delta^{(\beta)} + k_\beta) a_\beta^* + k_{\beta-1} a_{\beta-1}^* (\frac{n_{\beta-1}}{n_\beta}) - a_\beta^* k_{\beta-1}(\frac{n_{\beta-1}}{n_\beta}) + a_\beta^* k_\beta \\
&= r_p^{(\beta)} a_\beta - (\lambda^{(\beta)} + k_{\beta-1}(\frac{\rho_{\beta-1}}{\rho_\beta})) a_\beta^* + k_{\beta-1}(\frac{\rho_{\beta-1}}{\rho_\beta}) a_{\beta-1}^* \text{ for } 2 \le \beta \le N
\end{aligned} \tag{61}$$

Where $\lambda^{(\beta)} = (r_p^{(\beta)} + r_d^{(\beta)} + \delta^{(\beta)})$ for $1 \le \beta \le N$.

Steady State values for $a_\beta^{*(s)}$ (Full heritability case) :

$$\begin{aligned}
\frac{da_\beta^{*(s)}}{dt} &= 0 = r_p^{(\beta)} a_\beta - (\lambda^{(\beta)} + k_{\beta-1}(\frac{\rho_{\beta-1}}{\rho_\beta})) a_\beta^{*(s)} + k_{\beta-1}(\frac{\rho_{\beta-1}}{\rho_\beta}) a_{\beta-1}^{*(s)} \\
a_\beta^{*(s)} &= \frac{r_p^{(\beta)} a_\beta + k_{\beta-1}(\frac{\rho_{\beta-1}}{\rho_\beta}) a_{\beta-1}^{*(s)}}{(\lambda^{(\beta)} + k_{\beta-1}(\frac{\rho_{\beta-1}}{\rho_\beta}))} = \tilde{r}_p^{(\beta)} a_\beta + \tilde{k}_{\beta-1}^{(p)} a_{\beta-1}^{*(s)} \\
&= \tilde{r}_p^{(\beta)} a_\beta + \tilde{k}_{\beta-1}^{(p)} [\tilde{r}_p^{(\beta-1)} a_{\beta-1} + \tilde{k}_{\beta-2}^{(p)} a_{\beta-2}^{*(s)}] \\
&= \tilde{r}_p^{(\beta)} a_\beta + \tilde{k}_{\beta-1}^{(p)} \tilde{r}_p^{(\beta-1)} a_{\beta-1} + \tilde{k}_{\beta-1}^{(p)} \tilde{k}_{\beta-2}^{(p)} a_{\beta-2}^{*(s)} \\
&\text{For } \beta = 3,\ a_3^{*(s)} = \tilde{r}_p^{(3)} a_3 + \tilde{k}_2^{(p)} \tilde{r}_p^{(2)} a_2 + \tilde{k}_2^{(p)} \tilde{k}_1^{(p)} a_1^{*(s)} \\
&= \tilde{r}_p^{(3)} a_3 + \tilde{k}_2^{(p)} \tilde{r}_p^{(2)} a_2 + \tilde{k}_2^{(p)} \tilde{k}_1^{(p)} (\tilde{r}_p^{(1)} a_1 + \tilde{k}_N^{(p)} a_N^*) \\
&\boxed{a_\beta^{*(s)} = \tilde{r}_p^{(\beta)} a_\beta + \sum_{i=1}^{\beta-1} \left( \prod_{j=1}^{i} \tilde{k}_{\beta-j}^{(p)} \right) \tilde{r}_p^{(\beta-i)} a_{(\beta-i)} + \left( \prod_{j=1}^{\beta-1} \tilde{k}_{\beta-j}^{(p)} \right) \tilde{k}_N^{(p)} a_N^*}
\end{aligned} \tag{62}$$

Where, $\tilde{r}_p^{(\beta)} = \frac{r_p^{(\beta)}}{(\lambda^{(\beta)}+k_{\beta-1}(\frac{\rho_{\beta-1}}{\rho_\beta}))}$, $\tilde{k}_\beta^{(p)} = \frac{k_\beta(\frac{\rho_\beta}{\rho_{(\beta+1)}})}{(\lambda^{(\beta+1)}+k_\beta(\frac{\rho_\beta}{\rho_{\beta+1}}))}$ for $2 \le \beta \le (N-1)$

and for $\beta = N$, we get $a_N^* = \frac{\tilde{r}_p^{(N)} a_N + \sum_{i=1}^{N-1} \left( \prod_{j=1}^{i} \tilde{k}_{N-j}^{(p)} \right) \tilde{r}_p^{(N-i)} a_{N-i}}{1 - \left( \prod_{j=1}^{N-j} \tilde{k}_{N-j}^{(p)} \right) \tilde{k}_N^{(p)}}$

### 6 Calculation of $\rho_{(\beta+1)}/\rho_\beta$ :

In steady state, we know

$$\rho_\beta = 2 \left[ \prod_{i=1}^{\beta-1} \frac{k_i}{k_\beta + k_N \rho_N} \right] \frac{k_N \rho_N}{k_\beta + k_N \rho_N} \tag{63}$$

Thus we get the ratio of $\rho_{\beta+1}$ and $\rho_\beta$ as

$$\frac{\rho_{\beta+1}}{\rho_\beta} = \frac{k_\beta}{k_{\beta+1}} \tag{64}$$

Because, the rates $k_\beta$ of cell cycle transitions increase with increasing state $\beta$ or $k_{\beta+1} > k_\beta$. The ratio will always be less than 1 in steady state or the relative number of cells at cell cycle stages decrease with the cell cycle state $\beta$ or $\rho_{\beta+1} < \rho_\beta$ in the steady state limit.

### 7 Calculation of $a_{(\beta+1)}/a_\beta$ :

For $2 \leq \beta \leq (N-1)$ we get.

$$\frac{a^{(s)}_{(\beta+1)}}{a^{(s)}_{\beta}} = \frac{\tilde{r}^{(\beta+1)} + \sum_{i=1}^{\beta}\big(\prod_{j=1}^{i}\tilde{k}_{\beta+1-j}\big)\tilde{r}^{(\beta+1-i)} + \big(\prod_{j=1}^{\beta}\tilde{k}_{\beta+1-j}\big)\tilde{k}_N a_N}{\tilde{r}^{(\beta)} + \sum_{i=1}^{\beta-1}\big(\prod_{j=1}^{i}\tilde{k}_{\beta-j}\big)\tilde{r}^{(\beta-i)} + \big(\prod_{j=1}^{\beta-1}\tilde{k}_{\beta-j}\big)\tilde{k}_N a_N} \tag{65}$$

Considering the simple system where $\beta = 2$ we get,

$$\begin{aligned}\frac{a^{(s)}_{3}}{a^{(s)}_{2}} &= \frac{\tilde{r}^{(3)} + \tilde{k}_2\tilde{r}^{(2)} + \tilde{k}_2\tilde{k}_1\tilde{r}^{(1)} + a_N\tilde{k}_2\tilde{k}_1\tilde{k}_N}{\tilde{r}^{(2)} + \tilde{k}_1\tilde{r}^{(1)} + a_N\tilde{k}_1\tilde{k}_N}\\ &= \frac{\tilde{r}^{(3)} + \tilde{k}_2\Big(\tilde{r}^{(2)} + \tilde{k}_1\tilde{r}^{(1)} + a_N\tilde{k}_1\tilde{k}_N\Big)}{\tilde{r}^{(2)} + \tilde{k}_1\tilde{r}^{(1)} + a_N\tilde{k}_1\tilde{k}_N}\\ &= \frac{\tilde{r}^{(3)}}{\tilde{r}^{(2)} + \tilde{k}_1(\tilde{r}^{(1)} + a_N\tilde{k}_N)} + \tilde{k}_2\\ &= \frac{\tilde{r}^{(3)}}{a^{(s)}_2} + \tilde{k}_2\end{aligned} \tag{66}$$

Similarly for $\beta = 3$ we get,

$$\frac{a^{(s)}_{4}}{a^{(s)}_{3}} = \frac{\tilde{r}^{(4)}}{\tilde{r}^{(3)} + \tilde{k}_2(\tilde{r}^{(2)} + \tilde{k}_1\tilde{r}^{(1)} + \tilde{k}_1\tilde{k}_N a_N)} + \tilde{k}_3 \tag{67}$$

So, we get

$$\boxed{\frac{a^{(s)}_{(\beta+1)}}{a^{(s)}_{(\beta)}} = \frac{\tilde{r}^{(\beta+1)}}{a^{(s)}_{\beta}} + \tilde{k}_\beta} \tag{68}$$

$$\boxed{\frac{a^{(s)}_{(\beta+1)}}{a^{(s)}_{(\beta)}} = \frac{\tilde{r}^{(\beta+1)}}{\tilde{r}^{(\beta)} + \sum_{i=1}^{\beta-1}\big(\prod_{j=1}^{i}\tilde{k}_{\beta-j}\big)\tilde{r}^{(\beta-i)} + \big(\prod_{j=1}^{\beta-1}\tilde{k}_{\beta-j}\big)\tilde{k}_N a_N} + \tilde{k}_\beta} \tag{69}$$

Where, $a^{(s)}_{\beta} = \tilde{r}^{(\beta)} + \sum_{i=1}^{\beta-1}\big(\prod_{j=1}^{i}\tilde{k}_{\beta-j}\big)\tilde{r}^{(\beta-i)} + \big(\prod_{j=1}^{\beta-1}\tilde{k}_{\beta-j}\big)\tilde{k}_N a_N)$
and $\tilde{k}_\beta = \frac{k_{\beta+1}+\rho_N k_N}{\delta_{\beta+1}+k_{\beta+1}+\rho_N k_N} < 1.0$

### 8 Calculation of $a^*_{(\beta+1)}/a^*_\beta$ :

Similarly, it can be shown that for $2 \leq \beta \leq (N-1)$ we get,

$$\boxed{\frac{a^{*(s)}_{(\beta+1)}}{a^{*(s)}_{\beta}} = \frac{\tilde{r}_p^{(\beta+1)}a^{(s)}_{\beta+1}}{a^{*(s)}_{\beta}} + \tilde{k}^{(p)}_\beta} \tag{70}$$

$$\boxed{\frac{a^{*(s)}_{(\beta+1)}}{a^{*(s)}_{(\beta)}} = \frac{\tilde{r}_p^{(\beta+1)}a^{(s)}_{\beta+1}}{\tilde{r}_p^{(\beta)}a_\beta + \sum_{i=1}^{\beta-1}\Big(\prod_{j=1}^{i}\tilde{k}^{(p)}_{\beta-j}\Big)\tilde{r}_p^{(\beta-i)}a_{(\beta-i)} + \Big(\prod_{j=1}^{\beta-1}\tilde{k}^{(p)}_{\beta-j}\Big)\tilde{k}^{(p)}_N a^*_N} + \tilde{k}^{(p)}_\beta} \tag{71}$$

Where, $a^{(s)}_{\beta} = \tilde{r}_p^{(\beta)}a_\beta + \sum_{i=1}^{\beta-1}\Big(\prod_{j=1}^{i}\tilde{k}^{(p)}_{\beta-j}\Big)\tilde{r}_p^{(\beta-i)}a_{(\beta-i)} + \Big(\prod_{j=1}^{\beta-1}\tilde{k}^{(p)}_{\beta-j}\Big)\tilde{k}^{(p)}_N a^*_N$
and $\tilde{k}^{(p)}_\beta = \frac{k_{\beta+1}+\rho_N k_N}{\lambda_{\beta+1}+k_{\beta+1}+\rho_N k_N}$, where $\lambda^{(\beta+1)} = r_p^{(\beta+1)} + r_d^{(\beta+1)} + \delta^{(\beta+1)}$

#### 9 Minimal model for protein synthesis-degradation dynamics

Let us assume a protein $A$ is synthesized at a rate $r$ and degraded at a rate $\delta$ with the zeroth ($\phi \rightarrow A$; rate $r$)and first order ($A \rightarrow \phi$; rate $\delta$) reactions, respectively. The following equation gives the dynamics of concentration of protein $A$

$$\begin{aligned} \frac{dA(t)}{dt} &= r - \delta A \\ A(t) = \frac{1}{\delta}\left[r - (r - \delta A_0)\exp^{(-\delta t)}\right] &= \frac{r}{\delta}[1 - \exp(-\delta t)] + A_0 \exp(-\delta t). \end{aligned} \tag{72}$$

Here, $A_0$ is the initial concentration of the protein at time $t = 0$. At the steady-state limit ($t \rightarrow \infty$), the concentration of the protein ($A^s$) is determined by the ratio of synthesis rate and degradation rate as follows

$$\boxed{A^s = \frac{r}{\delta}} \tag{73}$$

#### 10 Reaction rates for the zeroth-, first- and second-order reaction within a small simulation volume $V_\epsilon$

Let us assume the protein concentration and zeroth-order reaction rate are represented by $\rho_A$ and $k_0$, respectively. $N_A$ and $n_A$ are abundances of proteins within a total cytoplasmic volume $V_{cyto}$ and a small volume $V_\epsilon$ used in the simulation, respectively.

##### Zeroth-order reaction:

The zeroth-order equation ($\phi \rightarrow A$) with reaction rate $k_0(\mu M s^{-1})$ is followed

$$\begin{aligned} \frac{d\rho_A}{dt} &= k_0 \\ \frac{d(N_A)}{dt} &= k_0 V_{cyto} = \tilde{k}_0 \end{aligned} \tag{74}$$

The zeroth-order equation for $n_A$ is

$$\boxed{\frac{d(n_A)}{dt} = \frac{d(N_A * V_\epsilon / V_{cyto})}{dt} = \left(\frac{V_\epsilon}{V_{cyto}}\right)\tilde{k}_0} \tag{75}$$

Thus, the reaction rate $\tilde{k}_0$ (copy $s^{-1}$) is scaled by a multiplicative factor of $\left(\frac{V_\epsilon}{V_{cyto}}\right)$.

##### First-order reaction:

The first-order equation ($A \rightarrow B$) with a reaction rate $k_1(s^{-1})$ is followed:

$$\begin{aligned} \frac{d\rho_B}{dt} &= k_1 \rho_A \\ \frac{d(N_B)}{dt} &= k_1 N_A \\ &\boxed{\frac{d(n_B)}{dt} = k_1 n_A} \end{aligned} \tag{76}$$

Because the above equation is volume independent, the first-order reaction rate $k_1(s^{-1})$ remains constant in any volume.

##### Second-order reaction:

The second-order equation $(A + A \rightarrow B)$ with a reaction rate $k_2(\mu M^{-1}s^{-1})$ is followed:

$$\begin{aligned} \frac{d\rho_B}{dt} &= k_2\rho_A\rho_A \\ \frac{d(n_B/V_\epsilon)}{dt} &= k_2(n_A/V_\epsilon)(n_A/V_\epsilon) \\ \boxed{\frac{d(n_B)}{dt}} &\boxed{= \left(\frac{k_2}{V_\epsilon}\right)n_An_A} \end{aligned} \tag{77}$$

Thus, the second order reaction rate $k_2(\mu M^{-1}s^{-1})$ is scaled by a factor $\left(\frac{1}{V_\epsilon}\right)$.

#### 11 Minimal non-linear signaling model

We consider a minimal non-linear model of protein phosphorylation to investigate the effect of increasing cell volume on the average abundance of the proteins through cell cycle stages. The reaction rules for the signaling network are following.

$$\begin{aligned} \phi \rightarrow E &: \text{ rate } r_E \\ \phi \rightarrow A &: \text{ rate } r_A \\ \text{E, A} \rightarrow \phi &: \text{ rate } \delta \\ \text{E+A} \rightarrow C &: \text{ rate } k_{on} \\ \text{C} \rightarrow E + A &: \text{ rate } k_{off} \\ \text{C} \rightarrow E + P &: \text{ rate } r_p \\ \text{P} \rightarrow A &: \text{ rate } r_d \\ \text{C} \rightarrow \phi &: \text{ rate } \delta_C \\ \text{P} \rightarrow \phi &: \text{ rate } \delta_P \end{aligned} \tag{78}$$

For simulation, we considered a small fraction of volume ($V_\epsilon$) within the cell and simulated the signaling reactions within that region. Because the above-mentioned signaling reactions(Eq. (78)) occur in the cytosolic volume of the cell ($V_{cyto}$), we rescaled the rates of the reactions for our simulation within a small volume $V_\epsilon$ (Sec. 10). Note that, the reaction rate for the zeroth-order ($r_E, r_A$) in units copy per sec reaction will be scaled by a multiplicative factor of ($V_\epsilon/V_{cyto}$). The reaction rate ($M^{-1}s^{-1}$) for the second-order reaction ($k_{on}$) will be changed by a multiplicative factor ($1/V_\epsilon$). The rates for the first-order reactions ($s^{-1}$) will be intact and not dependent on volume. For a typical Natural Killer cell, we consider the cytoplasmic volume to be $V_{cyto} = 4/3 * \pi * (R_{cell}^3 - R_{nu}^3) = 79.59\mu m^3$, where $R_{cell} = 3\ \mu m$ and $R_{nu} = 2\ \mu m$ represent radii of NK cell and NK cell nucleus, respectively. In our simulation, we initially started with 80 copies ($n_\epsilon$) of Enzymes(E) and Substrate(A) which gives the simulation volume to be $V_\epsilon = n_\epsilon/\rho = 0.132\ \mu m^3$. Here, we assumed the typical concentration ($\rho$) of signaling protein to be $\sim 1\mu M \approx 602$ molecules per $\mu m^3$. The dynamics of the $E, A$, complex $C$ and $P$ are given by the following equations in the situation when $r_E = r_A = r$ and $\delta_C = \delta_P = \delta$,

$$\begin{aligned} \frac{d[E]}{dt} &= r - \delta[E] - k_{on}[E][A] + k_{off}[C] + r_p[C], \\ \frac{d[A]}{dt} &= r - \delta[A] - k_{on}[E][A] + k_{off}[C] + r_d[P], \\ \frac{d[C]}{dt} &= k_{on}[E][A] - k_{off}[C] - \delta[C] - r_p[C], \\ \frac{d[P]}{dt} &= r_p[C] - \delta[P] - r_d[P]. \end{aligned} \tag{79}$$

The above equations give the total enzyme and substrate concentration as

$$\frac{d[E_{tot}]}{dt} = r - \delta[E_{tot}]$$
$$\frac{d[A_{tot}]}{dt} = r - \delta[A_{tot}] \tag{80}$$

Where, $E_{tot}$ and $A_{tot}$ are time-dependent total concentrations of the enzyme and substrate, respectively given by $E_{tot} = [E] + [C]$ and $A_{tot} = [A] + [C] + [P]$. In the steady state limit, the concentrations of $E^s_{tot}$ and $A^s_{tot}$ will be equal and constant $=r/\delta$ following Eqn. (72-73). $[E]$ and $[A]$ denote the concentrations of free enzyme and substrate.

At the steady state limit, the concentration of complex C remains constant over time,

$$\frac{d[C]_s}{dt} = 0 = k_{on}[E]_s[A]_s - k_{off}[C]_s - \delta[C]_s - r_p[C]_s$$
$$k_{on}(E^s_{tot} - [C]_s)(A^s_{tot} - [C]_s - [P]_s) = k_{off}[C]_s + \delta[C]_s + r_p[C]_s \tag{81}$$

At steady-state the concentration of phosphoprotein or product P remains fixed with time,

$$\frac{d[P]_s}{dt} = 0 = r_p[C]_s - \delta[P]_s - r_d[P]_s$$
$$[P]_s = \frac{r_p}{(\delta + r_d)}[C]_s = \alpha[C]_s \tag{82}$$

The product $P$ follows the same behavior as complex abundance with a multiplicative factor $\alpha = \frac{r_p}{(\delta+r_d)}$. Substituting $[P]_s$ in Eqn. (83), we get the exact solution of

$$[C]_s = \frac{(A^s_{tot} + \beta E^s_{tot} + K'_D) \pm \sqrt{(A^s_{tot} + \beta E^s_{tot} + K'_D)^2 - 4\beta E^s_{tot} A^s_{tot}}}{2\beta} \tag{83}$$

where $\beta = (1 + \alpha) = \frac{r_p+\delta+r_d}{(\delta+r_d)}$. Where, $K'_D = k'_{off}/k_{on} = (k_{off} + \delta + r_p)/k_{on}$. Because the ratio $\frac{4\beta E^s_{tot} A^s_{tot}}{(A^s_{tot}+\beta E^s_{tot}+K'_D)^2} < 1$, we can expand the solution of Eqn. (83) into Taylor series. Keeping up to first linear term (when $\beta$ or $K'_D$ are large), we get the approximated solution of $[C]_s$ as

$$[C]_s(V) \sim \frac{A^s_{tot} E^s_{tot}}{(A^s_{tot} + \beta E^s_{tot} + K'_D(V)).} \tag{84}$$

Here, the approximated steady-state concentration of complex $[C]_s$ depends on the cell volume.

#### Examples for monotonic and non-monotonic variations in complex ($C$) in steady state of cell cycle transitions

Note that, the expressions of Eq. (84) are calculated only considering signaling reactions between proteins, and no cell cycle transitions have been assumed. Below, we have discussed how cell cycle transition can regulate different protein behaviors across cell cycle stages based on the values of signaling parameters and cell cycle transition rates. Because cell volume increases through the cell cycle stages, protein concentrations decrease and thus effectively reduce the rate of complex (C) formation $k_{on}$ to max by a factor of two. Therefore, in Eq. (84) the limits of $K'_D$ across the cell cycles stages are within $[K'_D(V), 2K'_D(V)]$. The product $[P]_s$ will follow similar behavior to complex $[C]_s$ with a multiplicative factor $\alpha = \frac{r_p}{(\delta+r_d)}$

1. **Monotonically decreasing condition:** This situation arises when the protein degradation rate ($\delta$) is much faster than the cell cycle transition rates ($k_\beta$). In this situation, $A^s_{tot}$ and $E^s_{tot}$ reach the steady state limit, and their abundances becomes constant ($\approx r/\delta$) across cell cycle stages (see Eqn. (72)). As the cell volume increases across cell cycle stages, the function $1/(A^s_{tot} + \beta E^s_{tot} + K'_D(V))$ in Eq. (84) decays in consecutive cell cycle stages. As a result, the phosphoprotein product $P$ (Eq. (82)) shows monotonic decay from $G1$ through $M$ stages. The effect would be more pronounced for higher values of $K'_D$.

2. **Non-monotonic condition:** This situation arises when the protein degradation rates ($\delta$) is much slower than the cell cycle transition rates ($k_\beta$). As a result, $A^s_{tot}$ and $E^s_{tot}$ increase monotonically through cell cycle stages but have not achieved the steady state value ($\approx r/\delta$). However, the steady state of signaling proteins is achieved through many cell cycle transitions and divisions. Because, $1/(A^s_{tot}+\beta E^s_{tot}+K'_D(V))$ is a decaying function with cell volume or cell cycle stages, it competes with the growing function $A^s_{tot} \times E^s_{tot}$. If the decay rate of $1/(A^s_{tot}+\beta E^s_{tot}+K'_D(V))$ is much faster than the rate of growth by $A^s_{tot} \times E^s_{tot}$ at later cell cycle stages (e.g. $G2$ or $M$), we can get non-monotonic rise followed by a decay through $G1$ through $M$.

3. **No change condition:** This situation arises when $A^s_{tot}$ and $E^s_{tot}$ reach to the steady state value ($\approx r/\delta$) in the limit when the protein degradation rate ($\delta$) is much faster than the cell cycle transition rates ($k_\beta$). Additionally, high binding between A and E or the low $K'_D$ can result in no change in the abundances across the cell cycle stages Eq. (84). This situation can also arise if $1/(A^s_{tot}+\beta E^s_{tot}+K'_D(V))$ can counteract the increase in abundances of $A^s_{tot} \times E^s_{tot}$ in all cell cycle stages that occurs where protein degradation rate is comparable to cell cycle transition rates ($k_\beta$).

4. **Monotonically rise condition:** When $A^s_{tot}$ and $E^s_{tot}$ are monotonically increasing through cell cycle stages and but have not achieved the steady state value ($\approx r/\delta$). This situation arises when the protein degradation rates ($\delta$) is much slower than the cell cycle transition rates ($k_\beta$). Even if $1/(A^s_{tot}+\beta E^s_{tot}+K'_D(V))$ is decaying function it cannot compete with the higher growth rate of $A^s_{tot} \times E^s_{tot}$ across cell cycle stages.

#### Calculation of the variation in protein abundances for the non-monotonic *neutral* cases

In the *neutral* scenario, to obtain the maximum variation in phosphoprotein levels due solely to cell volume growth, we used an approximated formula. We first validated our simulation results (presented in **Figure 4**) by comparing them to the approximate formulas in **Supplementary Figure S8**. The phosphoprotein abundances from the simulations showed good agreement with those obtained using the approximate formulas (**Supplementary Text 1**, Section 11, Eqns. 92–94). Next, we applied the approximated formula to calculate the maximum decrease in protein variation for the monotonic decrease case. Using Eqn. (84), we calculated the maximum fold change in phosphoprotein to be $\frac{[C]_s(V)-[C]_s(2V)}{[C]_s(V)} \times 100\% \approx K'_D/(A^s_{tot}+\beta E^s_{tot}+2K'_D) \times 100\%$, which reaches its maximum value $\sim 50\%$ in the limit $\beta \to 1$ and small $(A^s_{tot}+\beta E_{tot})$ (the values are provided in **Supplementary Table S1**). In all other non-monotonic cases in the *neutral* scenario, the variation would be smaller than $\sim 50\%$ or the maximum variation in monotonic decrease case, as the cell volume-dependent propensity cannot counteract the increase in total A ($A_{tot}$) and total E ($E_{tot}$) in every cell cycle stages, which occurs in the monotonic decreasing case.
